# Spatial information transfer in hippocampal place cells depends on trial-to-trial variability, symmetry of place-field firing and biophysical heterogeneities

**DOI:** 10.1101/2020.09.17.301747

**Authors:** Ankit Roy, Rishikesh Narayanan

## Abstract

The relationship between the feature-tuning curve and information transfer profile of individual neurons provides vital insights about neural encoding. However, the relationship between the spatial tuning curve and spatial information transfer of hippocampal place cells remains unexplored. Here, employing a stochastic search procedure spanning thousands of models, we arrived at 127 conductance-based place-cell models that exhibited signature electrophysiological characteristics and sharp spatial tuning, with parametric values that exhibited neither clustering nor strong pairwise correlations. We introduced trial-to-trial variability in responses and computed model tuning curves and information transfer profiles, using stimulus-specific (SSI) and mutual (MI) information metrics, across locations within the place field. We found spatial information transfer to be heterogeneous across models, but to reduce consistently with increasing degrees of variability. Importantly, whereas reliable low-variability responses implied that maximal information transfer occurred at high-slope regions of the tuning curve, increase in variability resulted in maximal transfer occurring at the peak-firing location in a subset of models. Moreover, experience-dependent asymmetry in place-field firing introduced asymmetries in the information transfer computed through MI, but not SSI, and the impact of activity-dependent variability on information transfer was minimal compared to activity-independent variability. Biophysically, we unveiled a many-to-one relationship between different ion channels and information transfer, and demonstrated critical roles for *N*-methyl-D-aspartate receptors, transient potassium and dendritic sodium channels in regulating information transfer. Our results emphasize the need to account for trial-to-trial variability, tuning-curve shape and biological heterogeneities while assessing information transfer, and demonstrate ion-channel degeneracy in the regulation of spatial information transfer.

## INTRODUCTION

Biological organisms rely on information about their surroundings through different senses for survival. They receive, encode and process information about their surroundings in eliciting robust responses to challenges posed by the external environment. From an ethological perspective, it is essential that sensory information is efficiently encoded by neural circuits to ensure effective responses to environmental challenges. A dominant theme of neural circuit organization is the ability of individual neurons to encode specific features associated with the external environment, with different neurons responding maximally to distinct feature values. For instance, neurons in the primary visual cortex respond maximally to a specific visual orientation (Hubel & Wiesel, 1959), neurons in the cochlea respond maximally to specific tones (von Békésy & Wever, 1960) and place cells in the hippocampus act as spatial sensors by responding maximally to specific locations of an animal in its environment (O’Keefe, 1976). Central to this overarching design principle is the concept of tuning curves, whereby neurons that respond maximally to a given feature value also respond to nearby feature values, with the response intensity typically falling sharply with increasing feature distance from the peak-response feature. A fundamental question on neurons endowed with such tuning curves relates to the relationship between the turning curve and the sensory information transfer profile of the neuron across feature values. Although this relationship has been explored in neural responses across different sensory modalities (Bezzi, Samengo, Leutgeb, & Mizumori, 2002; Butts, 2003; Butts & Goldman, 2006; DeWeese & Meister, 1999; Montgomery & Wehr, 2010), the question on the relationship between spatial information transfer and spatial tuning curve within the place field of hippocampal place cells has not been quantitatively assessed.

In this morphologically realistic, conductance-based modeling study, we aim to fill this lacuna on the relationship between spatial tuning curves and spatial information transfer in individual hippocampal place cells *within* their respective place field. In addressing this, we systematically assessed spatial information transfer with reference to the expansive biophysical and physiological heterogeneities associated with hippocampal neurons, to the different forms and levels of trial-to-trial variability in place-cell responses to the same stimuli, and to the emergence of experience-dependent asymmetry in the hippocampal spatial tuning curves. Explicitly, we accounted for electrophysiologically-characterized heterogeneities in ion channel properties, intrinsic response characteristics and synaptic localization, and analyzed the impact of trial-to-trial variability that was either dependent or independent of activity. Instead of hand-tuning a single model and inferring the outcomes from that single model, we built a population of models of the CA1 pyramidal neuron through an unbiased stochastic search spanning 22 different parameters. These parameters defined neuronal passive properties and characteristics of ten distinct electrophysiologically characterized ion channel subtypes from hippocampal neurons. We searched 12,000 stochastic model instances, which were driven by 80 spatially dispersed synapses carrying theta-modulated place-field information. We declared 127 of these models that manifested 22 signature somato-dendritic intrinsic properties and sharp place-field tuning (high firing rate and low spatial width) to be valid models. Although these valid models manifested signature physiological characteristics of CA1 pyramidal neurons, they exhibited significant heterogeneity in the parametric combinations that yielded them (Basak & Narayanan, 2018, 2020; Rathour & Narayanan, 2014, 2019).

We subjected this heterogeneous population of models to different degrees of activity-independent or activity-dependent trial-to-trial variability, introduced into the presynaptic firing profiles. We found that the introduction of activity-independent variability resulted in a progressive increase in neural firing frequency, accompanied by reductions in tuning width and theta-band power, with increasing degree of variability. We assessed spatial information transfer using two sets of information transfer metrics based on mutual information (MI) and stimulus-specific information (SSI). When these place cells acted as reliable sensors of spatial location with low trial-to-trial variability, the neural rate code conveyed maximal spatial information (MI or SSI) at the high-slope, and not the peak-firing, locations of the tuning curve. However, owing to differences in parametric heterogeneity in this population of models, there was significant heterogeneity in spatial information transfer across models although they received identical distributions of afferent synaptic patterns. The heterogeneity manifested in terms of the quantitative value of the amount of information transferred, and in terms of how they responded to increases in the degree of trial-to-trial variability. Specifically, with increases in trial-to-trial variability, whereas one subpopulation of models switched to transferring peak stimulus-specific spatial information at the peak-firing locations, another subpopulation continued to transfer peak information at the high-slope locations. We demonstrated the dependence of the spatial information transfer profile on the type of trial-to-trial variability, whereby activity-dependent variability had little impact on spatial information transfer compared to the significant reduction introduced by activity-independent variability. Importantly, we show that the model population manifested parametric degeneracy, whereby models with similar intrinsic measurements, similar tuning curves and similar information transfer metrics exhibited immense heterogeneity in and weak pair-wise correlations across underlying parameters.

To further delineate the relationship of spatial information transfer with place-cell characteristics and its components, we assessed the impact of experience-dependent asymmetry in the place-field firing rate profile. We found that the MI profile showed a dependence on the asymmetric nature of the firing profile, but the peak values of SSI profile were largely invariant to the asymmetry. Finally, we found heterogeneity in the impact of knocking out individual ion channels on these information metrics, pointing to a many-to-one relationship between different ion channel subtypes and spatial information transfer. As direct experimentally testable predictions, our analyses unveiled a potent reduction in information transfer consequent to virtually knocking out transient potassium channels, NMDA receptors or dendritic sodium channels.

Together, our analyses emphasize the need to account for neuronal heterogeneities in assessing sensory information transfer and show that synergistic interactions among several neural components regulate the specifics of information transfer and its relationship to tuning curve characteristics. These neural components include the ubiquitous biophysical and physiological heterogeneities, the type and degree of trial-to-trial variability, the specific metrics employed to assess information transfer and behavior-dependent alterations to the sensory tuning curves.

## METHODS

The computational modelling of the place cell was performed by employing a morphologically realistic CA1 pyramidal neuron of rat hippocampus. A morphologically reconstructed model (n123; Fig. 1A) was obtained from Neuromorpho.org (Ascoli, Donohue, & Halavi, 2007). Several active and passive mechanisms were incorporated into the model to mimic intrinsic functional properties of a CA1 pyramidal neuron. The passive properties arising due to the lipid bilayer was modelled as a capacitive current, and to represent the leak channels a resistive current was included. The three parameters which regulated the passive electrical properties of the neuron are axial resistivity (*R*_*a*_), specific membrane resistivity (*R*_*m*_) and specific membrane capacitance (*C*_*m*_). In the base model, *R*_*a*_ was set to 120 Ω.cm and the specific membrane capacitance was set to 1 μF/cm^2^ for the entire neuron (Table 1, Figure 1B). The specific membrane resistivity was non-uniform and varied in a sigmoidal manner (Basak & Narayanan, 2018; Golding, Mickus, Katz, Kath, & Spruston, 2005; Narayanan & Johnston, 2007; Rathour & Narayanan, 2014) as a function of the distance of the point from the soma (*x*) (Figure 1B):

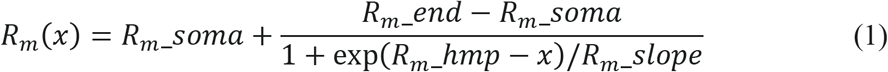

**Table 1:** Model parameters, their base values and ranges for stochastic search. For all parameters, the range uniformly spanned 0.5–2× of the respective base model value.

|  | Parameter (unit) | Symbol | Base value | Range |
| --- | --- | --- | --- | --- |
| **Passive properties** |  |  |  |  |
| **$R_a$ (uniform across the neuron)** |  |  |  |  |
| 1 | Axial resistivity ($\Omega\text{-cm}$) | $R_a$ | 120 | 100–250 |
| **$R_m$ (sigmoidal reduction with distance from soma)** |  |  |  |  |
| 2 | Maximum value ($\text{k}\Omega\text{ cm}^{-2}$) | $R_{m\_soma}$ | 125 | 62.5–250 |
| 3 | Minimum value ($\text{k}\Omega\text{ cm}^{-2}$) | $R_{m\_end}$ | 85 | 42.5–170 |
| 4 | Half-maximal point of sigmoid ($\mu\text{m}$) | $R_{m\_hmp}$ | 300 | 150–600 |
| 5 | Slope of sigmoid ($\mu\text{m}$) | $R_{m\_slope}$ | 50 | 25–100 |
| **Active properties** |  |  |  |  |
| **Spike-generating channels (uniform across all somatodendritic compartments)** |  |  |  |  |
| 6 | Maximum conductance for NaF ($\text{mS cm}^{-2}$) | $\bar{g}_{Na}$ | 16 | 8–32 |
| 7 | Maximum conductance for KDR ($\text{mS cm}^{-2}$) | $\bar{g}_{KDR}$ | 10 | 5–20 |
| **HCN channel (sigmoidal increase with distance from soma)** |  |  |  |  |
| 8 | Maximum somatic conductance ($\mu\text{S cm}^{-2}$) | $\bar{g}_{h\_soma}$ | 25 | 12.5–50 |
| 9 | Fold increase | $\bar{g}_{h\_fold}$ | 12 | 6–24 |
| 10 | Half-maximal point of sigmoid ($\mu\text{m}$) | $\bar{g}_{h\_hmp}$ | 320 | 160–640 |
| 11 | Slope of sigmoid ($\mu\text{m}$) | $\bar{g}_{h\_slope}$ | 50 | 25–100 |
| **T-type calcium channel (sigmoidal increase with distance from soma)** |  |  |  |  |
| 12 | Maximum somatic conductance ($\mu\text{S cm}^{-2}$) | $\bar{g}_{CaT\_soma}$ | 80 | 40–160 |
| 13 | Fold increase | $\bar{g}_{CaT\_fold}$ | 30 | 15–60 |
| 14 | Half-maximal point of sigmoid ($\mu\text{m}$) | $\bar{g}_{CaT\_hmp}$ | 350 | 175–700 |
| 15 | Slope of sigmoid ($\mu\text{m}$) | $\bar{g}_{CaT\_slope}$ | 50 | 25–100 |
| **A-type potassium channel (linear increase with distance from soma)** |  |  |  |  |
| 16 | Maximum somatic conductance ($\text{mS cm}^{-2}$) | $\bar{g}_{KA\_soma}$ | 3.1 | 1.55–6.2 |
| 17 | Fold increase per 100$\mu\text{m}$ | $\bar{g}_{KA\_fold}$ | 8 | 4–16 |
| **N-type calcium channel (till 340 $\mu\text{m}$ from soma in apical dendrites)** |  |  |  |  |
| 18 | Maximum conductance | $\bar{g}_{CaN}$ | 15 | 7.5–30 |
| **R-type calcium channel (dendritic localization)** |  |  |  |  |
| 19 | Maximum conductance in dendrites ($\mu\text{S cm}^{-2}$) | $\bar{g}_{CaR}$ | 15 | 7.5–30 |
| **L-type calcium channel (perisomatic, till 50 $\mu\text{m}$ from soma in apical dendrites)** |  |  |  |  |
| 20 | Maximum conductance ($\text{mS cm}^{-2}$) | $\bar{g}_{CaL}$ | 1.20 | 0.6–2.4 |
| **Small-conductance calcium-activated potassium channel (dendritic localization)** |  |  |  |  |
| 21 | Maximum conductance of SK ($\mu\text{S cm}^{-2}$) | $\bar{g}_{SK}$ | 1.5 | 0.75–3 |
| **M-type potassium channel (perisomatic, till 50 $\mu\text{m}$ from soma)** |  |  |  |  |
| 22 | Maximum conductance ($\mu\text{S cm}^{-2}$) | $\bar{g}_{KM}$ | 1 | 0.5–2 |

**Figure 1.**
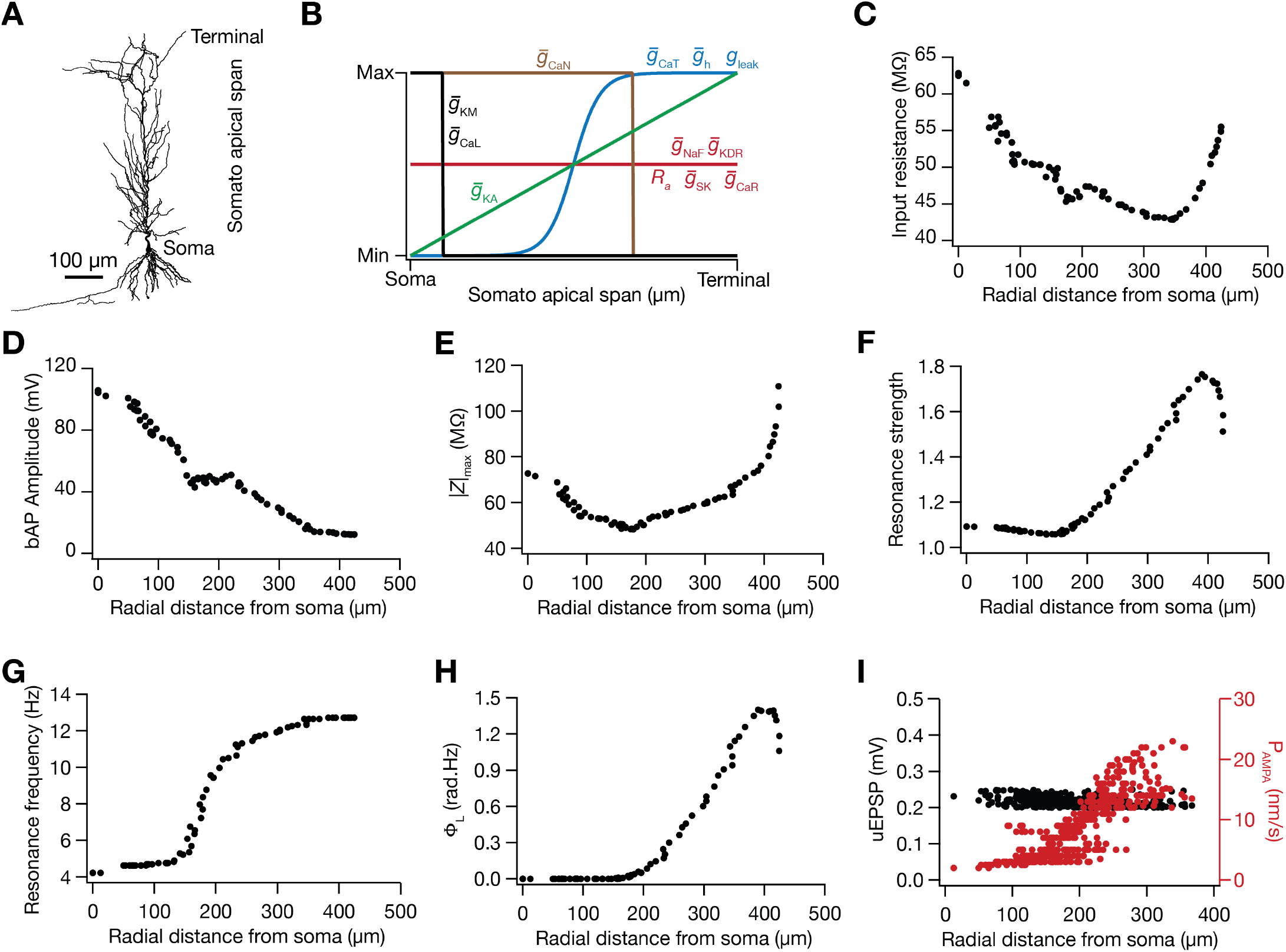
Base model of rat hippocampal CA1 pyramidal neurons, showing its intrinsic and synaptic properties along the somato-apical trunk. (A) Two-dimensional reconstruction of the 3D morphologically realistic model employed in this study. (B) Distribution of parameters governing the passive properties (*g*_*leak*_ and *R*_*a*_) and ten different active ion channels (*g*_*h*_, *g*_*NaF*_, *g*_*KDR*_, *g*_*KA*_, *g*_*KM*_, *g*_*SK*_, *g*_*CaN*_, *g*_*CaL*_, *g*_*CaR*_ and *g*_*CaT*_) along the somato-apical span to match multiple intrinsic measurements at the soma and along the apical dendrites, including input resistance (C), backpropagating action potential amplitude (D) maximum impedance amplitude (E), strength of resonance (F) resonance frequency (G), total inductive phase (H) and the maximum AMPAR permeability (I), all as functions of radial distance from the soma. The distance-dependent profile of maximum AMPAR permeability, *P*_AMPA_ (*I*, right vertical axis) was set such that the somatic unitary excitatory postsynaptic potentials (uEPSPs) was around 0.2 mV, irrespective of synaptic location (*I*, left vertical axis).

In equation 1, *x* is the radial distance from soma, and the parameters and their base-model values are provided in Table 1. The neuron was compartmentalised using the *d*_λ_ rule (Carnevale and Hines, 2006), such that the length of each compartment was lesser than one-tenth of λ_100_, the space constant at 100 Hz. In the base model, this resulted in the compartmentalization of the neuron into 879 distinct compartments.

To model the active properties of the neuron, 10 different types of ion channels were incorporated into the base model, based on electrophysiological characterization from CA1 pyramidal neurons. The ion channels incorporated were the fast sodium (NaF), delayed rectifier potassium (KDR), *A*-type potassium (KA), *M*-type potassium (KM), small-conductance calcium activated potassium (SK), *T*-type calcium (CaT), *N*-type calcium (CaN), *R*-type calcium (CaR), *L*-type calcium (CaL) and hyperpolarisation activated cyclic nucleotide gated (HCN or *h*). The current through these channels due to Na^+^, K^+^ ions were modelled in an ohmic formulation with the reversal potentials of Na^+^, K^+^ and *h* channels being 55, –90 and –30 mV respectively. The current due to calcium ions was modeled as per the Goldman-Hodgkin-Katz (GHK) conventions with the internal calcium concentration as 50 nM and external calcium concentration as 2 mM. The equations underlying the kinetics of these channels were obtained from prior electrophysiological recordings: NaF, KDR and KA (Hoffman, Magee, Colbert, & Johnston, 1997; Magee & Johnston, 1995; M. Migliore, Hoffman, Magee, & Johnston, 1999), HCN (Magee, 1998), KM (Shah, Migliore, Valencia, Cooper, & Brown, 2008), SK (Sah & Clements, 1999; Sah & Isaacson, 1995), CaT (Shah, Migliore, & Brown, 2011), CaN (M. Migliore, Cook, Jaffe, Turner, & Johnston, 1995), CaR and CaL (Magee & Johnston, 1995; Poirazi, Brannon, & Mel, 2003).

These ion channels were distributed along the somatodendritic axis to match experimental recordings (Table 1 provides the distributions and the parameter values in the base model). Specifically, the fast sodium and the delayed rectifier potassium were uniformly distributed (Bittner, Andrasfalvy, & Magee, 2012; Hoffman, et al., 1997; Magee & Johnston, 1995). The *A*-type potassium channel density increased linearly (Hoffman et al. 1997) as a function of distance from soma, *x* (Fig. 1B):

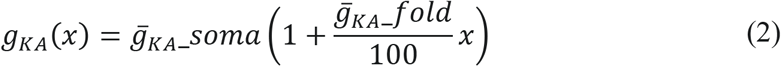

The HCN and *T*-type calcium channel density were set in a sigmoidal manner (Fig. 1B), increasing with radial distance from the soma (Lorincz, Notomi, Tamas, Shigemoto, & Nusser, 2002; Magee, 1998; Magee & Johnston, 1995; Narayanan & Johnston, 2007; Rathour & Narayanan, 2014):

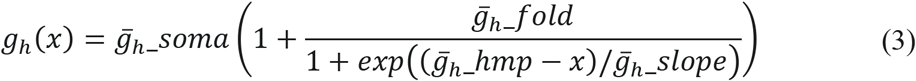

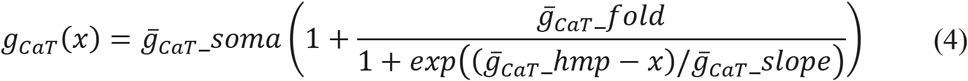

The *M*-type potassium and *L*-type calcium channels were perisomatic (Hu, Vervaeke, & Storm, 2007; Magee & Johnston, 1995). The SK and the *R*-type calcium channels were distributed uniformly across the apical dendrites (Lin, Lujan, Watanabe, Adelman, & Maylie, 2008; Magee & Johnston, 1995; Ngo-Anh, et al., 2005). The *N*-type calcium channels were uniformly distributed till 340 μm of radial distance along the apical dendrite (Magee & Johnston, 1995).

### Intrinsic physiological measurements

We matched several physiological measurements of CA1 pyramidal neurons in our model: back propagating action potential (bAP) amplitude, input resistance (*R*_in_) and impedance-based measurements like resonance frequency (*f*_R_), maximum resonance amplitude (|*Z*|_max_), strength of resonance (*Q*) and total inductive phase (Φ_L_) at different somato-dendritic locations were used to tune the base model and validate models generated by the stochastic search process (Hoffman, et al., 1997; Narayanan, Dougherty, & Johnston, 2010; Narayanan & Johnston, 2007, 2008; Rathour & Narayanan, 2014; Spruston, Schiller, Stuart, & Sakmann, 1995).

To measure input resistance of a somato-dendritic compartment, a hyperpolarizing current step of 100 pA was injected for 500 ms into the compartment. The local change in the membrane potential as a result of the step current was measured and the ratio of the local voltage deflection to the step current amplitude was taken to be the input resistance (Fig. 1C). For measuring the bAP amplitude, a step current of 1 nA was given at the soma for 2 ms. This generated a single action potential at the soma which actively back propagated along the dendrites. The amplitude of the bAP was measured at different locations along the somato-apical trunk (Fig. 1D).

To quantify the frequency dependence of neuronal responses, we employed impedance based physiological measurements across the somato-dendritic arbor (Basak & Narayanan, 2018, 2020; Narayanan, et al., 2010; Narayanan & Johnston, 2007, 2008; Rathour & Narayanan, 2014): resonance frequency (*f*_R_), maximum impedance amplitude (|*Z*|_max_), strength of resonance (*Q*) and total inductive phase (Φ_L_). To measure these a chirp stimulus, defined as a current stimulus with constant amplitude (peak to peak 100 pA) and linearly increasing frequency with time (0–15 Hz in 15 s), was injected in the compartment where the measurement was required. The local voltage response was recorded. To compute the impedance as a function of frequency, the Fourier spectrum of voltage response was divided with the Fourier spectrum of the current giving us the impedance profile as a complex quantity. The magnitude of impedance as a function of frequency was calculated using the following equation,

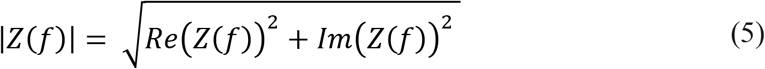

In equation 5, Re(*Z*(*f*)) is the real part of the impedance profile and Im (*Z*(*f*)) is the imaginary part of the impedance profile and |*Z*(*f*)| is the magnitude of impedance. The maximum impedance amplitude was measured and the frequency at which it occurred was taken to be the resonance frequency. The strength of resonance was measured by taking ratio of the maximum impedance amplitude to the impedance amplitude at 0.5 Hz. For the phase related measures, the impedance phase profile was computed:

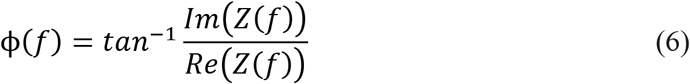

In equation 6, ϕ(*f*) is the phase as a function of frequency. The total inductive phase was measured by calculating the area under the positive portion of phase profile:

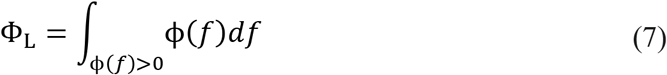

### Synapses and normalization of somatic unitary synaptic potential

The model contained excitatory synapses with colocalized NMDAR and AMPAR, with an NMDAR-to-AMPAR ratio of 1.5, with 80 such synapses randomly dispersed across the apical dendritic arbor (Basak & Narayanan, 2018, 2020). The current through the NMDAR were divided into current due to three ions, Na^+^, K^+^ and Ca^2+^. The dependence of current due to each of these ions as a function of voltage and time was modelled in GHK formulation (Anirudhan & Narayanan, 2015; Ashhad & Narayanan, 2013; Basak & Narayanan, 2018, 2020; Narayanan & Johnston, 2010):

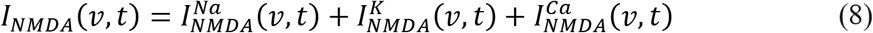

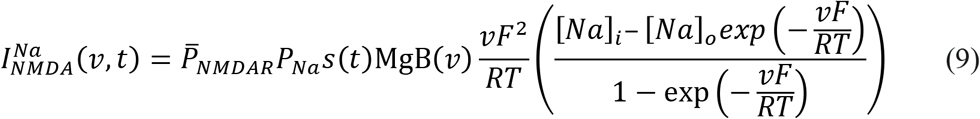

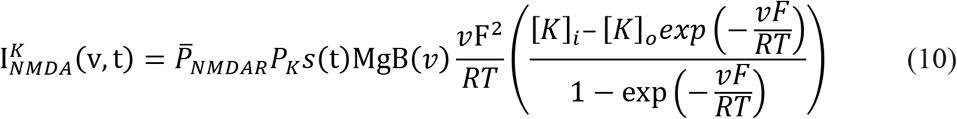

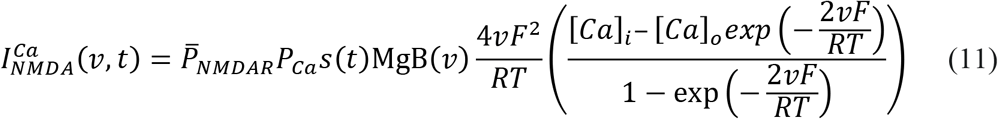

Here, 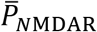 defined the maximum permeability of NMDA receptors. The relative permeability ratios were set to *P*_*Ca*_ = 10.6, *P*_*Na*_ = 1 and *P*_K_ = 1. The ionic concentrations were set as, [*Na*]_*i*_ = 18 mM, [*Na*]_*o*_ = 140 mM, [*K*]_*i*_ = 140 mM, [*K*]_*o*_ = 5 mM, [*Ca*]_*i*_ = 100 nM and [*Ca*]_*o*_ = 2 mM. The magnesium dependence of the NMDAR current was calculated as follows (Jahr & Stevens, 1990):

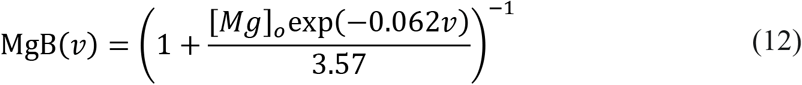

with [*Mg*]_*o*_ = 2 mM. The kinetics of the NMDAR current was determined by *s*(*t*) :

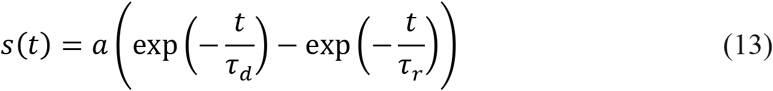

Here *a* is a normalisation constant such that 0 ≤ *s*(*t*) ≤ 1, *τ*_*d*_ is the decay constant, *τ*_*r*_ is the rise time, with *τ*_*r*_ = 5 ms and default *τ*_*d*_ = 50 ms (Narayanan and Johnston, 2010; Ashhad and Narayanan, 2013).

The current through the AMPA receptor was mediated by two ions, Na^+^ and K^+^.

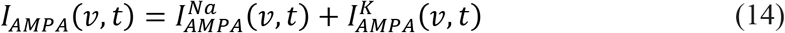

In equation 14,

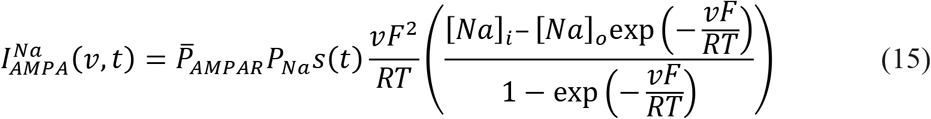

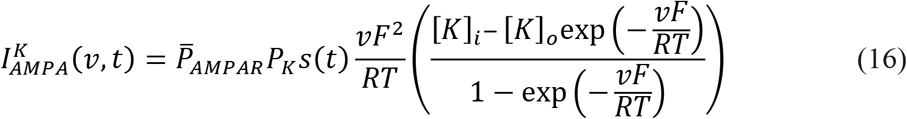

In equations 15–16, 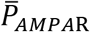 defined the maximum permeability of AMPA receptors. The relative permeability ratios were set to *P*_*Na*_ = 1 and *P*_*K*_ = 1. The *s*(*t*) was modelled in a manner similar to NMDAR with τ_*r*_ = 2 ms and τ_*d*_ = 10 ms. To normalize the unitary EPSP values associated with each synapse, we ensured that the attenuation along the dendritic cable did not affect the unitary somatic EPSP amplitude. Hence, the AMPAR permeabilities at the somato-apical trunk was tuned such that it produced a unitary somatic response of ~0.2 mV irrespective of the synaptic location (Andrasfalvy & Magee, 2001; Magee & Cook, 2000).

### Place cell inputs and synaptic localization

The input to this neuron was fed through colocalized AMPAR-NMDAR synapses. As the virtual animal traversed through the place field the presynaptic neurons fired action potentials. Their firing rates were modeled in a stochastic manner, driven by a Gaussian modulated cosinusoidal function, mimicking place cell inputs to the neuron (Basak & Narayanan, 2018, 2020; Seenivasan & Narayanan, 2020). The presynaptic firing drove the opening of the colocalized synaptic NMDAR and AMPARs, resulting in synaptic currents (equations 8–16) flowing into the model neuron. The Gaussian modulated cosinusoidal function that defined the probability of occurrence of a presynaptic spike to each synapse in the neuron was computed as (Basak & Narayanan, 2018, 2020; Seenivasan & Narayanan, 2020):

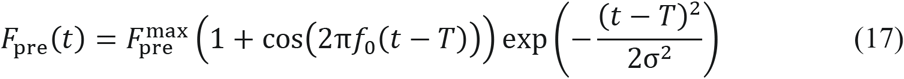

In equation 17, *T* (5 s) defined the center of the place field, *f*_0_ is the frequency of the cosine (8 Hz), 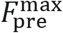 is the maximal input firing rate, σ is the standard deviation of the Gaussian (1 s). In our analyses, the virtual animal was assumed to traverse a linear arena at constant velocity, implying the equivalence of time and space as the independent variable in equation (17). The input current resulting from synaptic activation produced post-synaptic action potentials and caused place cell like firing activities in the model neuron.

In introducing experience-dependent asymmetry in place-field firing (Harvey, Collman, Dombeck, & Tank, 2009; Mehta, Barnes, & McNaughton, 1997; Mehta, Lee, & Wilson, 2002; Mehta, Quirk, & Wilson, 2000), we replaced the symmetric Gaussian profile in equation (17) by a horizontally reflected Erlang distribution to construct an asymmetric place-field envelope (Seenivasan & Narayanan, 2020). In this scenario, the Erlang-modulated cosinusoidal function that defined the probability of occurence of a presynaptic spike to each synapse in the neuron was computed as:

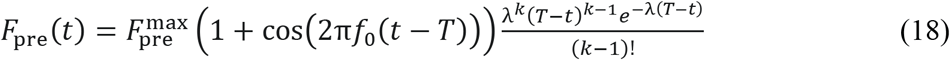

In equation (18), the parameters *λ* (=5) and *k* (=25) governed the extent of asymmetry (Seenivasan & Narayanan, 2020).

### Trial-to-trial variability in place cell responses

For simulating trial-to-trial variability in the place cell firing profile with different degrees of variability, noise was introduced into the presynaptic firing rate profile (equation 17) associated with each synapse. Simulations were performed with Gaussian white noise (GWN) which was introduced either additively (AGWN) or multiplicatively (MGWN):

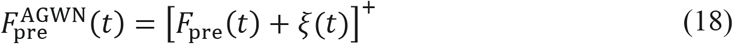

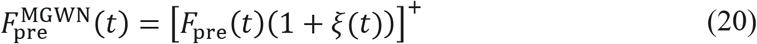

In equations (19–20), [*F*]^+^ = max(*F*, 0) represents rectification to avoid negative firing rates, *ξ*(*t*) defined a GWN with zero mean and standard deviation *σ*_noise_. The value of *σ*_noise_ was increased to enhance the degree of trial-to-trial variability, with *F*_pre_ (*t*) defined by a Gaussian-(equation 17) or an Erlang-envelope (equation 18) to assess the impact of trial-to-trial variability in symmetric or asymmetric place-field firing profiles, respectively. As AGWN (equation 19) introduced trial-to-trial variability across stimulus locations, irrespective of the strength of afferent synaptic activity, this form of variability is *activity-independent*. On the other hand, the degree of trial-to-trial variability introduced by MGWN is progressively higher with increasing strength of afferent synaptic activity (equation 20), thereby manifesting as *activity-dependent* trial-to-trial variability.

### Neuronal voltage response during place-field traversal

Spikes were detected from the place-cell voltage response to afferent synaptic stimuli (equations 17–20) by setting a voltage threshold on the rising phase of the voltage values. These spike timings were employed to compute the firing rate of the place as a function of time (*F*(*t*)) through convolution with a Gaussian kernel (*σ*=200 ms). The maxima (*F*_max_) and the full-width at half maximum (*FWHM*) of the place-cell firing profile were employed as relative measures of place-field tuning sharpness. Specifically, high *F*_max_ and low *FWHM* were indicative of a sharply tuned place cell responses (Basak & Narayanan, 2018, 2020). As animals traverse through the place field of a given hippocampal place cell, these neurons are known to produce characteristic sub-threshold voltage ramps (Harvey, et al., 2009). To assess such ramps, we filtered the voltage traces using a 0.75 s wide median filter, which removed the spikes and exposed the sub-threshold structure of the voltage response during place-field traversal. The maximum value of these ramps was taken as peak ramp voltage (*V*_ramp_). Since the firing rate of the presynaptic neurons were modulated with a sinusoid of theta frequency (8 Hz, equations 17–18), we analyzed whether the post synaptic voltage traces reflected this temporal modulation. The voltage trace at the soma was filtered using a 50 ms wide median filter, to eliminate spikes but retain theta-frequency temporal modulation, and the Fourier spectrum of the filtered signal was computed. The power at 8 Hz of this power spectrum was employed as theta power (Basak & Narayanan, 2018, 2020; Seenivasan & Narayanan, 2020).

### Spatial information transfer within a place field: Mutual information metrics

To quantify the information transmitted through the firing pattern of a place cell, we employed two sets of information metrics. The first set employed the computation of mutual information (MI), with space within the place field considered as the stimulus and the neuronal firing-rate considered the response. The aforementioned equivalence of time and space as the independent variable in equations 17–20 allowed us to compute spatial information transfer from the firing rate response.

Mutual information was computed at 20 different locations (*N*_loc_) from the instantaneous firing-rate profile obtained for 30 different trials. To compute MI at these 20 locations, each location was subdivided into 4 bins, and the associated firing rate response was quantized into 20 bins. Mutual information between the spatial stimulus (*S*) and firing-rate response (*F*) was calculated at each *N*_loc_ as:

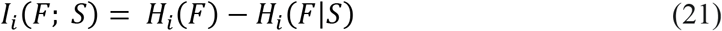

where, *I*_*i*_(*F*;*S*) denoted mutual information between the response and the spatial stimulus at the *i*^th^ location (*i* = 1 … *N*_loc_), and *F* defined the firing rate for *S*. The response entropy *H*_*i*_(*F*) was calculated as:

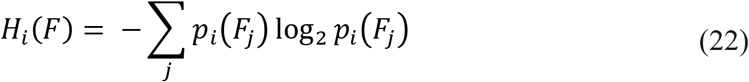

where, *p*_*i*_(*F*_*j*_) represented the probability of the firing rate lying in the *j*^th^ response bin within the *i*^th^ spatial location, and was computed as:

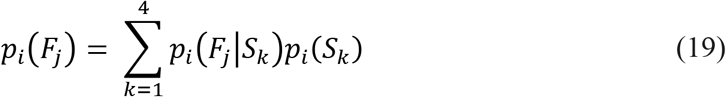

In equation (23), *p*_*i*_(*F*_*j*_|*S*_*k*_) represented the conditional probability that the response was in the *j*^th^ firing rate bin, given that the stimulus was in the *k*^th^ spatial bin within the *i*^th^ spatial location. *p*_*i*_(*S*_*j*_) denoted the probability that the virtual animal was in the *k*^th^ spatial bin within the *i*^th^ spatial location, which was considered to follow a uniform distribution given the constant velocity assumption.

The noise entropy term *H*_*i*_(*F*|*S*) in equation (21) was computed as:

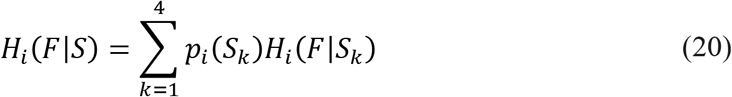

where *H*_*i*_(*F*|*s*_*k*_) represented the conditional noise entropy for the *k*^th^ spatial bin within the *i*^th^
spatial location, calculated as:

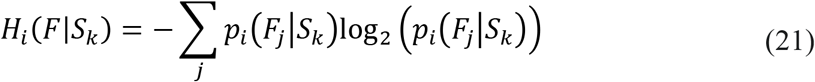

where *p*_*i*_(*F*_*j*_|*S*_*k*_) denoted the conditional probability of the firing rate being in the *j*^th^ bin given that the stimulus was in the *k*^th^ spatial bin within the *i*^th^ location.

### Spatial information transfer within a place field: Stimulus-specific information metrics

The second set of metrics that we employed to compute spatial information transfer was derived from stimulus-specific information (SSI), obtained for 30 different trials of the entire traversal spanning all spatial locations. SSI has been proposed and employed as a measure of information in neuronal response about a particular stimulus, and conveys the average specific information spanning all responses to a particular stimulus. To calculate the SSI the spatial stimulus and the firing rate response were segregated into 80 and 40 bins, respectively. The SSI was calculated using the expression given below (Butts, 2003; Butts & Goldman, 2006; Montgomery & Wehr, 2010):

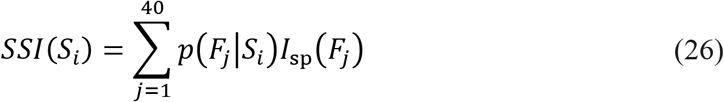

where *p*(*F*_*j*_|*S*_*i*_) is the conditional probability of the firing rate being in the *j*^*th*^ response bin given that the *i*^*th*^ stimulus location was presented, and the specific information *I*_sp_(*F*_*j*_) (DeWeese & Meister, 1999) was computed as:

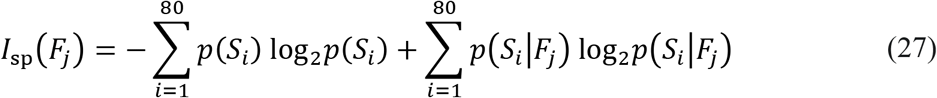

Here, *p*(*F*_*j*_) is the probability of the firing rate being in the *j*^*th*^ response bin and *p*(*S*_*i*_|*F*_*j*_) defined the conditional probability for the stimulus in the *i*^th^ bin given that the firing rate was in the *j*^th^ response bin. The first term in equation (27) represents the entropy of the stimulus ensemble *H*(*S*_*i*_) and the second term represents the entropy of the stimulus distribution conditional on a particular firing rate response *H*(*S*_*i*_/*F*_*j*_), providing *I*_sp_(*F*_*j*_) = *H*(*S*_*i*_) − *H*(*S*_*i*_/*F*_*j*_) (Butts, 2003; Butts & Goldman, 2006; Montgomery & Wehr, 2010). Before employing *I*_sp_(*F*_*j*_) for computing the SSI, bias in *I*_sp_(*F*_*j*_) calculation was corrected using the Treves-Panzeri correction procedure (Bezzi, et al., 2002; Montgomery & Wehr, 2010; Panzeri, Senatore, Montemurro, & Petersen, 2007; Panzeri & Treves, 1996; Treves & Panzeri, 1995):

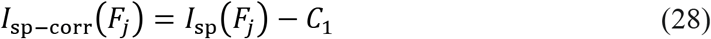

where 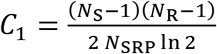 with *N*_S_ representing the total number of stimulus bins, *N*_R_ denoting the total number of response bins and *N*_SRP_ depicting the total number of stimulus-response pairs.

Spatial information transfer as a function of space within a place field was found to be bimodal or trimodal in several scenarios. To quantify the information and compare the information transfer across models and across the different degrees of trial-to-trial variability, several MI-based and SSI-based information metrics were developed and employed (listed in Table 3).

### Exploring parametric dependencies in spatial information transfer

A single hand-tuned model does not account for the numerous biophysical heterogeneities inherent to neural structures, and the results obtained with a single model could be biased by the specific selection of parametric values. A simple methodology to account for the biophysical heterogeneities with signature electrophysiological properties of specific neuronal subtype under consideration is to build a population of models. We employed a multi-parametric multi-objective stochastic search (MPMOSS) algorithm to arrive at a population of models that would satisfy the several biophysical heterogeneities (by allowing the multiple parameters to span an experimental range, shown in Table 1) and would match with bounds on several electrophysiological measurements (Table 2). Since this procedure involves a uniform random sampling of parameter values, it is unbiased and provides a good strategy to search for interdependencies between parametric combinations that yield signature electrophysiological characteristics.

**Table 2:**
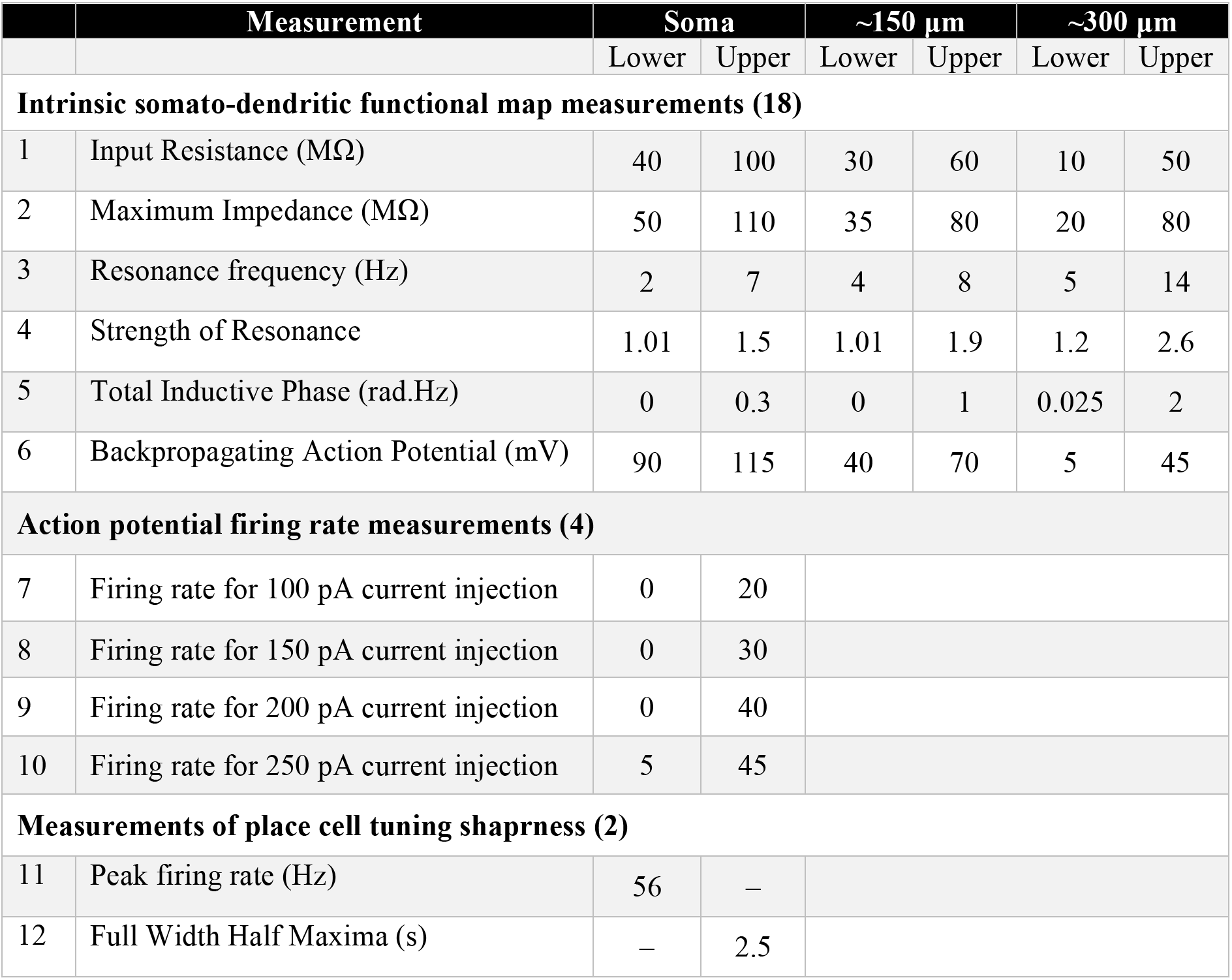
Intrinsic somato-dendritic measurements of CA1 pyramidal neurons and their electrophysiological bounds employed for validating models.

| | Measurement | Soma | | ~150 $\mu\text{m}$ | | ~300 $\mu\text{m}$ | |
| --- | --- | --- | --- | --- | --- | --- | --- |
|  |  | Lower | Upper | Lower | Upper | Lower | Upper |
| Intrinsic somato-dendritic functional map measurements (18) |  |  |  |  |  |  |  |
| 1 | Input Resistance ($\text{M}\Omega$) | 40 | 100 | 30 | 60 | 10 | 50 |
| 2 | Maximum Impedance ($\text{M}\Omega$) | 50 | 110 | 35 | 80 | 20 | 80 |
| 3 | Resonance frequency (Hz) | 2 | 7 | 4 | 8 | 5 | 14 |
| 4 | Strength of Resonance | 1.01 | 1.5 | 1.01 | 1.9 | 1.2 | 2.6 |
| 5 | Total Inductive Phase (rad.Hz) | 0 | 0.3 | 0 | 1 | 0.025 | 2 |
| 6 | Backpropagating Action Potential (mV) | 90 | 115 | 40 | 70 | 5 | 45 |
| Action potential firing rate measurements (4) |  |  |  |  |  |  |  |
| 7 | Firing rate for 100 pA current injection | 0 | 20 |  |  |  |  |
| 8 | Firing rate for 150 pA current injection | 0 | 30 |  |  |  |  |
| 9 | Firing rate for 200 pA current injection | 0 | 40 |  |  |  |  |
| 10 | Firing rate for 250 pA current injection | 5 | 45 |  |  |  |  |
| Measurements of place cell tuning sharpness (2) |  |  |  |  |  |  |  |
| 11 | Peak firing rate (Hz) | 56 | – |  |  |  |  |
| 12 | Full Width Half Maxima (s) | – | 2.5 |  |  |  |  |

To match physiological outcomes, these models were then validated on the basis of sharpness of their place-cell firing properties (*F*_max_ > 56 Hz and *FWHM* < 2.5 s; 2 measurements), six signature intraneuronal functional maps (Basak & Narayanan, 2018, 2020; Narayanan & Johnston, 2012) of back propagating action potential amplitude (*bAP*), input resistance (*R*_in_), resonance frequency (*f*_R_), maximum impedance amplitude (|*Z*|_max_), strength of resonance (*Q*) and total inductive phase (Φ_L_), each validated at three locations (soma, ~150 μm and ~300 μm from soma on the apical trunk; total 18 measurements) and firing rate at the soma resulting from step current injections of 100 pA, 150 pA, 200 pA and 250 pA (4 measurements). Only the models that matched the bounds on these 24 measurements (Table 2) were declared valid. To explore interdependencies among parameters that resulted in the valid models, which showed sharp place-field tuning and manifested signature intrinsic electrophysiological properties, pairwise Pearson’s correlation coefficients spanning the parameters of all valid models were computed. To assess the impact of individual channels in the model on spatial information transfer, we removed each channel individually from the model (by setting the conductance value associated with that channel to zero) and assessed how the information measures changed due to the removal of this ion channel.

### Computational details

All simulations were performed using custom-written software in the NEURON simulation environment (Carnevale & Hines, 2006), at 34° C with an integration time step of 25 μs. Unless otherwise stated, all simulations were performed with a resting potential of –65 mV. Analysis was performed using custom-built software written in Igor Pro programming environment (Wavemetrics). Statistical tests were performed using statistical computing language R (www.R-project.org), and the *p* values are reported while presenting the results, or in the respective figure panels or associated captions. In qualitatively defining weak and strong correlations, we employed the nomenclature followed by (Evans, 1996) by placing thresholds on the absolute value of the Pearson’s correlation coefficient: 0–0.19: Very weak; 0.2–0.39: Weak; 0.4–0.59: Moderate; 0.6–0.79: Strong; 0.8–1: Very Strong. To avoid potential misinterpretations arising from representing data by merely their summary statistics (Marder & Taylor, 2011; Rathour & Narayanan, 2019), all data points from the population of neural models are depicted as beeswarm or scatter plots.

## RESULTS

We built a morphologically realistic, conductance-based model of a CA1 pyramidal cell, incorporating electrophysiologically characterized passive and active mechanisms (Fig. 1A). The model contained 10 distinct biophysically constrained ion channel subtypes that were distributed along the somatodendritic arbor to match experimental findings (Fig. 1B). We hand-tuned the base model parameters (Table 1) to match several intrinsic somato-dendritic electrophysiological properties (Table 2) of rat CA1 pyramidal neurons (Fig. 1C–H). We tuned the strength of synaptic connections such that the somatic unitary AMPAR EPSP was set to ~0.2 mV (Fig. 1I) irrespective of synaptic location within the *stratum radiatum* of the CA1 pyramidal neuron (~350 μm of apical dendrites from the soma).

### Ion-channel degeneracy in the concomitant emergence of sharply tuned spatial firing profile and intrinsic physiological properties of the neuron

As a first step in evaluating the impact of heterogeneous ion channel combinations on sharp tuning of place cell responses, we generated 12,000 random models by independent selection of parameter values from their respective uniform distributions (Table 1). We randomly dispersed 80 distinct synaptic locations (of the 428 possible locations) across the *stratum radiatum* where presynaptic afferent inputs impinged. These 80 synapses received independent presynaptic inputs governed by equation (17), and the somatic voltage response of the neuron was recorded to compute the place-field firing rate profile.

We validated the firing rate profiles of these randomly generated neuronal models for sharpness of place field tuning by placing thresholds on maximum firing rate within the place field (> 56 Hz) and the width of the firing rate profile (<2.5 s), and found 1024 of the 12,000 models (~8.5%) to satisfy these constraints (Fig. S1). We picked five models within these 1024, with similar place-field firing profiles reflected as similar values of *F*_max_ and *FWHM* and asked if similar place-field tuning required similar parametric combinations (Fig. S1A–B). Consistent with prior findings with models endowed with fewer ion channels (Basak & Narayanan, 2018, 2020), we found that sharp-tuning of place field profiles could be achieved with disparate combinations of underlying biophysical parameters, pointing to ion channel degeneracy in the expression of sharp place-field tuning (Fig. S1C). Across all 1024 sharply-tuned models, whose *F*_max_ and *FWHM* are depicted in Fig. S1D–E, the parameters spanned the entire valid range of parameters pointing to the absence of any parametric clustering in arriving at sharp spatial tuning (Fig. S1F). We explored pairwise correlations of the parameters underlying these place-cell models with sharply tuned firing profiles, and found most of the correlation coefficients to be weak (Fig. S1F).

Whereas place-field tuning constitutes one aspect of CA1 pyramidal neuron physiology, their well-characterized signature somato-dendritic intrinsic properties form a defining electrophysiological attribute. To match our model population with these signatures, we validated the 1024 sharply tuned models against 22 distinct electrophysiological measurements (Table 2): each of input resistance, backpropagating action potential amplitude, maximal impedance amplitude, resonance frequency, resonance strength and total inductive phase at 3 different somato-dendritic locations; and action potential firing rate in response to somatic pulse current injections at 4 different current values. Of the total 12,000 models generated, we found 127 (~1.06%) models to match all 24 measurement bounds (Table 2) and were declared valid. We picked five models within these 127 valid models, with similar place-field firing profiles (Fig. S2A) and similar intrinsic measurements across the somato-dendritic axis (Fig. S2B–F). We assessed the parameters associated with five models and found that the concomitant expression of similar place-field tuning and similar intrinsic properties could be achieved with disparate combinations of underlying biophysical parameters, pointing to ion channel degeneracy in the expression of sharp place-field tuning (Fig. S2G). Across all 127 models that were intrinisically-valid (Fig. 2A–G) and sharply-tuned (Fig. 2H–J), the parameters spanned the entire valid range of parameters pointing to the absence of any parametric clustering in these models (Fig. 3). We explored pairwise correlations of the parameters underlying these models, and found most of the correlation coefficients to be weak (Fig. 3).

**Figure 2.**
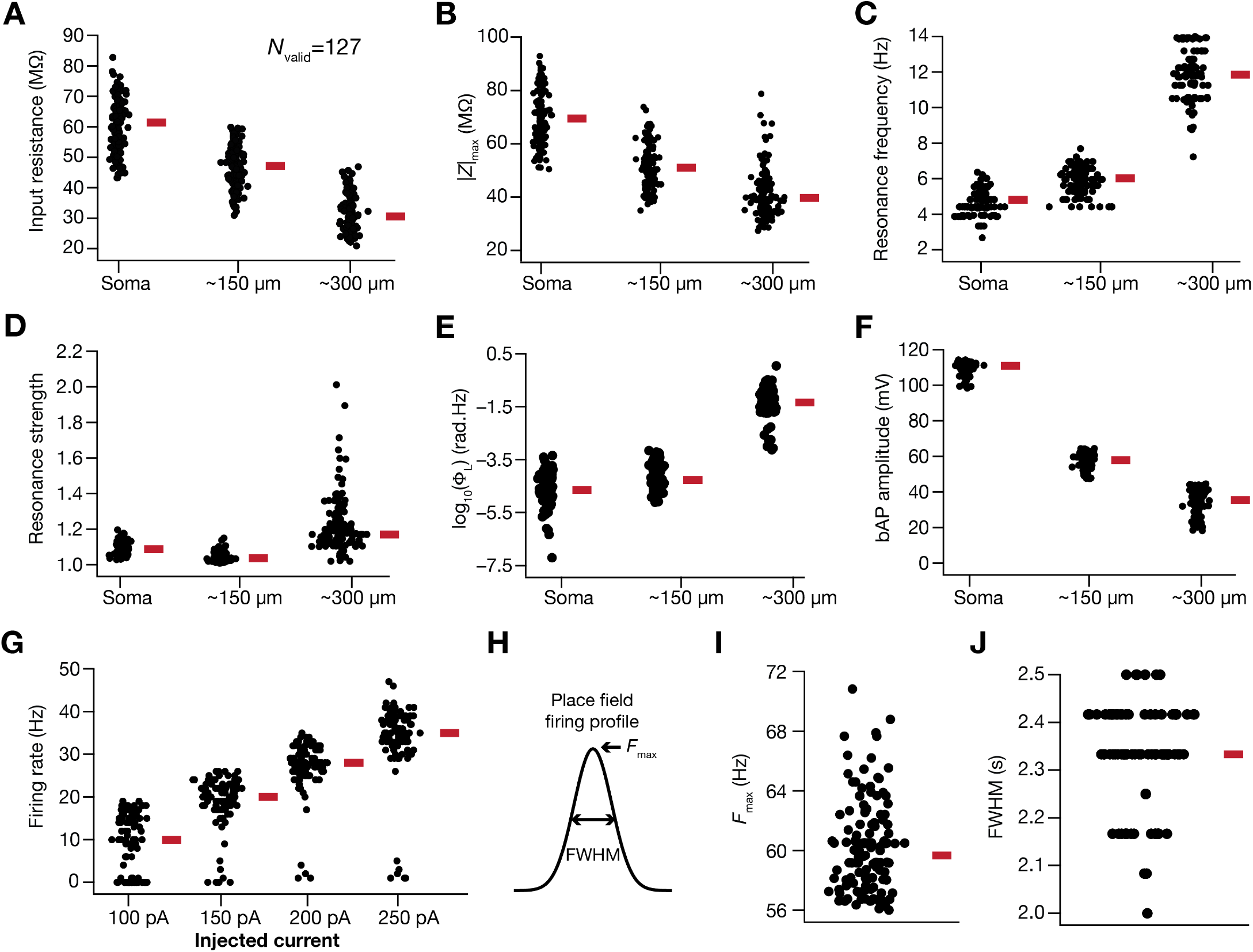
A subset of models generated through a stochastic search process showed sharp place-cell tuning and manifested signature somato-dendritic intrinsic measurements of CA1 pyramidal neurons. Out of 12000 randomly generated models, 127 satisfied 20 intrinsic somato-dendritic measurements and manifested sharply-tuned place field firing. (A–G) The intrinsic measurements for the 127 valid models are shown: input resistance (*R*_in_, A), maximum impedance amplitude (|*Z*|_max_, B), resonating frequency (*f*_R_, C), strength of resonance (*Q*, D), total inductive phase (Φ_L_, E) and back-propagating action potential (bAP) amplitude (F), each of them at three locations (soma, ~150 μm from soma and ~300 μm from soma) on the apical trunk; and the firing rate for step currents of 100 pA, 150 pA, 200 pA and 250 pA at the soma (G). (H) A typical place-field firing profile illustrating the measurement of maximum firing rate (*F*_max_) anad the temporal distance between the places with half the maximum value of firing rate (*FWHM*). A relative criterion on tuning sharpness, involving high *F*_max_ (>56 Hz) and low *FWHM* (<2.5 s), was applied to obtain the 127 valid place-cell models (out of the 12000 randomly generated models). (I–J) Place field firing measurements *F*_max_ and *FWHM* at the soma for the 127 models.

**Figure 3.**
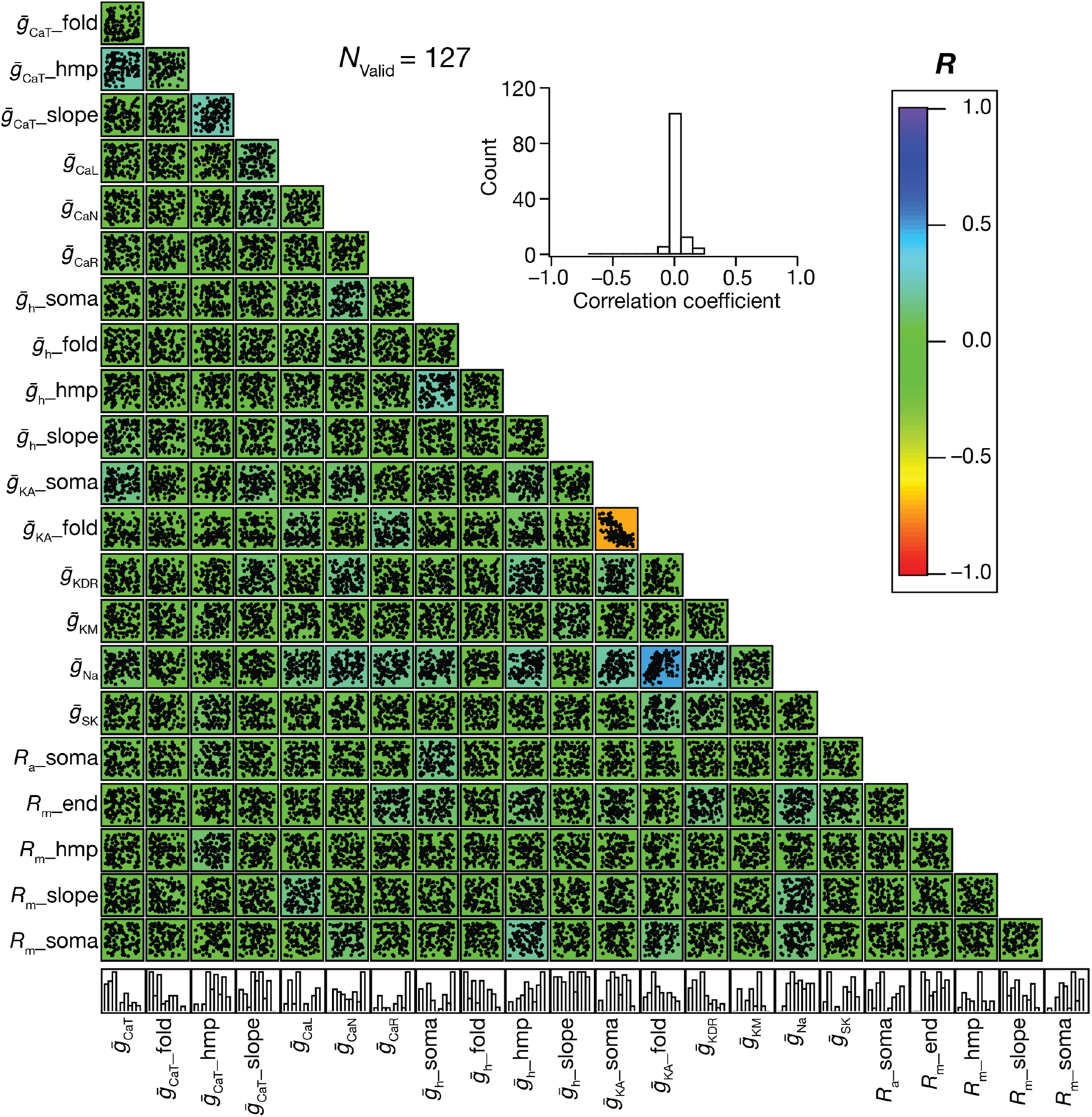
Models showing sharp place-field firing and signature intrinsic characteristics exhibited wide parametric variability and weak pair-wise correlations among underlying parameters. Pairwise scatter plot matrix of parametric values defining the 127 valid models superimposed on the corresponding correlation coefficient matrix. Inset shows the histogram of all the correlation coefficient values.

Together, the unbiased stochastic search procedure provided us with a population of place-cell models that exhibited several signature electrophysiological properties, and manifested sharp place-field tuning in their firing rate profiles. This population of models did not exhibit parametric clustering or strong parametric correlations, pointing to the expression of ion channel degeneracy in achieving these physiological goals. We employed this population of place-cell models for assessing the impact of several biophysical and physiological characteristics on spatial information transfer within the place field.

### Heterogeneities in the regulation of spatial information transfer by trial-to-trial variability in place cell responses

The firing profile of a place cell within its place field represents a spatial tuning curve, with the spatial location at the center of a place-field eliciting the peak firing response and the response progressively falling for spatial stimuli on either side of this peak (*e.g.*, Fig. 4A–B). Within the place field of this neuron, does maximal spatial information transfer occur at the peak of this tuning curve or at the high-slope regions of the tuning curve? Prior studies in other brain regions have shown that the answer to this question depends on several factors, with trial-to-trial variability playing a prominent role in regulating the relationship between the tuning curve and information transfer (Butts & Goldman, 2006; Montgomery & Wehr, 2010). To address this question for spatial information within the place field of individual place cells, we introduced trial-to-trial variability in neural responses by introducing synaptic noise into the afferent input frequency (equation 19).

**Figure 4.**
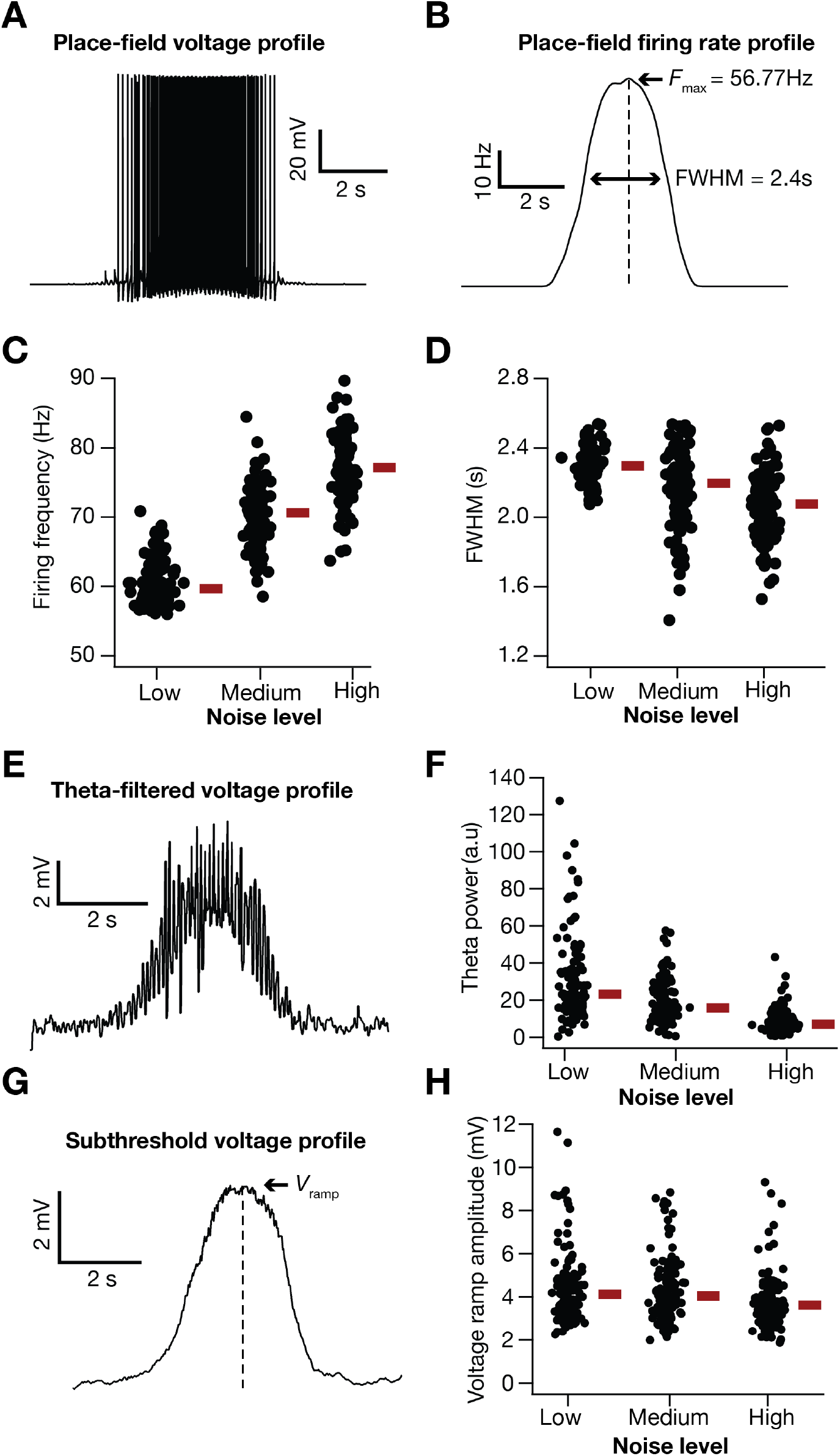
Impact of additive Gaussian white noise (AGWN) on place-cell characteristics. (A–B) Voltage trace (A) and corresponding firing rate profile (B) during traversal of a place field in a typical valid place-cell model in the presence of AGWN (*σ*_noise_=5×10^−4^ Hz^2^). (C–D) Impact of different levels of AGWN on the peak firing frequency, *F*_max_ (C) and full-width at half maximum, *FWHM* (D) of the 127 valid place-cell models. The red bars represent the respective median values. *F*_max_: Kruskal Wallis test, *p*=2.2×10^−12^, Wilcoxon Signed Rank test, Low *vs*. Medium *p*=3.6×10^−8^, Medium *vs.* High *p*=5.3×10^−6^, Low *vs.* High = 3.3×10^−10^. FWHM: Kruskal Wallis test, *p*=8.8×10^−8^, Wilcoxon Signed Rank test, Low *vs*. Medium *p*=4.3×10^−4^, Medium *vs.* High *p*=2.3×10^−4^, Low *vs.* High = 5.3×10^−6^. (E) Voltage profile in (A) filtered to emphasize theta-frequency oscillations during traversal of a place field. (F) Impact of different levels of AGWN on theta power of the 127 valid place-cell models. Kruskal Wallis test, *p*=1.2×10^−8^, Wilcoxon Signed Rank test, Low *vs*. Medium *p*=7.0×10^−4^, Medium *vs.* High *p*=6.1×10^−7^, Low *vs.* High = 5.3×10^−8^. (G) Voltage profile in (A) filtered to emphasize subthreshold voltage ramp during traversal of a place field. (H) Impact of different levels of AGWN on voltage ramp amplitude of the 127 valid place-cell models. Kruskal Wallis test, *p*=2×10^−4^, Wilcoxon Signed Rank test, Low *vs*. Medium *p*=0.4152, Medium *vs.* High *p*=3.4×10^−3^, Low *vs.* High = 7.7×10^−5^. When present, the red bars represent the respective median values. AGWN *σ*_noise_ values: *Low*: 5×10^−4^ Hz^2^, *Medium*: 1×10^−3^ Hz^2^, *High*: 5×10^−3^ Hz^2^.

The introduction of synaptic noise as additive Gaussian white noise (AGWN) manifested as trial-to-trial variability in the firing rate responses, enhanced the firing rate (Fig. 4C) and reduced the width (Fig. 4D) of place cell responses. Across all 127 valid models, progressive increase in trial-to-trial variability, introduced by increasing *σ*_noise_ (equation 19), resulted in a progressive increase in the peak firing rate (Fig. 4C), and progressive reductions in the FWHM (Fig. 4D), theta power (Fig. 4E–F) and the voltage ramp (Fig. 4G–H) of the place-field response profile. We performed 30 trial simulations for each of the 127 valid place-cell models, obtained their firing rate profiles for 3 different levels of noise (Fig. 5A–C; designated as low, medium and high) and computed stimulus-specific information (SSI; Fig. 5D–F) and mutual information (MI; Fig. 5G–I) for all these 127 models.

**Figure 5.**
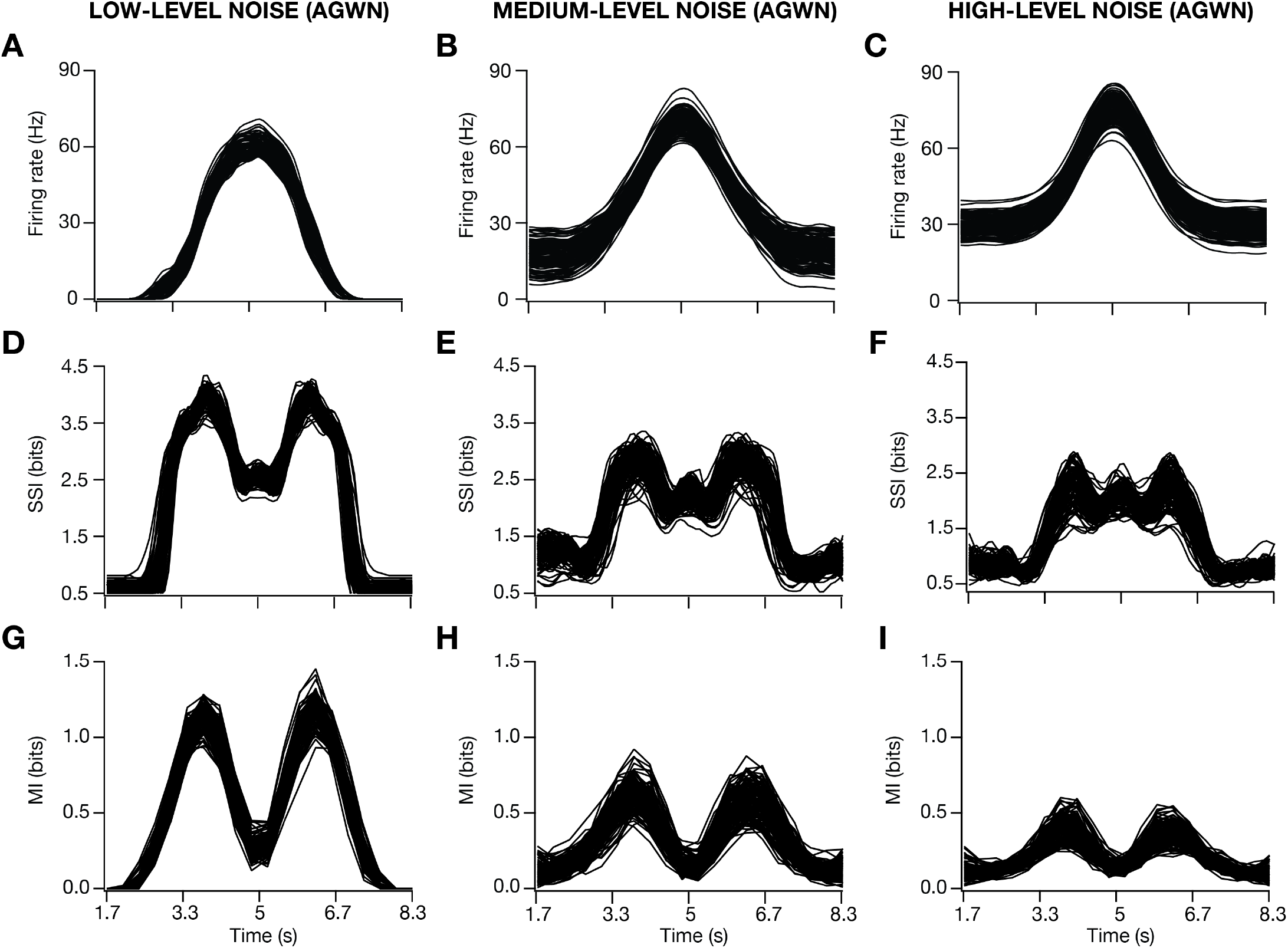
Enhanced trial-to-trial variability, imposed as an additive Gaussian white noise (AGWN), reduced spatial information transfer in place-cell models. (A–I) Firing rate profiles (A–C), stimulus specific information (SSI) profiles (D–F), and mutual information profiles (G–I) as functions of time, shown for low (plots on the left), medium (plots in the middle), high (plots on the right) levels of AGWN. AGWN *σ*_noise_ values: *Low*: 5×10^−4^ Hz^2^, *Medium*: 1×10^−3^ Hz^2^, *High*: 5×10^−3^ Hz^2^.

Similar to the heterogeneities observed in place-cell measurements in the presence of different degrees of trial-to-trial variability (Fig. 4C–D; Fig. 5A–C), we noted marked heterogeneity in spatial information, assessed with the SSI and MI profiles across models (Fig. 5D–I). Importantly, at low levels of trial-to-trial variability, the SSI (Fig. 5D) and the MI (Fig. 5G) showed maximal spatial information transfer at the high-slope locations of the corresponding spatial tuning curves (Fig. 5A). Consequently, both the SSI and the MI profiles were bimodal when low degrees of trial-to-trial variability was introduced, although the values of SSI at high-firing locations were higher compared to MI values at these locations. With increased trial-to-trial variability, introduced as AGWN, the out-of-field firing rates increased (Fig. 5B–C) while also enhancing the peak firing rate (Fig. 5B–C; Fig. 4C).

Progressively enhancing trial-to-trial variability by increasing *σ*_noise_ resulted in a marked reduction in spatial information across models, while still manifesting heterogeneity in spatial information transfer across the model population (Fig. 5E–F; Fig. 5H–I). Whereas the MI profile maintained bimodality despite reduction in the transferred information with higher degrees of trial-to-trial variability (Fig. 5H–I), there was a progressive transition from a bimodal (Fig. 5D) to a trimodal (Fig. 5E–F) distribution of the SSI profiles. The transition in the SSI profile was consequent to the suppression in spatial information transfer at the high-slope locations of the tuning curve, with relatively small changes to spatial information transfer at the high-firing locations (Fig. 5D–F).

To further assess this transition in the SSI profile with enhanced trial-to-trial variability, we increased *σ*_noise_ to larger values and computed the values of the SSI at the high-slope locations (*SSI*_slope_, the average value from the two peaks of the SSI, computed for symmetric firing profile; Fig. 6A) and at the peak-firing locations (*SSI*_peak_; Fig. 6A). We computed the ratio *SSI*_peak_/*SSI*_slope_ and plotted this as a function of *σ*_noise_ (Fig. 6A). A value less than unity for this ratio indicates that maximal stimulus specific spatial information was transferred at the high-slope regions, whereas a value above unity reflects maximal SSI at the peak-firing location. Whereas *SSI*_peak_/*SSI*_slope_ was less than unity for low values of *σ*_noise_ across all models (Fig. 5D, Fig. 6A), two sub-populations of models emerged with higher values of *σ*_noise_. In one subpopulation (*N*=87), *SSI*_peak_/*SSI*_slope_ was always lower than unity even with higher degrees of trial-to-trial variability (teal and orange plots in Fig. 6A, bottom panel; example SSI profiles in Fig. 6B); in a second smaller subpopulation (*N*=27), this ratio was less than unity for low degrees of trial-to-trial variability but transitioned to values higher than unity for higher degrees of trial-to-trial variability (black and purple plots in Fig. 6A, bottom panel; example SSI profiles in Fig. 6C). Thus, whereas a large proportion of models transferred maximal spatial information at the high-slope locations irrespective of the degree of trial-to-trial variability, a subpopulation of models switch to transferring maximal information at the peak-firing locations with higher degrees of trial-to-trial variability.

**Figure 6.**
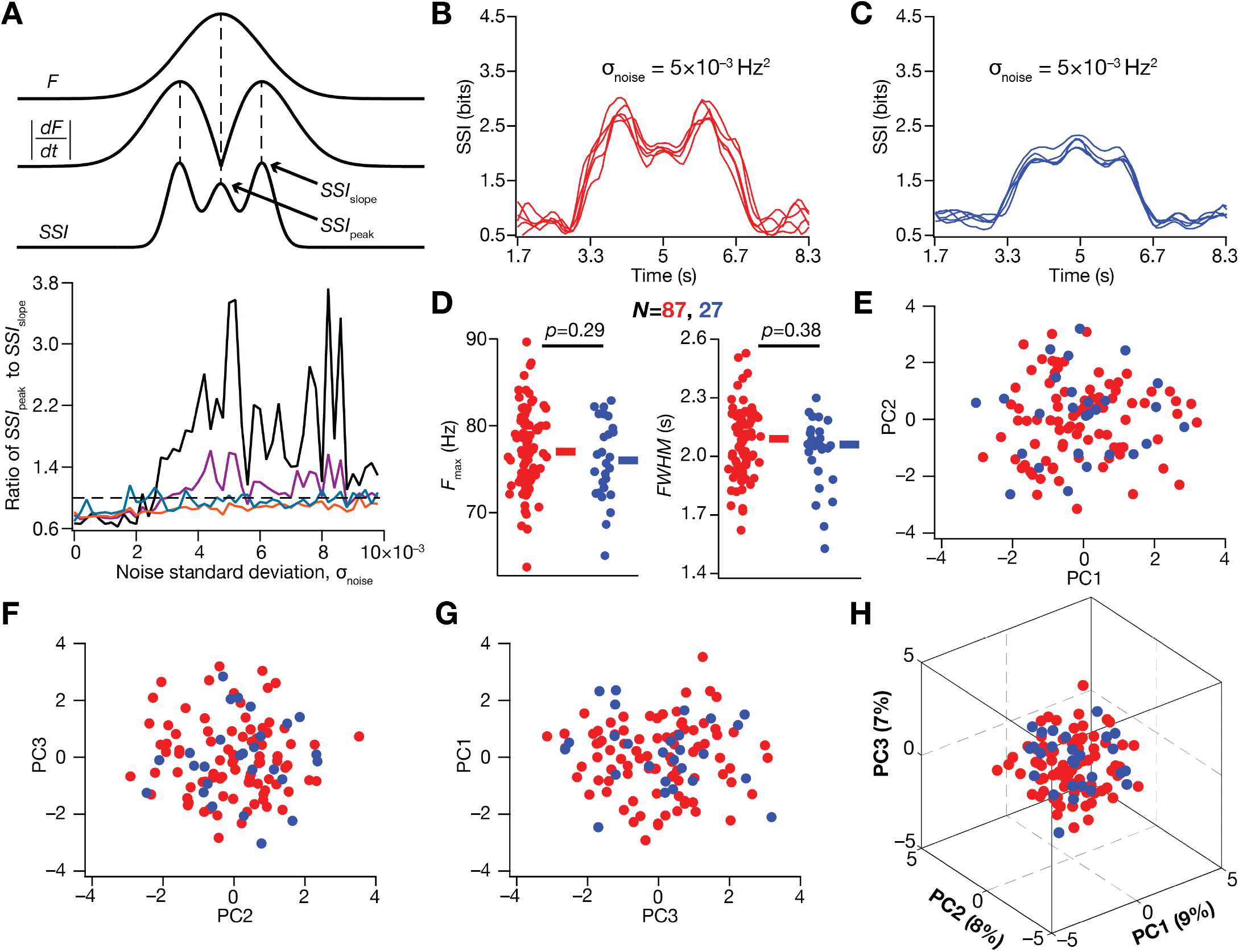
Heterogeneous impact of enhanced trial-to-trial variability on spatial information transfer in place cells. (A) *Top*, Illustration of the measurements *SSI*_peak_ and *SSI*_slope_. *SSI*_peak_ depicts the SSI value at the location where the place-field firing profile (*F*) is at its peak, and *SSI*_slope_ represents the SSI value at the location where the absolute slope of the place-field firing profile, 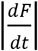, is at its peak. *Bottom*, Traces from four representative models showing the heterogeneity in the evolution of *SSI*_peak_/*SSI*_slope_ as a function of enhanced trial-to-trial variability. (B–C) There were broadly two classes of models, one where the *SSI*_peak_ was low even at high noise levels (B; several representative examples shown in red), and another where *SSI*_peak_ was the highest SSI when noise level was high (C; several representative examples shown in blue). (D) Peak firing rate (left) and FWHM (right) of the two classes of model subpopulations. The rectangles besides each plot represent the respective median value. *σ*_noise_=5×10^−3^ Hz^2^. *p* values provided correspond to the Wilcox rank sum test. (E–H) Principal component analyses on the parameters underlying the two classes of models shown in B (red) and C (blue). Shown are the coefficients associated with these model parameters with reference to the first three principal components. The percentage variance explained by each principal component is provided within parentheses in panel H.

We found that there were no significant differences in the peak firing rate or the width of the place-field firing profiles of models within the two model subpopulations, the ones showing higher SSI at high-slope *vs*. high-firing locations with high degrees of trial-to-trail variability (Fig. 6D). Were there systematic differences in the parameters that defined models within these two subpopulations? To answer this question, we performed principal component analysis (PCA) on parameters that governed the models within the two subpopulations (Fig. 6E–H). We asked if there were distinct clusters representative of the two subpopulations in the reduced dimensional space, pointing to structured parametric differences between these two populations. We found that the three principal dimensions explained merely 24% of the total variance, and there was considerable overlap in the coefficients associated with these two subpopulations, suggesting the absence of systematic parametric differences in the subpopulations (Fig. 6E–H).

We developed 12 distinct profile-specific metrics for quantifying the SSI (Fig. 7A) and MI (Fig. 7H) profiles for the 127 models for three levels of noise. These quantitative metrics confirmed the considerable heterogeneities in spatial information transfer across the model population (Fig. 7). These results showed that across models, information transferred reduced with increase in trial-to-trial variability, with symmetry in spatial information transfer at the two-high slope regions (Fig. 7B–C, Fig. 7I–J). These quantitative metrics also corroborated the emergence of the two subpopulations (Fig. 6) at high values of *σ*_noise_; specifically, the value of *SSIdip* (Fig. 7F) was greater than zero in a small sub-population of models, indicating that these models transfer maximal information at the peak-firing location compared to the high-slope locations (Fig. 7A). The value of *MIdip* (Fig. 7M), however, was always negative across all measured values of *σ*_noise_.

**Figure 7.**
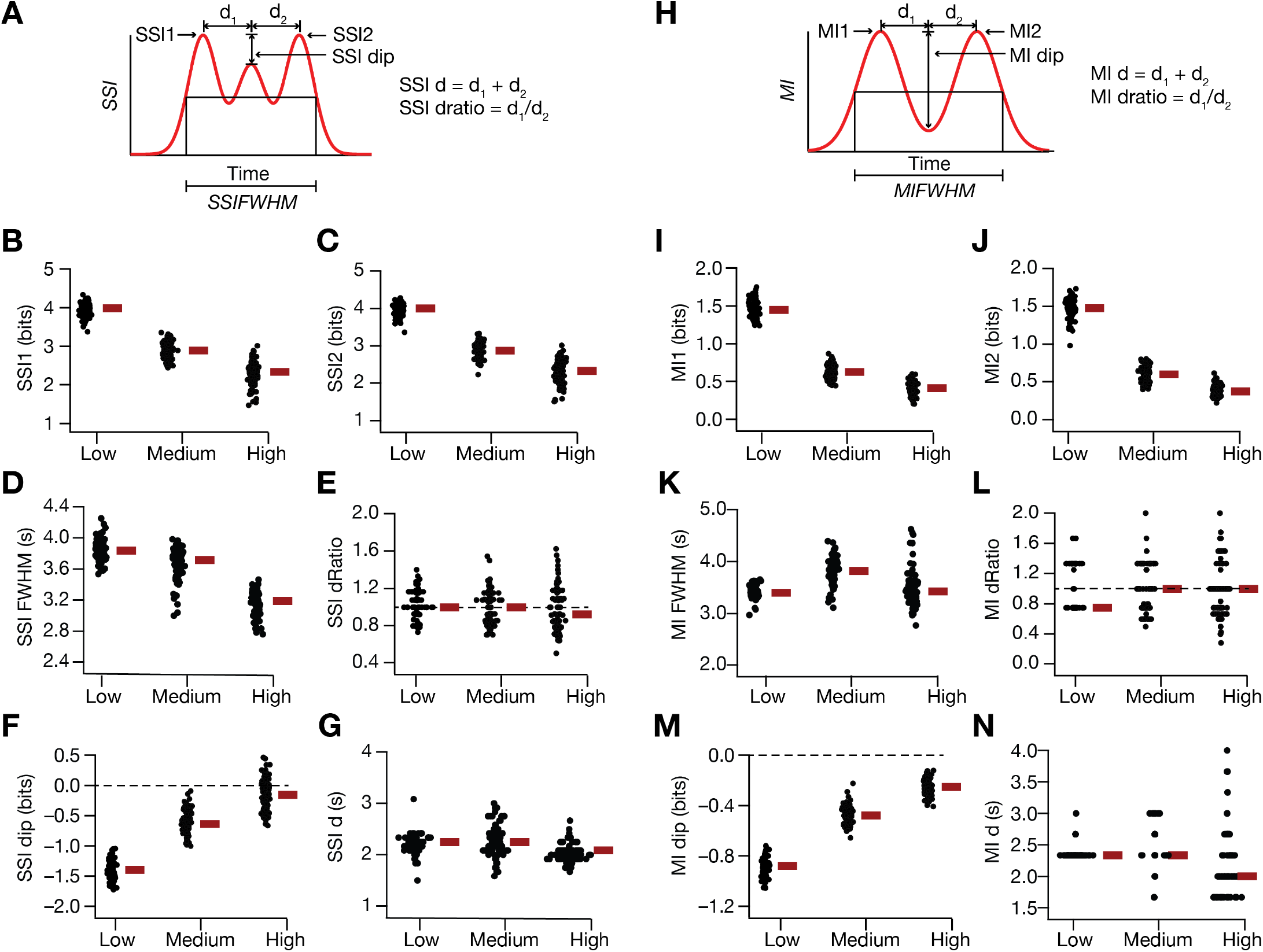
Quantification of the reduction in spatial information transfer as a consequence of enhanced trial-to-trial variability, imposed as an additive Gaussian white noise (AGWN) in place-cell models. (A) Idealized representation of stimulus-specific information (SSI) as a function of time, illustrating the various metrics developed here for quantifying spatial information transfer in place cell models. (B–G) SSI metrics for the population of valid models depicting the impact of three levels of noise on the first (B, *SSI1*) and second (C, *SSI2*) peaks of SSI, the full width half maximum of the SSI profile (D, *SSIFWHM*), the ratio of the first peak-to-center distance to the center-to-second peak distance (E, *SSI dRatio*), the difference between the SSI value at the place field center to the peak SSI value (F, *SSI dip*) and the difference between the location of *SSI1* and *SSI2* (G, *SSI d*). (H–N) Same as (A–G) for mutual information profiles of the valid model population. AGWN *σ*_noise_ values: *Low*: 5×10^−4^ Hz^2^, *Medium*: 1×10^−3^ Hz^2^, *High*: 5×10^−3^ Hz^2^.

Together, our results point to a critical role for the degree of trial-to-trial variability in regulating both qualitative and quantitative aspects of spatial information transfer profile in hippocampal place cells. Specifically, the amount of information transfer progressively reduced with an increase in the degree of trial-to-trial variability, with a qualitative transition from high spatial information transfer occurring at the high-slope to high-firing locations of the spatial tuning curve. In addition, these observations unveil heterogeneities in the place-cell model population in terms of their information transfer capabilities, and in terms of how information transfer changes with increased degrees of trial-to-trial variability. These results emphasize the need to account for biological heterogeneities in neural populations while assessing information transfer characteristics.

### Degeneracy in the emergence of place cells manifesting similar rate-based spatial information transfer profiles

We computed the SSI and MI profiles for the five similar models shown in Fig. S2, and found they possessed similar SSI and MI metrics as well (Table S1). The parametric values of these models, which have similar place-field firing profiles, similar intrinsic measurements and similar spatial information transfer profiles, however were distributed over the entire span of the respective parametric space (Fig. S2G). These point to the ability of disparate combinations of neuronal parameters to provide similar spatial information transfer profiles, thereby pointing to the expression of degeneracy in concomitantly achieving similar intrinsic properties and similar rate-based spatial information transfer in place cells.

In further exploring the dependencies of spatial information transfer on model parameters, we asked if any of the model parameters values would predict spatial information transfer with different degrees of trial-to-trial variability. To do this, we computed pairwise correlations between 20 physiological measurements (3 somato-dendritic measurements of *R*_in_, |*Z*|_max_, *f*_R_, *Q*, Φ_L_ and bAP; *F*_max_ and FWHM for place-field profiles in the absence of noise) that defined the 127 valid models and the 12 information transfer measurements (Table 3) that were obtained from the place-field responses of these models with low (Fig. S3), medium (Fig. S4) and high (Fig. S5) degrees of trial-to-trial variability. Although there were expected strong correlations between some of the information metrics — such as strong positive correlations between *SSI1 vs*. *SSI2* and *SSI1*/*SS2 vs*. *MI1*/*MI2*, and strong negative correlations between *SSI1*/*SSI2 vs*. *SSIdip* across all three values of *σ*_noise_ — the pairwise correlations between information metrics and model measurement values were weak (Fig. S3–S5).

**Table 3:** Quantitative metrics of information transfer.

| Measurement name | Symbol |
| --- | --- |
| **SSI-based information metrics (Fig. 7A)** |  |
| 1<sup>st</sup> peak of the SSI curve | SSI1 |
| 2<sup>nd</sup> peak of the SSI curve | SSI2 |
| Full width at half maximum of the SSI curve | SSI FWHM |
| Ratio of the distance between middle peak with 1<sup>st</sup> peak and the distance between middle peak and 2<sup>nd</sup> peak of the SSI curve | SSI dRatio |
| SSI middle peak value – average of SSI peak values at the slopes | SSI dip |
| Temporal distance between the two peaks in the SSI curve | SSId |
| **MI-based information metrics (Fig. 7H)** |  |
| 1<sup>st</sup> peak of the MI curve | MI1 |
| 2<sup>nd</sup> peak of the MI curve | MI2 |
| Full width at half maximum of the MI curve | MI FWHM |
| Ratio of the distance between middle peak with 1<sup>st</sup> peak and the distance between middle peak and 2<sup>nd</sup> peak of MI curve | MI dRatio |
| MI middle peak value – average of MI peak values at the slopes | MI dip |
| Temporal distance between the two peaks in MI curve | Mid |

Our model contained 80 synaptic locations that were randomly distributed across the apical dendritic arbor. Our outcomes thus far froze synaptic locations at one specific randomized localization and varied ion channel conductances exploring parametric dependencies of spatial information transfer. In another set of simulations, we fixed the model parameters to reflect the base model profile (Table1; Figure 1) and varied localization of the 80 distinct synapses along the dendritic arbor. Specifically, we dispersed the 80 synapses that received the presynaptic afferent inputs across the apical dendritic arbor to 400 combinations of distinct locations, computed the firing rate profile and the information transfer profiles and plotted the associated measurements (Fig. S6). We found that the introduction of heterogeneities in synaptic localization profiles introduced heterogeneities in spatial firing profiles (Fig. S6A–B) and in the spatial information transfer measured through SSI (Fig. S6C–H) or MI metrics (Fig. S6I–N). However, we also noted that spatial firing profiles endowed with similar firing rate and information transfer metrics could be obtained with distinct combinations of synaptic localization profiles.

Together, these results demonstrated the ability of several disparate ion-channel parametric combinations and different synaptic localization profiles to elicit similar place cell firing profiles endowed with similar information transfer profiles, thereby pointing to the expression of parametric degeneracy in the regulation of spatial information transfer in place cells.

### Regulation of spatial information transfer by experience-dependent asymmetry in place-field response profiles

Our simulations thus far resulted in symmetric place field firing profiles (*e.g.*, Fig. 4B) with a symmetric subthreshold voltage ramp (*e.g.*, Fig. 4G), consequent to the symmetric input structure defined by a Gaussian (equation 17). However, electrophysiological lines of evidence from behavioral experiments point to an experience-dependent asymmetric expansion of hippocampal place fields in the direction opposite to the movement of the animal (Harvey, et al., 2009; Mehta, et al., 1997; Mehta, et al., 2002; Mehta, et al., 2000). What is the impact of such experience-dependent asymmetry on spatial information transfer within a single place filed through place-cell rate code? To address this, we first altered the input structure to a horizontally-reflected Erlang distribution (equation 19) which yielded an asymmetric place-field firing (Fig. S7A–B) profile (Seenivasan & Narayanan, 2020). Consistent with our observations with the symmetric place-field firing profile (Fig. 4), enhanced trial-to-trial variability resulted in increase in *F*_max_ (Fig. S7C) accompanied by reductions in FWHM (Fig. S7D), theta power (Fig. S7E–F) and subthreshold ramp voltage (Fig. S7G–H). The subthreshold voltage ramp profile was asymmetric (Fig. S7G), and reflected the asymmetric firing rate profile (Seenivasan & Narayanan, 2020).

We computed the asymmetric firing rate profiles for all valid models with low (Fig. 8A), medium (Fig. 8B) and high (Fig. 8C) degrees of trial-to-trial variability introduced as AGWN to the input structure (equation 18). We found the baseline and the peak firing rates to shift with increased *σ*_noise_, manifesting heterogeneities across models in the populations (Fig. 8A–C). Strikingly, the stimulus-specific information transfer profiles were relatively insensitive to the asymmetry in the firing rate profile (Fig. 8D–F), although the MI profiles reflected the asymmetry (Fig. 8G–I). Specifically, the first and the second peaks were not significantly different for SSI profiles (Fig. 8D–F, Fig. 9A–B; Wilcox signed rank test between first and second peaks: Low: *p*=0.1264; Medium: *p*=0.1383; High: *p*=0.2927), but the second peak was significantly larger than first peak for MI profiles (Fig. 8G–I, Fig. 9G–H; Wilcox signed rank test between first and second peaks: Low: *p* = 2.2×10^−16^; Medium: *p* = 2.2×10^−16^; High: *p* = 5.5×10^−11^) especially for low degrees of trial-to-trial variability.

**Figure 8.**
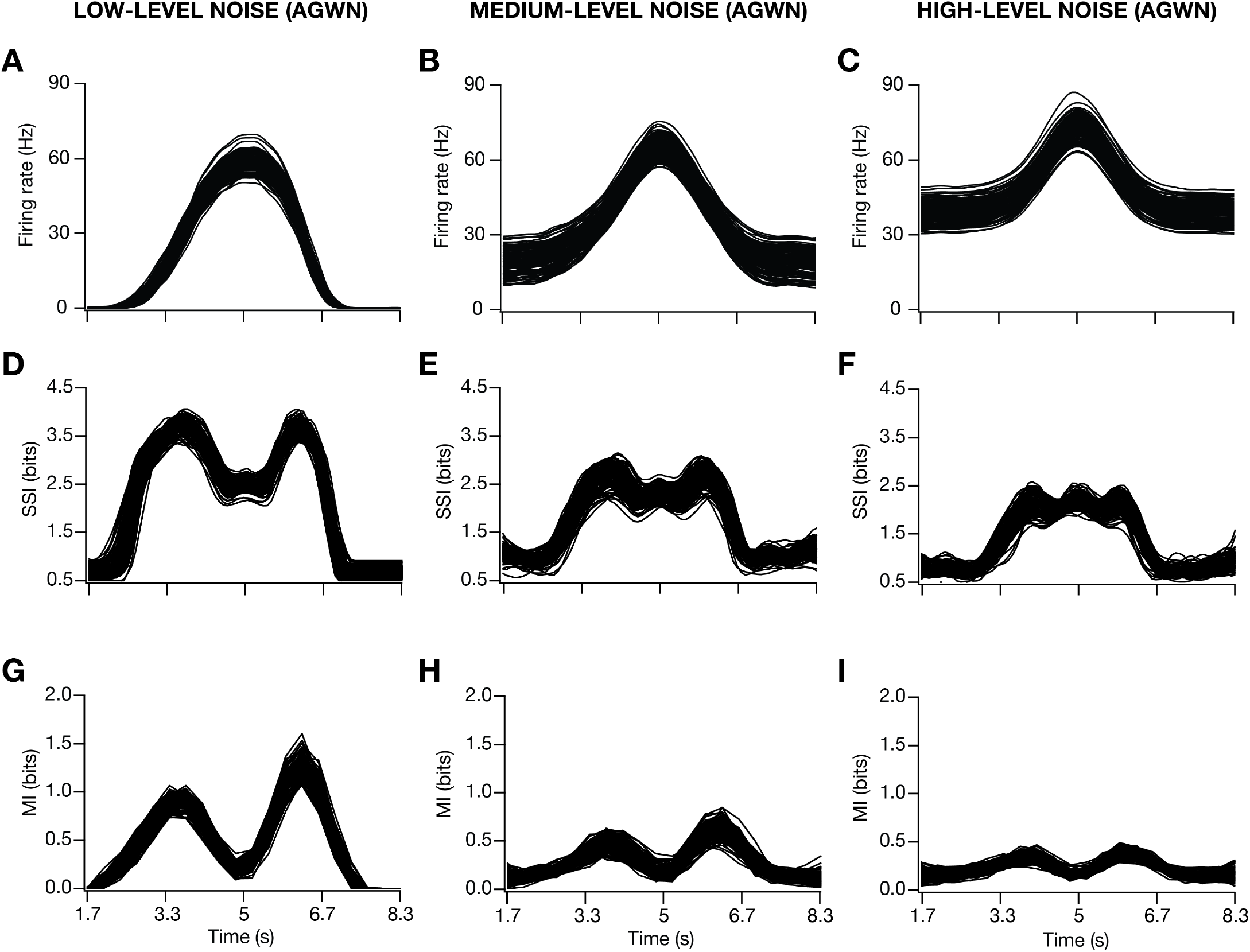
Enhanced trial-to-trial variability, imposed as an additive Gaussian white noise (AGWN), reduced spatial information transfer in models with asymmetric place-field firing. (A–I) Firing rate profiles (A–C), stimulus specific information (SSI) profiles (D–F), and mutual information profiles (G–I) as functions of time, shown for low (plots on the left), medium (plots in the middle), high (plots on the right) levels of AGWN. AGWN *σ*_noise_ values: *Low*: 5×10^−4^ Hz^2^, *Medium*: 1×10^−3^ Hz^2^, *High*: 5×10^−3^ Hz^2^.

**Figure 9.**
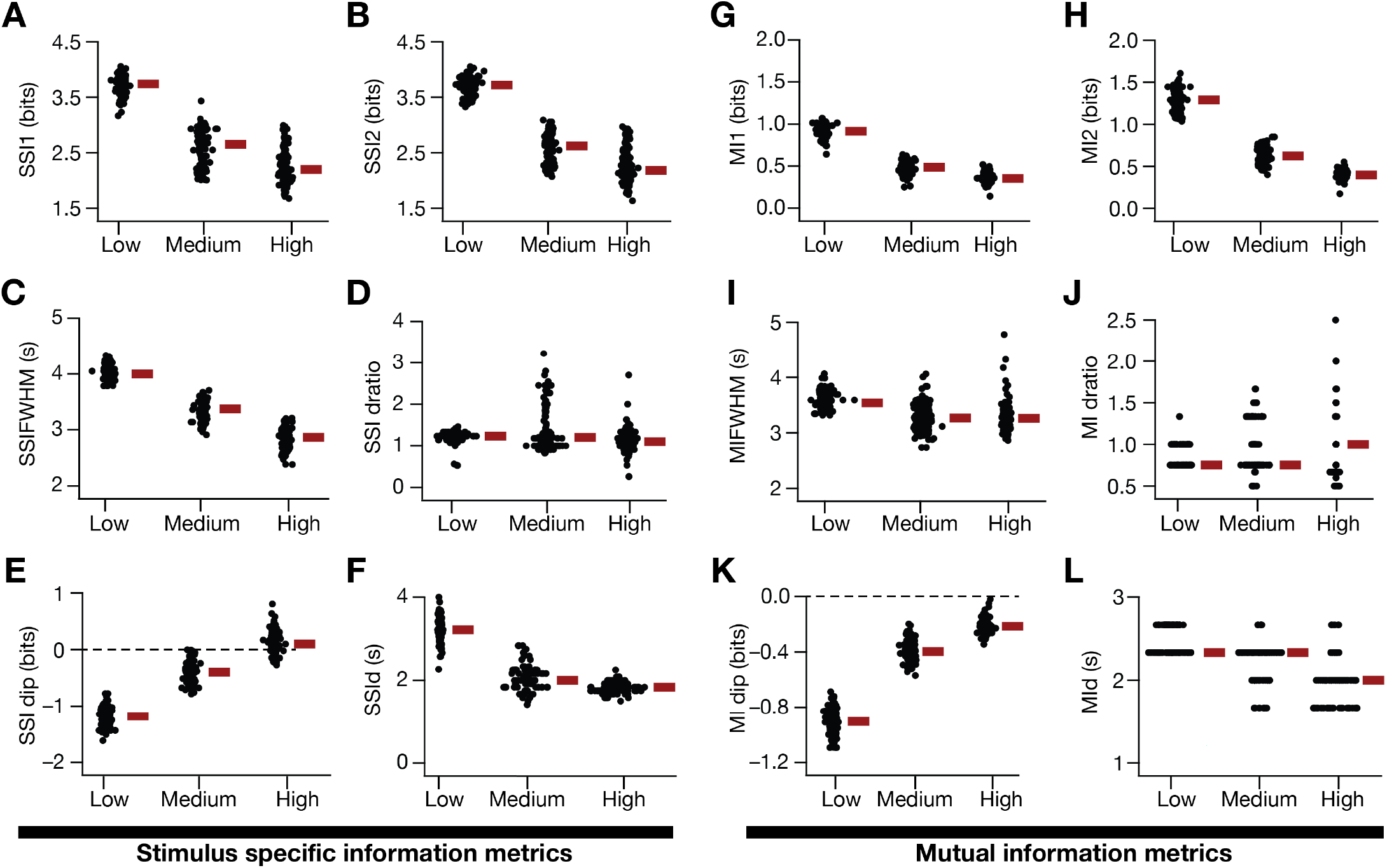
Quantification of the reduction in spatial information transfer as a consequence of enhanced trial-to-trial variability, imposed as an additive Gaussian white noise (AGWN) in models with asymmetric place-field firing. (A–F) SSI metrics for the population of valid models depicting the impact of three levels of noise on the first (A, *SSI1*) and second (B, *SSI2*) peaks of SSI, the full width half maximum of the SSI profile (C, *SSIFWHM*), the ratio of the first peak-to-center distance to the center-to-second peak distance (D, *SSI dRatio*), the difference between the SSI value at the place field center to the peak SSI value (E, *SSI dip*) and the difference between the location of *SSI1* and *SSI2* (F, *SSI d*). (G–L) Same as (A–F) for mutual information profiles of the valid model population. AGWN *σ*_noise_ values: *Low*: 5×10^−4^ Hz^2^, *Medium*: 1×10^−3^ Hz^2^, *High*: 5×10^−3^ Hz^2^.

Consistent with our observations with a symmetric place-field profile, there was marked reduction in spatial information transfer, measured either as SSI or MI (Fig. 8D–I; Fig. 9A–B; Fig. 9G–H), with increased trial-to-trial variability. With low degrees of trial-to-trial variability, we observed that the highest information transfer occurred at the high-slope regions of the firing rate profile, computed either through SSI (Fig. 9E) or MI (Fig. 9K). With increase in degree of trial-to-trial variability, in a manner similar to our findings with symmetric firing profiles (Fig. 6–7) a subpopulation of models switched to transferring maximal SSI at the peak of the firing rate profile (Fig. 9E; High *σ*_noise_; subpopulation with *SSIdip* > 0), but no such transition occurred in the MI profile (Fig. 9K). There were several models which transferred similar amount of spatial information, but were endowed with disparate parametric combination, pointing to the expression of degeneracy with asymmetric firing profiles. Pairwise correlations between model physiological measurements and information metrics were mostly weak, irrespective of the degree of trial-to-trial variability (Fig. S8–S10). Together, these results showed that the introduction of asymmetry in place-field firing profile introduced asymmetries in the spatial information transfer profiles computed through MI, but not through SSI.

### The impact of activity-dependent trial-to-trial variability on spatial information transfer was minimal

We had introduced trial-to-trial variability as an AGWN, whereby the variability was independent of spatial location and synaptic activity (equation 19). To understand the impact of trial-to-trial variability that was dependent on synaptic activity, we introduced trial-to-trial variability as a multiplicative GWN (equation 20) and repeated our analyses on spatial information transfer for the population of valid models, both with symmetric as well as asymmetric firing profiles (Fig. 10, Fig. S11–S20). Although we observed heterogeneity in firing profiles and information transfer, and found models expressing similar information transfer despite being governed by disparate parametric combinations, we found the impact of trial-to-trial variability with the higher range of *σ*_noise_ (compared to those employed for AGWN) to be minimal on place cell properties (Fig. S11), SSI and MI profiles (Fig. 10, Fig. S12, Figs. S16–S17) or pair-wise correlations between intrinsic and information metrics (Figs. S13–S15; Figs. S18–S20). The value of *σ*_noise_ employed for achieving “high” degree of trial-to-trial variability (=0.5 Hz^2^) was the highest possible, as increases beyond that resulted in depolarization-induced block of action potential firing in several models. Experience-dependent asymmetry in firing profiles introduced asymmetry in the MI profiles, but not the SSI profile, even with MGWN-based trial-to-trial variability (Fig. S16–S17). In summary, our results showed that the impact of activity-dependent trial-to-trial variability is minimal compared to activity-independent variability in trial-to-trial responses, across different levels of noise and with symmetric or asymmetric place-field firing profiles.

**Figure 10.**
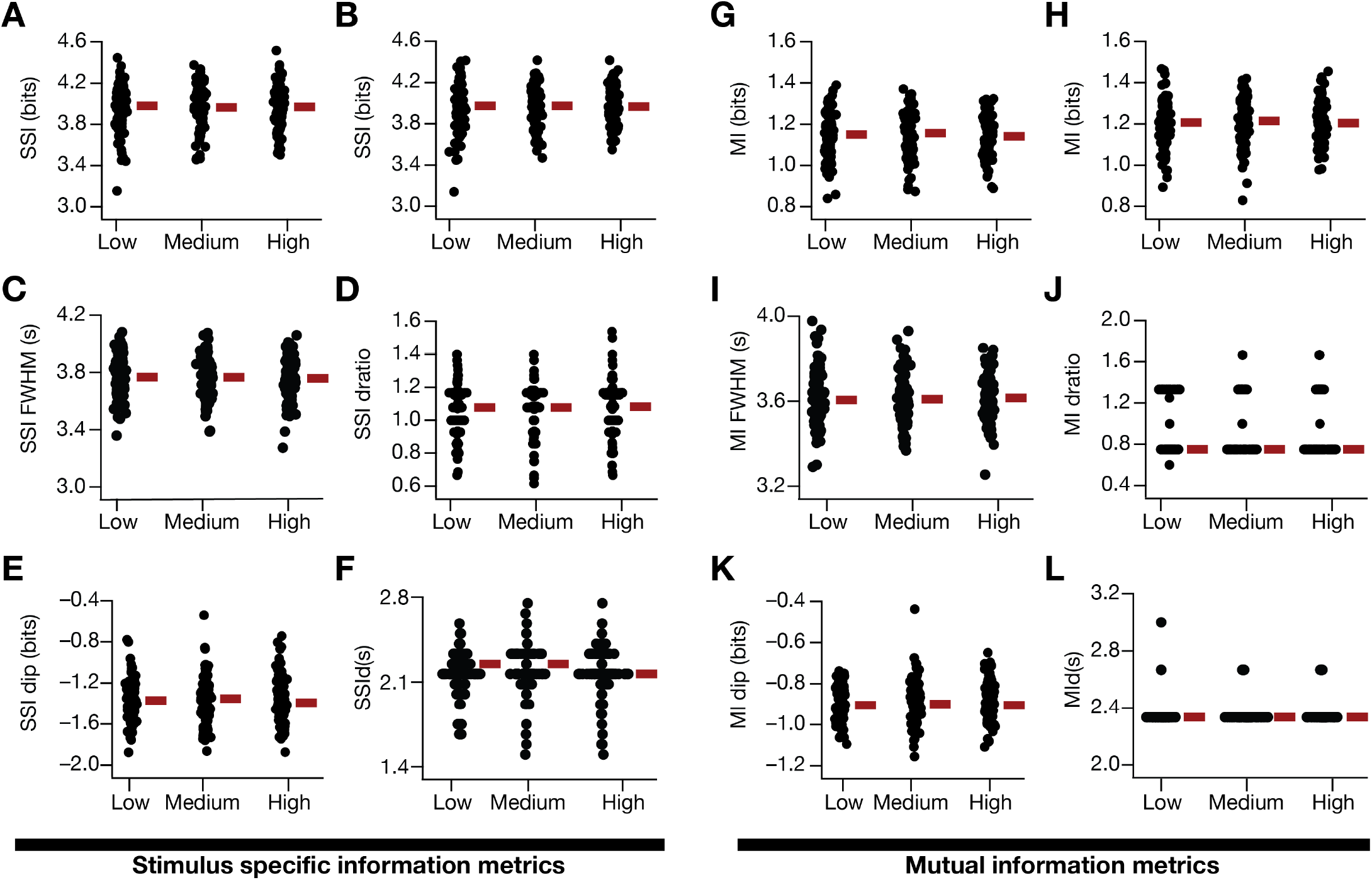
Minimal impact of enhanced activity-dependent trial-to-trial variability, imposed as a multiplicative Gaussian white noise (MGWN), on spatial information transfer. (A–F) SSI metrics for the population of valid models depicting the impact of three levels of noise on the first (B, *SSI1*) and second (C, *SSI2*) peaks of SSI, the full width half maximum of the SSI profile (D, *SSIFWHM*), the ratio of the first peak-to-center distance to the center-to-second peak distance (E, *SSI dRatio*), the difference between the SSI value at the place field center to the peak SSI value (F, *SSI dip*) and the difference between the location of *SSI1* and *SSI2* (G, *SSI d*). (G–L) Same as (A–F) for mutual information profiles of the valid model population. MGWN variance values: *Low*: 0.01 Hz^2^, *Medium*: 0.1 Hz ^2^, *High*: 0.5 Hz ^2^.

### Regulation of spatial information transfer by ion channel conductances and synaptic receptors

Our results established degeneracy in the emergence of place cells with similar spatial information transfer profiles, and also showed an absence of strong correlations with any physiological measurement. What contributes to such degeneracy whereby it is possible for models to achieve similar information transfer profiles despite significant differences in channel expression profiles? Are there specific ion channels that play critical regulatory roles in spatial information transfer within a place field?

We took advantage of our conductance-based modeling framework, and employed the virtual knockout approach (Basak & Narayanan, 2018, 2020; Jain & Narayanan, 2020; Mittal & Narayanan, 2018; Mukunda & Narayanan, 2017; Rathour & Narayanan, 2014; Seenivasan & Narayanan, 2020) to assess the contribution of individual ion channels to spatial information transfer. Specifically, we systematically assessed information transfer profiles in each of the valid models after virtually knocking out individual ion channels by setting their conductance value to zero (Fig. S21). We computed the SSI and MI metrics for the virtual knockout models (VKM) for each of the 8 active ion channels (Fig. 11). Virtual knockout of the spike generating conductances — NaF and KDR — was infeasible because the neuron ceases spiking on setting these conductance values to zero.

**Figure 11.**
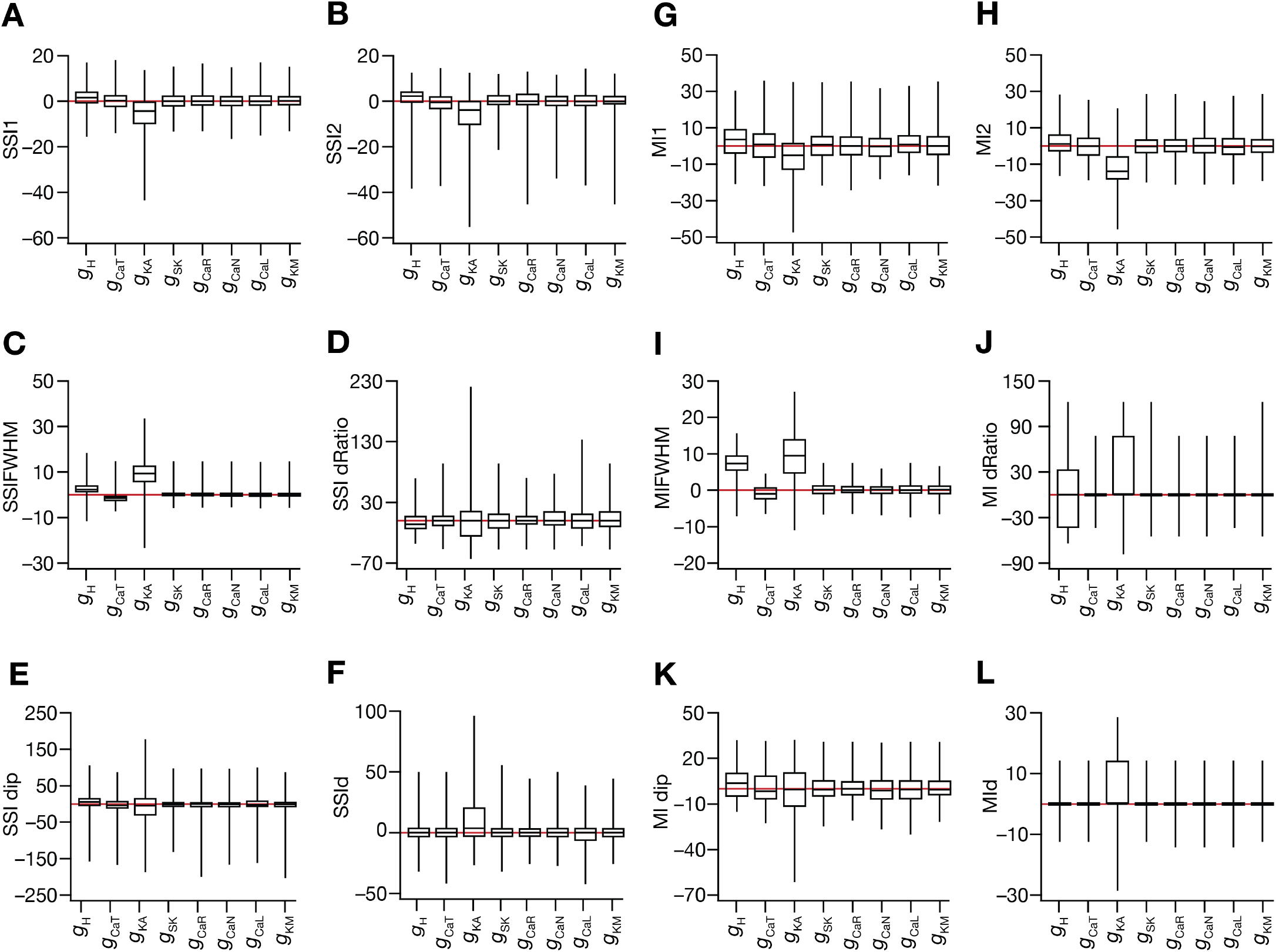
Heterogeneous impact of virtually knocking out individual ion channels on spatial information transfer in the place cell population. (A–F) Box plots showing the median and the quartiles of percentage changes in SSI-based spatial information metrics depicted in Fig. 7A as a consequence of virtually knocking out each of the 8 individual ion channels (NaF and KDR, the spike generating conductances were not knocked out because models cease spiking upon elimination of these channels). (G–L) Box plots showing the median and the quartiles of percentage changes in MI-based spatial information metrics depicted in Fig. 7H as a consequence of virtually knocking out each of the 8 individual ion channels. Plots are shown for the valid place-cell population. Red lines indicate a zero-change scenario.

In terms of information transfer, we found that the impact of knocking out individual channels was heterogeneous across the model population. There were models where the SSI (Fig. 11A–B) or MI (Fig. 11G–H) values increased after knocking out the channel, but there were also models where these values decreased upon knockout. Among the channels assessed, we found the *A*-type potassium channel to have the maximal impact on spatial information transfer. Specifically, virtual knockout of the *A*-type potassium channel resulted in reductions in SSI (Fig. 11A–B) and MI (Fig. 11G–H) values (Wilcoxon signed rank *p* values: *SSI1*: 7.8×10^−9^, *SSI2*: 1.6×10^−10^, *MI1*: 2.7×10^−5^, *MI2*: 6.2×10^−15^), and increased the FWHM values of both SSI (Fig. 11C) and MI (Fig. 11I) profiles (Wilcoxon signed rank test *p* values: *SSIFWHM*: 8×10^−14^, *MIFWHM*: 2.2×10^−16^). These observations offer a clear testable prediction that *A*-type potassium channels play a critical role in regulating spatial information transfer in hippocampal place cells. These results also establish a many-to-one mapping between the different ion channels and the efficacy of spatial information transfer, whereby different ion channels could contribute towards maintaining efficacious information transfer with heterogeneous contributions across neurons in the population. This many-to-one mapping provides a substrate for the expression of degeneracy where different combinations of ion channels could maintain similar functional outcomes in terms of spatial information transfer efficacy.

Finally, as the role of NMDA receptors and dendritic spikes mediated by sodium channels expressed in the dendrites have been considered critical in place-cell physiology (Basak & Narayanan, 2018, 2020; Nakazawa, McHugh, Wilson, & Tonegawa, 2004; M. E. Sheffield & Dombeck, 2015; M. E. J. Sheffield, Adoff, & Dombeck, 2017), we explored the roles of these NMDARs and dendritic NaF channels in regulating spatial information transfer in our heterogeneous model population. To evaluate the role of dendritic fast sodium channels, we recomputed place-field firing rate and spatial information transfer profiles after setting the value of 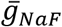 to zero in apical dendritic compartments (Fig. S22A–B). Although there were heterogeneities in the impact of deleting dendritic sodium channels, we found a significant reduction in spatial information transfer computing either as SSI (Fig. 12A–B) or as MI (Fig. 12G–H). To assess the role of NMDARs, we recomputed place-field firing rate and spatial information transfer profiles after setting the value of 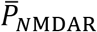 in equations (9–11) to zero (Fig. S22C–D). Deletion of NMDARs resulted in a significant reduction in spatial information transfer (SSI: Fig. 12A–B; MI: Fig. 12G–H).

**Figure 12.**
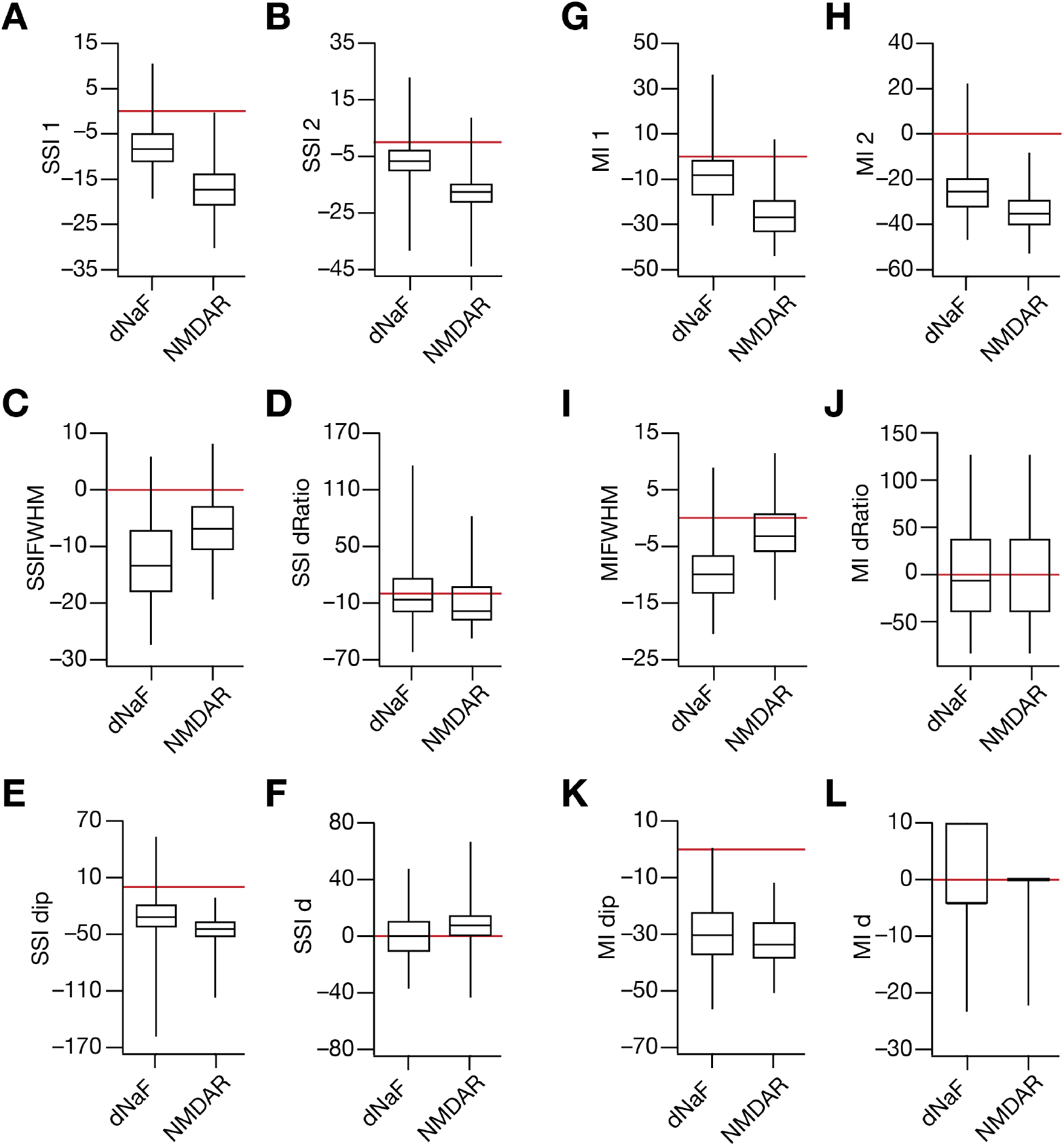
Elimination of dendritic sodium channels or NMDA receptors critically reduces spatial information transfer in the place cell population. (A–L) Box plots showing the median and the quartiles of percentage changes in SSI-based (A–F) and MI-based (G–L) spatial information metrics as a consequence of eliminating dendritic fast sodium channels (dNaF) or NMDA receptors (NMDAR). Plots are shown for the valid place-cell population. Red lines indicate a zero-change scenario.

Together, these results unveiled a many-to-one relationship between the different ion channels and spatial information transfer, while also providing testable predictions on the roles of *A*-type potassium channels, NMDARs and dendritic sodium channels in regulating spatial information transfer within a single place field of hippocampal place cells.

## DISCUSSION

### Conclusions

We demonstrated that hippocampal neurons, when they act as reliable (*i.e.,* low trial-to-trial response variability) sensors of animal location by spatially modulating their firing rate, transfer peak spatial information at the high-slope locations (and not at peak firing location) of the firing rate tuning curve within their place field. However, we showed that there was significant heterogeneity across a population of models that received identical distributions of afferent synaptic patterns, owing to differences in ion channel composition of these models. The heterogeneity manifested in terms of the quantitative value of the amount of information transferred, and in terms of how they responded to increases in the degree of trial-to-trial variability. Specifically, with increases in trial-to-trial variability, whereas one subpopulation of models switched to transferring peak stimulus-specific spatial information at the peak-firing locations, another subpopulation continued to transfer peak information at the high-slope locations. These heterogeneities in spatial information transfer did not show strong relationships between heterogeneities in intrinsic or tuning properties of the models. We demonstrated the dependence of the spatial information transfer profile on the type of trial-to-trial variability, whereby activity-dependent variability had little impact on spatial information transfer compared to the significant reduction introduced by activity-independent variability. Furthermore, we show that the model population manifested parametric degeneracy, whereby models with similar intrinsic measurements, similar tuning curves and similar information transfer metrics exhibited immense heterogeneity in and weak pair-wise correlations across underlying parameters.

To further delineate the relationship of spatial information transfer with place-cell characteristics and its components, we assessed the impact of experience-dependent asymmetry in the place-field firing rate profile. We found that mutual information metrics showed a dependence on the asymmetric nature of the firing profile, where information transfer was maximal in the second half of the place-field where the firing rate dropped at a higher rate. However, the peak values of stimulus-specific information metrics were largely invariant to the asymmetric slopes of the firing rate profile on either side of the peak-firing location. Finally, we asked if there were specific ion channels that played critical roles in regulating spatial information transfer by recomputing information metrics in models that lacked each of 8 different ion channels. We found heterogeneity in the impact of knocking out individual ion channels on these information metrics, pointing to a many-to-one relationship between different ion channel subtypes and spatial information transfer. Our analyses unveiled a potent reduction in information transfer consequent to knocking out transient potassium channels, NMDA receptors or dendritic sodium channels, providing direct experimentally testable predictions.

Together, our analyses emphasize the need to account for neuronal heterogeneities in assessing sensory information transfer and show that synergistic interactions among several neural components regulate the specifics of information transfer and its relationship to tuning curve characteristics. In what follows, we present the implications of our study to place-cell physiology, also outlining the limitations of our study and potential directions for the future.

### Trial-to-trial variability and spatial information transfer

Our results show that trial-to-trial variability in neural responses results in a marked reduction in spatial information transfer within a single place-field, in a manner that is dependent on how the noise was introduced. In demonstrating this, we had introduced trial-to-trial variability employing either an additive or a multiplicative GWN. The incorporation of synaptic additive noise is physiologically similar to a scenario where there is either a location-independent increase in afferent excitation or a reduction in tonic or spatially-uniform inhibition (Duguid, Branco, London, Chadderton, & Hausser, 2012; Grienberger, Milstein, Bittner, Romani, & Magee, 2017). Such a scenario, which could be a result of physiological plasticity or pathological synaptopathies, would enhance response variability in a location-independent manner. Our results demonstrate that the presence of such location- and activity-independent enhancement in trial-to-trial variability critically reduces spatial information transfer within a place field, irrespective of whether the place field profiles are symmetric (Fig. 5, Fig. 7) or asymmetric (Figs. 8–9). With enhanced trial-to-trial variability of this form, our results show that the location of maximal SSI transitions from the high-slope regions to the peak-firing location in a subpopulation of models (Fig. 6).

In striking contrast, incorporation of trial-to-trial variability as a multiplicative noise had little impact on spatial information transfer for a wide range of noise variance values, and the location of maximal SSI was always tuned to the high-slope regions of the tuning curve (Fig. 10). Multiplicative noise, activity-dependent trial-to-trial variability, is physiologically similar to noise consequent to variability in synaptic release and receptor kinetics. In such a scenario, the amount of variability is dependent on the extent of synaptic activation, and therefore is activity-dependent. In place cells, as excitatory afferent activity is higher within the place field of the neuron (highest at the center of the place field), such multiplicative noise translates to location-dependent variability in neural responses. Our results show that the ability of such activity-dependent noise, especially with strong excitatory drives observed during place-field traversal, in altering spatial information transfer is minimal.

These results emphasize the importance of assessing the source of trial-to-trial variability and asking whether the variability is dependent or independent of activity, and caution against a generalization of all types of trial-to-trial variability to yield similar outcomes. Further explorations on the dependence of spatial information transfer on the specific types and sources of variability should account for several experimental details, some of which are listed below. First, although we consider two mutually exclusive versions of trial-to-trial variability (dependent or independent of activity), variability in neuronal responses under awake, behaving conditions is conceivably a mixture of both versions. Second, there are theoretical and electrophysiological lines of evidence for a critical role for asynchronous synaptic release, induced by active reverberation in recurrent circuits (such as the CA3, a presynaptic counterpart to the CA1 neurons studied here), on information transfer (Lau & Bi, 2005; Volman & Levine, 2009). Third, there are lines of evidence of stimulus independent noise improving the detection of subthreshold stimulus (Stacey & Durand, 2000, 2001, 2002). Fourth, although we employed white noise sources in our analyses, it has been demonstrated that the color of the noise is a critical determinant of how information transfer is affected (Gingl, Kiss, & Moss, 1995). Finally, in our analyses the trial-to-trial variability was introduced solely as synaptic noise. However, other factors such as noisy biochemical processes and stochasticity of ion channels could also contribute to the trial-to-trial variability, with different noise color and different ways of interactions with the inputs (Faisal, Selen, & Wolpert, 2008). It is essential that future studies incorporate these additional layers of mechanisms to the model and examine how different sources of variability, each with potentially different characteristics, synergistically affect stimulus-specific information content. It is possible that one or the other version dominates under specific physiological/pathological conditions, and therefore it is important that the variability-inducing mechanisms are delineated before the impact of such variability is assessed.

### Place-cell characteristics and spatial information transfer

An important insight obtained from our study pertains to parametric degeneracy in effectuating spatial information transfer in place cells, with reference to ion channels and parameters that govern place cell biophysics and physiology. Specifically, we demonstrate that several disparate combinations of ion channel conductances and model parameters could elicit similar spatial information transfer across locations through the place-cell firing rate profile within a place field (Figs. S1–S2; Fig. 3). Ion-channel degeneracy in the hippocampal formation is ubiquitous, and expresses across different scales of analyses (Mishra & Narayanan, 2019; Mittal & Narayanan, 2018; Rathour & Narayanan, 2019). In hippocampal CA1 pyramidal neurons, the expression of degeneracy has been demonstrated with reference to the concomitant emergence of several somato-dendritic intrinsic properties (R. Migliore, et al., 2018; Rathour, Malik, & Narayanan, 2016; Rathour & Narayanan, 2012, 2014; Srikanth & Narayanan, 2015), spike-triggered average (Das & Narayanan, 2014, 2015, 2017; Das, Rathour, & Narayanan, 2017; Jain & Narayanan, 2020), short-(Mukunda & Narayanan, 2017) as well as long-term (Anirudhan & Narayanan, 2015) plasticity profiles. More specifically, with reference to the firing properties of CA1 pyramidal neurons as place cells, degeneracy has been shown to express in the sharpness of place-field firing properties with reference to biophysical as well as morphological parameters (Basak & Narayanan, 2018, 2020), which has been confirmed in this study with a larger set of ion channels incorporated into the model. Finally, from the spatial encoding and information transfer perspective, an earlier study had quantitatively defined efficiency of phase coding in hippocampal place cells and showed that similar spatial information transfer could be achieved with disparate ion channel combinations (Seenivasan & Narayanan, 2020). The findings of this study, demonstrating ion channel degeneracy with reference to spatial information transfer through the rate code within a single place field, further strengthen the expression of degeneracy in encoding systems such as the hippocampus.

In encoding systems, it is essential that encoding of information occur concurrently with maintenance of homeostasis of intrinsic neuronal properties, including neuronal firing rate (Rathour & Narayanan, 2019). In our study, we showed that similar amounts of spatial information transfer and similar firing rate (both with reference to place-field firing and responses to pulse currents) could concomitantly occur with disparate combinations of ion channel conductances and parameters that govern their expression (Table S1, Fig. S2). It has been shown that the balance between excitation, inhibition and intrinsic excitability (E–I–IE balance) is essential for achieving concomitant efficient phase coding as well as activity homeostasis. In our study, we had fixed the excitatory synaptic weights to account for synaptic democracy (Fig. 1I) and did not incorporate spatially-uniform inhibition (Grienberger, et al., 2017) as this would have translated to merely a negative bias term across locations (Basak & Narayanan, 2018). We also found that there were no correlations between information measurements and other intrinsic measurements (*e.g.*, Figs. S3–S5). Future studies could alter excitatory synaptic weights associated with place-field inputs and explore the balance between excitation, location-dependent inhibition and the heterogeneous intrinsic excitability properties of hippocampal pyramidal neurons to assess the role of E–I–IE in the emergence of efficient information transfer through rate codes as well. Specifically, such studies could validate models based on their ability to transfer maximal spatial information through the rate code (*i.e.*, efficient rate coding) *and* concomitantly maintain intrinsic homeostasis, and ask if E–I–IE was essential to achieve these when the search space involves excitatory/inhibitory synaptic weights and ion channel conductances (Seenivasan & Narayanan, 2020). Importantly, such models could maximize the *joint* spatial information transfer occurring through the rate as well as the phase codes (Mehta, et al., 2002; O’Keefe & Burgess, 2005) within a place field, and explore the constraints required for such efficient encoding to occur simultaneously with the expression of intrinsic homeostasis.

Degeneracy in the emergence of similar spatial information transfer and signature intrinsic properties emerged as a consequence of a many-to-one relationship between ion channels and spatial information transfer. This was inferred from our analysis with virtual knockout models (Fig. 11–12), whereby removal of any of the several ion channels resulted in heterogeneously altering spatial information transfer in the model population. These results emphasize the importance of recognizing the many-to-one relationship between underlying parameters and information transfer. These observations were feasible only because we employed a heterogeneous population of models, derived from an unbiased stochastic search that covered heterogeneities in the underlying parameters. If we had employed a single hand-tuned model to arrive at our conclusions, that single model and its specific composition would have biased our results. In such a scenario, the identification of the aforementioned many-to-one relationship and the consequent heterogeneities on the impact of individual ion channels on information transfer wouldn’t have been feasible. These results emphasize the critical role of synergistic interactions among different ion channels in effectuating behavior, and underscore that the impact of any ion channel subtype is dependent on the relative expression profiles of other channels and receptors in the specific model under consideration.

Degenerate systems show dominance of specific underlying parameters in regulating *specific* physiological measurements (Basak & Narayanan, 2018, 2020; Drion, O’Leary, & Marder, 2015; Mishra & Narayanan, 2019; Mittal & Narayanan, 2018; Mukunda & Narayanan, 2017; Rathour, et al., 2016; Rathour & Narayanan, 2014, 2019). In our analyses, although we found that all ion channels had the ability to reduce or increase spatial information transfer in a model-dependent manner (Fig. 11–12), certain parameters played a crucial role in regulating information transfer. Specifically, our analyses provide specific experimentally testable predictions on the critical roles of dendritic sodium channels, NMDA receptors and *A*-type potassium channels in regulating spatial information transfer (Figs. 11–12). Interestingly, these three components play critical roles in regulating the prevalence of dendritic spikes and in the sharpness of place-cell tuning profiles (Basak & Narayanan, 2018, 2020; Gasparini, Migliore, & Magee, 2004; Golding, Jung, Mickus, & Spruston, 1999; Golding & Spruston, 1998; Losonczy & Magee, 2006), and form strong candidates in regulating spatial information transfer. Further studies could test the roles of these channels in regulating information transfer in hippocampal pyramidal neurons employing electrophysiological recordings during place-field traversal in the presence of pharmacological agents. As these components alter dendritic spiking in opposite directions (suppressing NMDA receptors or sodium channels suppresses dendritic spiking, whereas suppression of *A*-type potassium channels enhances dendritic spiking), such studies could also potentially assess the requirement of an intricate balance between mechanisms that promote and those that prevent dendritic spike initiation in maintaining efficient spatial information transfer (Basak & Narayanan, 2020).

Our results proffer a testable prediction that experience-dependent asymmetry in place-field profiles does not markedly alter SSI, but alters MI. As experience-dependent asymmetry is considered to be predictive, reduction in spatial information transfer during the early parts of place-field firing would have rendered this predictive capability to be ineffectual. Our observations demonstrate that although the low values of slope during the early parts of firing profile reduces mutual information as a consequence of the asymmetry, stimulus specific information remains high. Further explorations could test this prediction on electrophysiologically obtained individual place cells transitioning with experience (Mehta, et al., 1997), employing different metrics of spatial information transfer.

Finally, an important outcome of the analyses presented here relates to the widespread prevalence of biological heterogeneities across all scales of analysis, and all physiological metrics. These heterogeneities are ubiquitous, spanning ion channel expression and localization profiles, morphological characteristics, neuronal intrinsic properties, tuning properties with reference to external features (in our case, space), information transfer profiles, the location of high-information transfer (high-slope *vs*. peak-firing), responses of information metrics to trial-to-trial variability and quantitative metrics associated with these properties. Therefore, it is extremely critical to account for these heterogeneities in experimental analyses and in computational models of brain function, at all scales of analyses spanning the genes-to-behavior range. With specific reference to hippocampal place fields, future studies should explore the impact of systematic gradients in neuronal properties and ion channel expression along the dorso-ventral, proximo-distal and superficial-deep axes of the hippocampus (Cembrowski, et al., 2016; Cembrowski & Spruston, 2019; Danielson, et al., 2016; Dougherty, Islam, & Johnston, 2012; Dougherty, et al., 2013; Kjelstrup, et al., 2008; Lee, et al., 2014; Malik, Dougherty, Parikh, Byrne, & Johnston, 2016; Marcelin, et al., 2012; Maroso, et al., 2016; Mizuseki, Diba, Pastalkova, & Buzsaki, 2011; Strange, Witter, Lein, & Moser, 2014; Sun, et al., 2017) on spatial information transfer within a single place field. Furthermore, the question on how spatial information transfer is regulated by activity-dependent plasticity and behavioral state-dependent neuromodulation of ion channels and receptors is critical in understanding the emergence of spatial information transfer in the context of novel place-field formation (Bittner, et al., 2015; Bittner, Milstein, Grienberger, Romani, & Magee, 2017; Cohen, Bolstad, & Lee, 2017; M. E. J. Sheffield, et al., 2017; Zhao, Wang, Spruston, & Magee, 2020).

## Acknowledgments

This work was supported by the DBT-Wellcome Trust India Alliance (Senior fellowship to RN; IA/S/16/2/502727), Human Frontier Science Program (HFSP) Organization (RN), the Department of Biotechnology through the DBT-IISc partnership program (RN), the Revati and Satya Nadham Atluri Chair Professorship (RN), the Ministry of Human Resource Development (RN) and Kishore Vaigyanik Protsahan Yojana (AR). The authors thank the members of the cellular neurophysiology laboratory for helpful discussions and for comments on a draft of this manuscript.

