## Supplementary material for "Spatial information transfer in hippocampal place cells depends on trial-to-trial variability, symmetry of place-field firing and biophysical heterogeneities": Figures S1-S22, Table S1

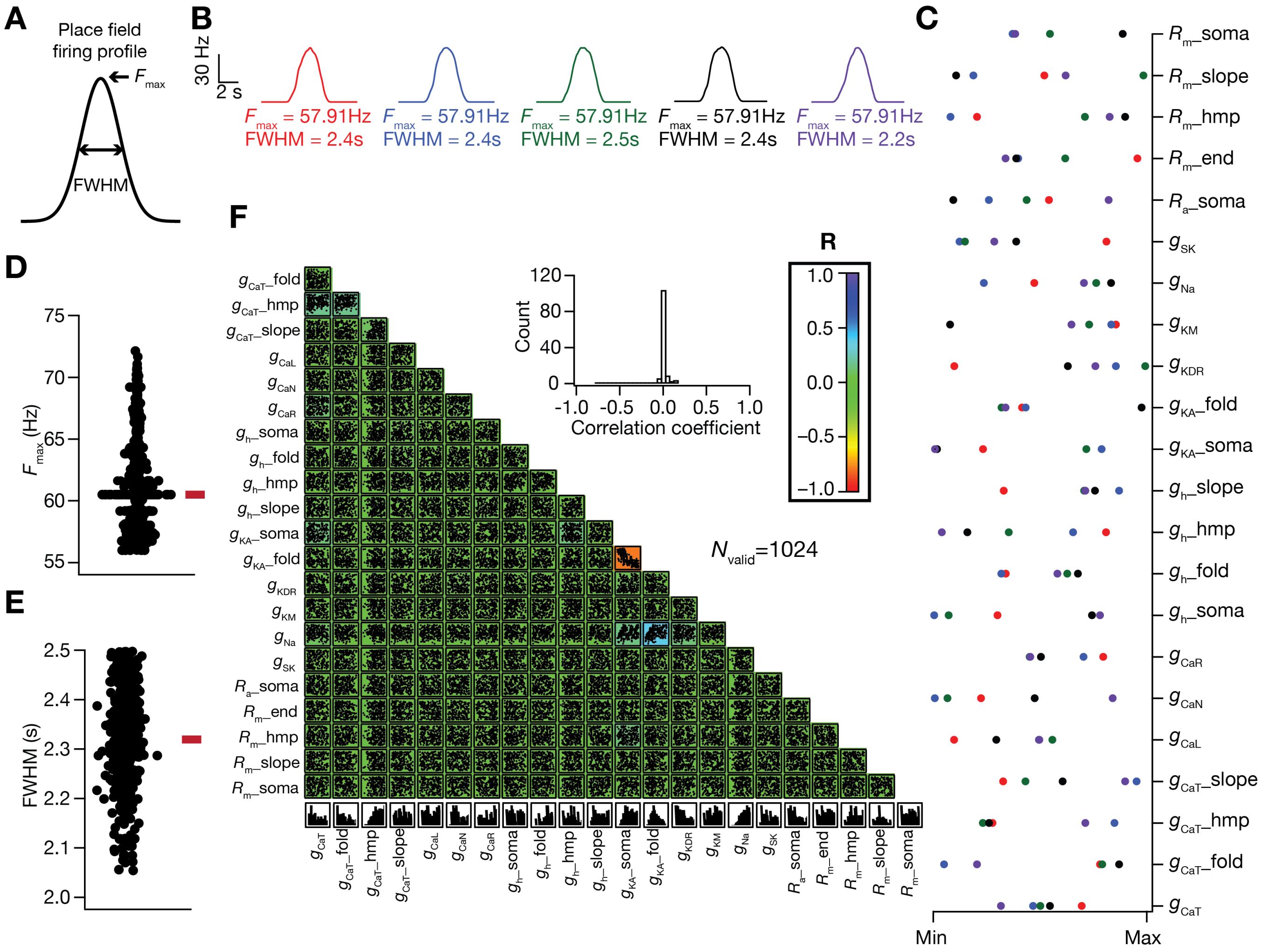

**Supplementary Figure S1. Similar sharp tuning of place-field firing was obtained with disparate combinations of neuronal active and passive properties.** (A) A typical place-cell firing profile illustrating the measurement of maximum firing rate ($F_{max}$) and the temporal distance between the places with half the maximum value of firing rate (*FWHM*). Based on a relative criterion on tuning sharpness, involving high $F_{max}$ (>56 Hz) and low *FWHM* (<2.5 s), 1024 sharply-tuned valid place-cell (models out of the 12000 randomly generated models) models were obtained. (B) Place-cell firing profiles of 5 models with similar place-cell measurements. (C) The values of parameters that defined the 5 models depicted in B. It may be noted that the values of these model parameters spanned a large span of the allocated range (Table 1), although their functional characteristics (C) were similar. (D–E) $F_{max}$ (D) and FWHM (E) of all the 1024 valid models. (F) Pairwise scatter plot matrix superimposed on top of the color-coded pairwise correlation coefficient matrix for the 1024 models that exhibited sharp place-field tuning. The inset shows a histogram of all the correlation coefficients.

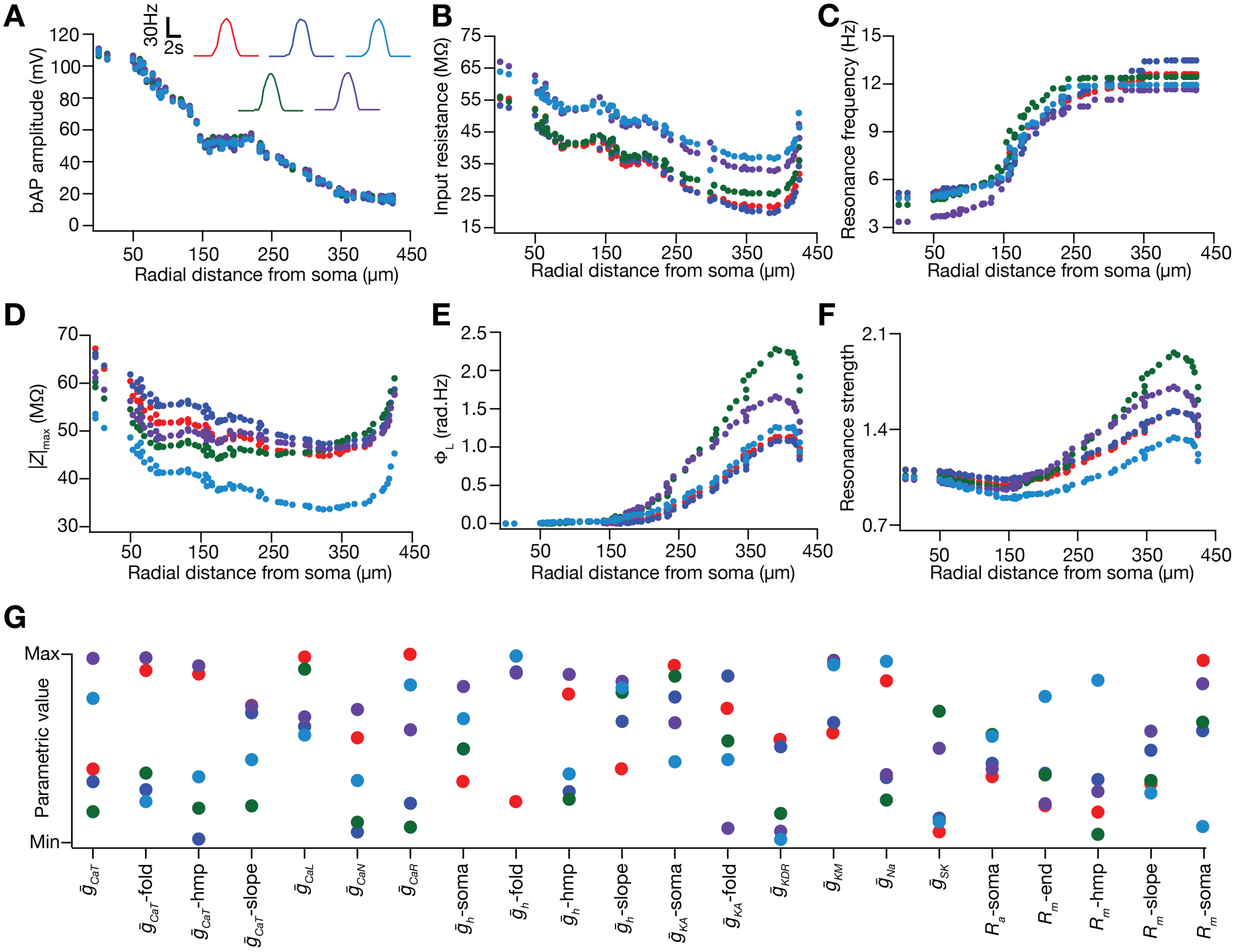

**Supplementary Figure S2. Expression of ion channel degeneracy in the concomitant emergence of sharply-tuned place-cell firing profiles and several functional maps of intrinsic neuronal properties.** Five models with similar place cell tuning profile (A, insets; $F_{max}$= 57.25, 57.25, 57.37, 58.06, 57.91 Hz; *FWHM*= 2.33, 2.25, 2.25, 2.25, 2.42 s for the five models) and similar intraneuronal functional maps of backpropagating action potential amplitude (A), input resistance (B), resonance frequency (C), maximum impedance amplitude (D), total inductive phase (E) and strength of resonance (F). (G) The values of the 22 parameters that defined the 5 models depicted in A–F. It may be noted that the values of these model parameters spanned a large span of the allocated range (Table 1), although their functional characteristics (A–F) were similar.

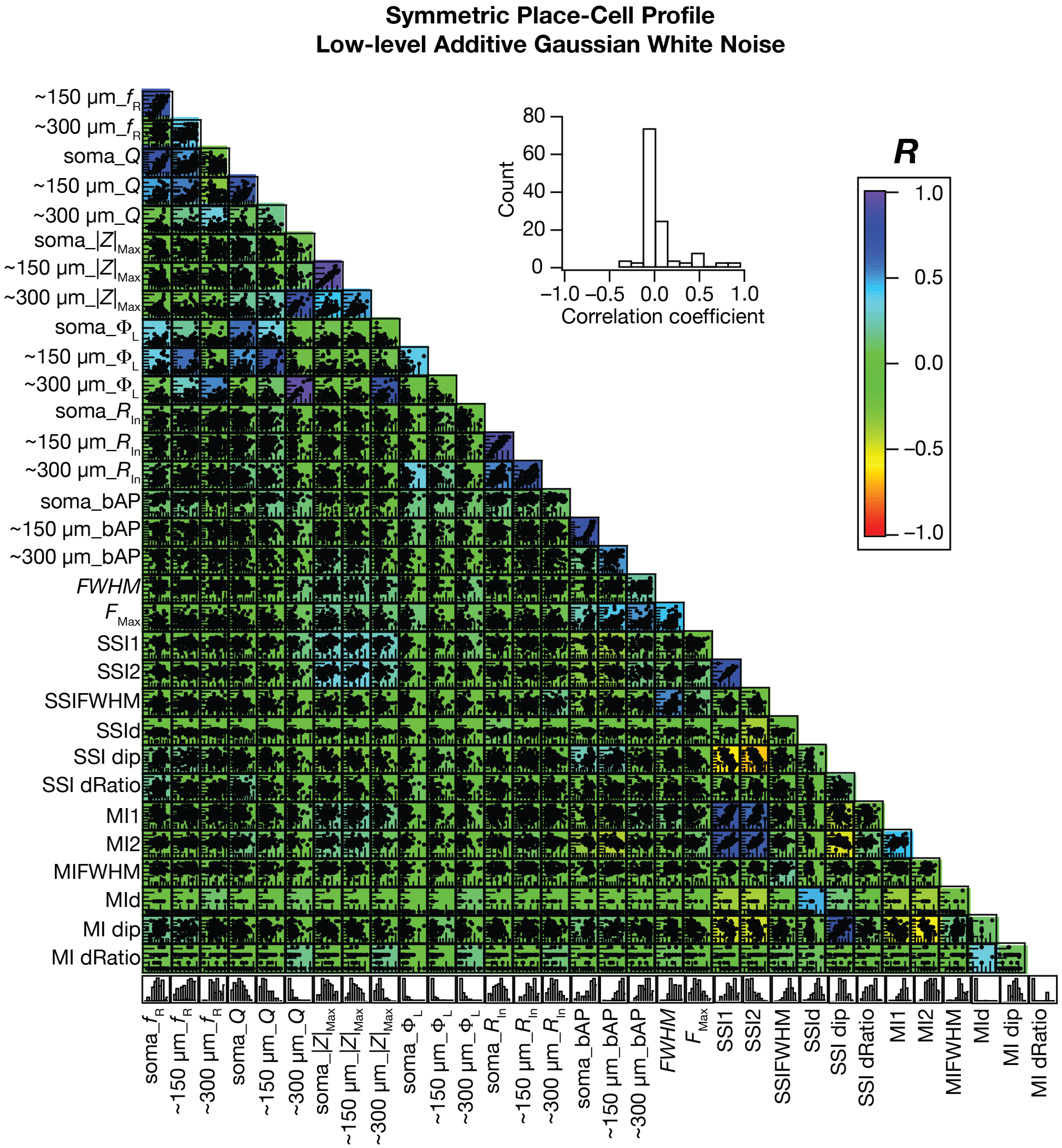

**Supplementary Figure S3. Impact of low-level additive Gaussian white noise on pairwise correlations between intrinsic measurements, place-cell firing properties and information measurements.** Pairwise scatter plot matrix of intrinsic physiological measures, place cell firing properties and information measures (defined in Fig. 7) superimposed on the corresponding correlation coefficient matrix for the 127 valid models. Inset shows the histogram of all the correlation coefficient values.

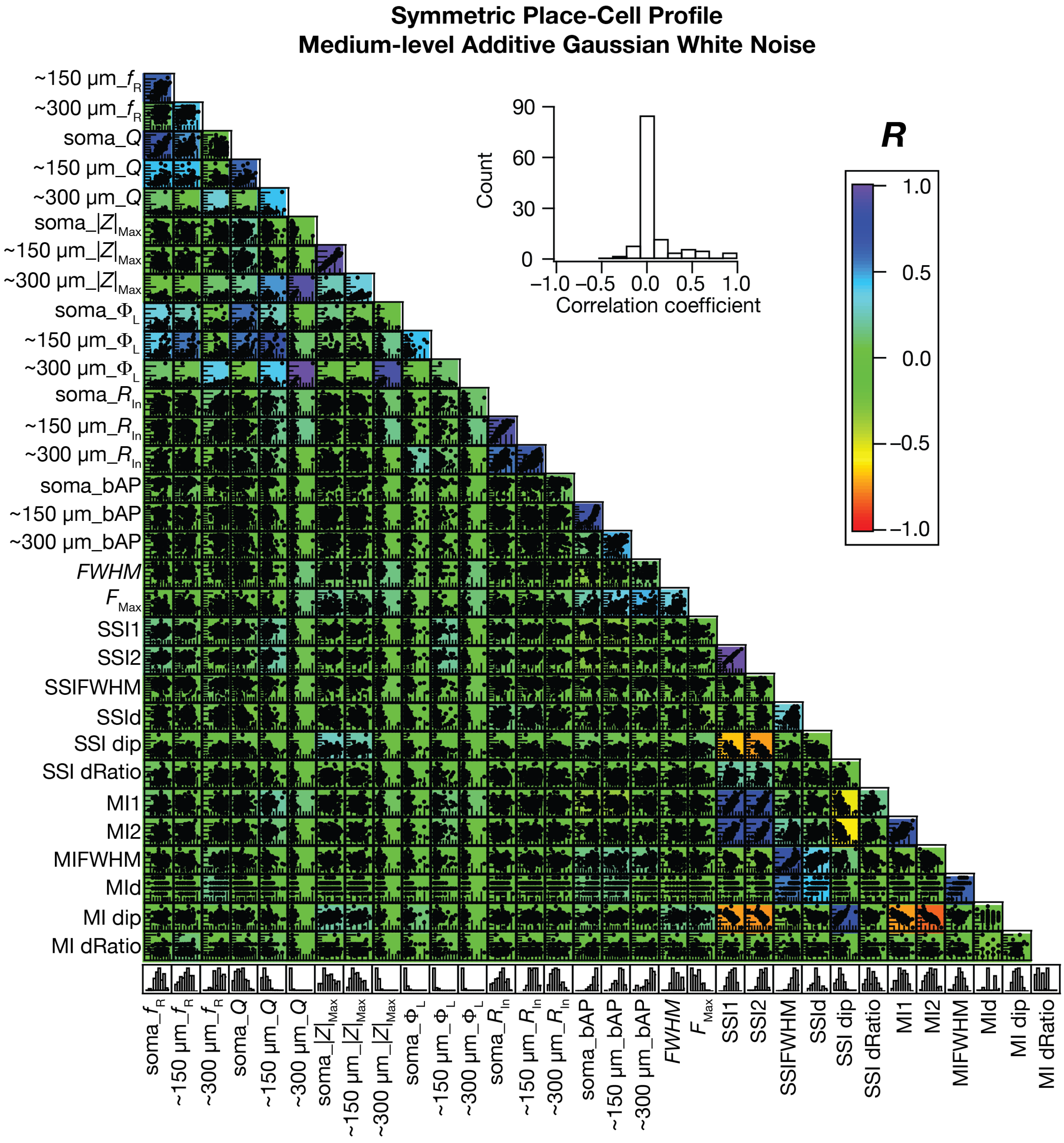

**Supplementary Figure S4. Impact of medium-level additive Gaussian white noise on pairwise correlations between intrinsic measurements, place-cell firing properties and information measurements.** Pairwise scatter plot matrix of intrinsic physiological measures, place cell firing properties and information measures (defined in Fig. 7) superimposed on the corresponding correlation coefficient matrix for the 127 valid models. Inset shows the histogram of all the correlation coefficient values.

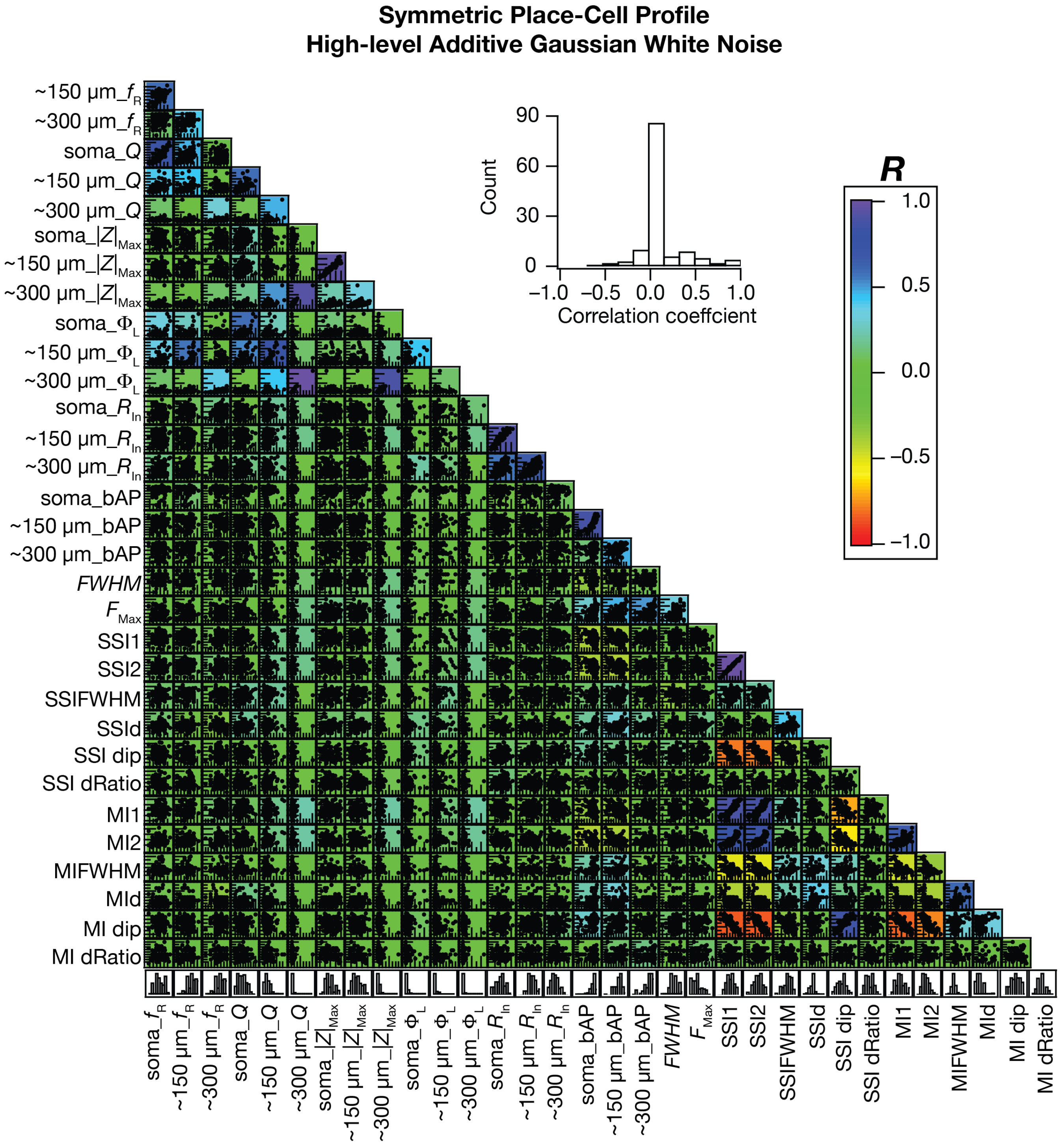

**Supplementary Figure S5. Impact of high-level additive Gaussian white noise on pairwise correlations between intrinsic measurements, place-cell firing properties and information measurements.** Pairwise scatter plot matrix of intrinsic physiological measures, place cell firing properties and information measures (defined in Fig. 7) superimposed on the corresponding correlation coefficient matrix for the 127 valid models. Inset shows the histogram of all the correlation coefficient values.

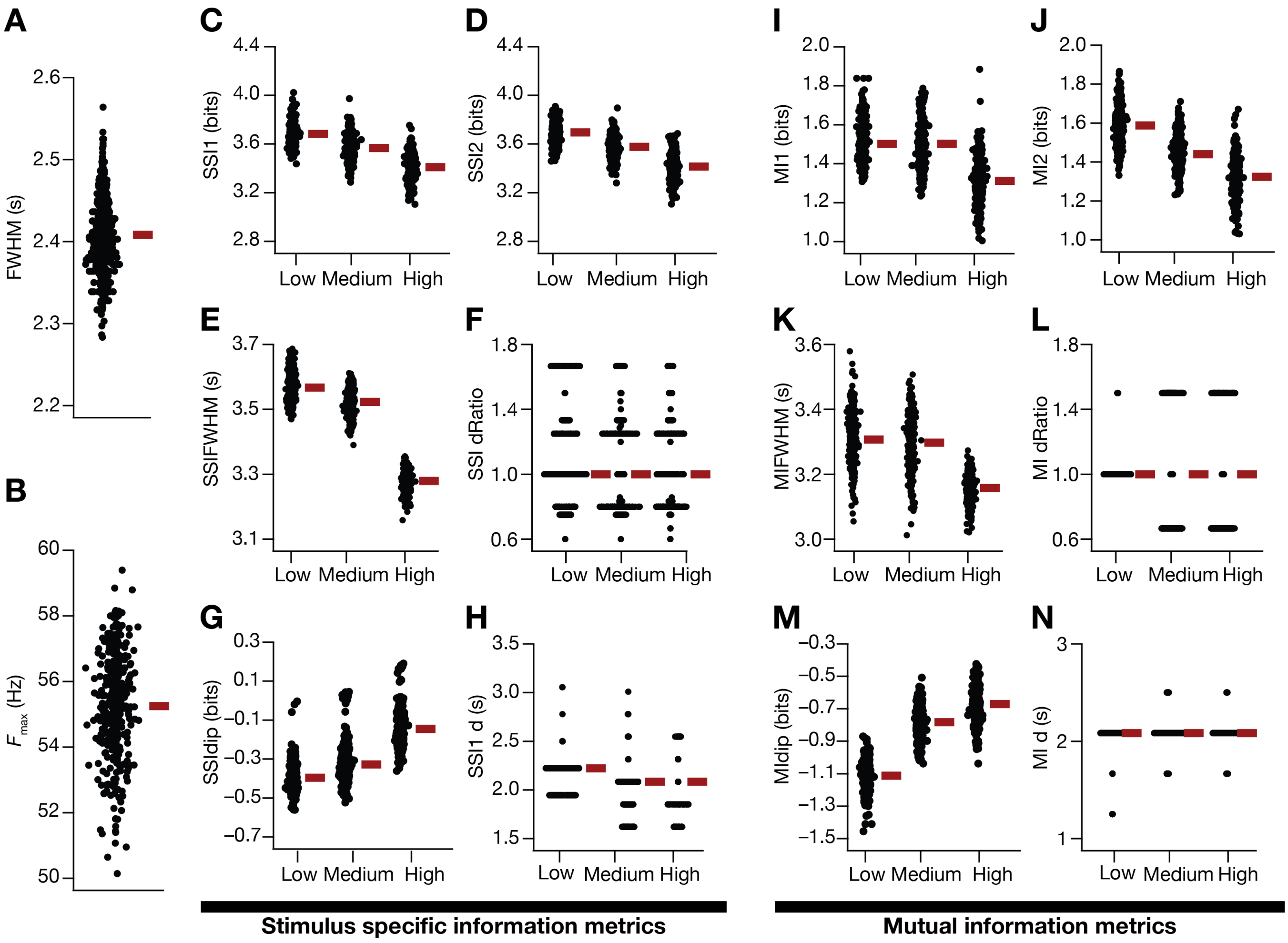

**Supplementary Figure S6. Quantification of the reduction in spatial information transfer as a consequence of enhanced trial-to-trial variability in models endowed with heterogeneity in synaptic localization profiles.** (A–B) Peak firing rate (A) and FWHM (B) of models endowed with heterogeneity in synaptic localization profiles. (C–H) SSI metrics for the population of valid models depicting the impact of three levels of noise on the first (C, *SSI1*) and second (D, *SSI2*) peaks of SSI, the full width half maximum of the SSI profile (E, *SSIFWHM*), the ratio of the first peak-to-center distance to the center-to-second peak distance (F, *SSI dRatio*), the difference between the SSI value at the place field center to the peak SSI value (G, *SSI dip*) and the difference between the location of *SSI1* and *SSI2* (H, *SSI d*). (I–N) Same as (C–H) for mutual information profiles of the valid model population. AGWN $\sigma_{\text{noise}}$ values: *Low*: $5{\times}10^{-4}$ Hz$^2$, *Medium*: $1{\times}10^{-3}$ Hz$^2$, *High*: $5{\times}10^{-3}$ Hz$^2$.

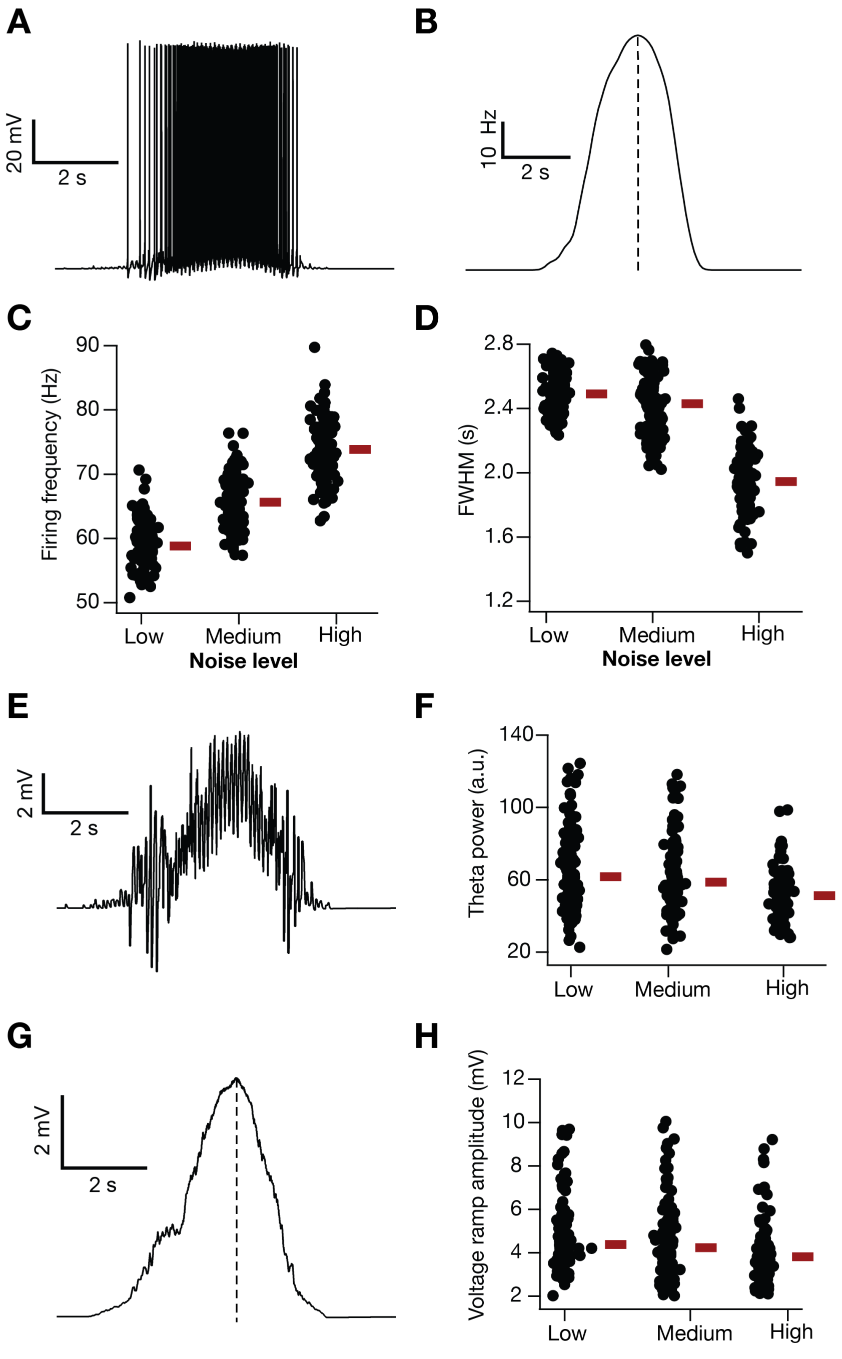

**Supplementary Figure S7. Impact of additive Gaussian white noise (AGWN) on place-cell models manifesting asymmetric place-field tuning profiles.** (A–B) Voltage trace (A) and corresponding firing rate profile (B) during traversal of a place field in a typical valid place-cell model in the presence of AGWN ($\sigma_{\text{noise}} = 5 \times 10^{-4}$ Hz$^2$). Notice asymmetry in firing profile where the temporal duration before the place-field center (dotted line) is larger than that after the place-field center. (C–D) Impact of different levels of AGWN on the peak firing frequency, $F_{\max}$ (C) and full-width at half maximum, *FWHM* (D) of the 127 valid place-cell models. The red bars represent the respective median values. $F_{\max}$: Kruskal Wallis test, $p$=2.3×$10^{-16}$, Wilcoxon Signed Rank test, Low *vs*. Medium $p$=3.4×$10^{-4}$, Medium *vs*. High $p$=4.6×$10^{-4}$, Low *vs*. High = 9.2×$10^{-6}$. FWHM: Kruskal Wallis test, $p$=3.2×$10^{-10}$, Wilcoxon Signed Rank test, Low *vs*. Medium $p$=5.6×$10^{-3}$, Medium *vs*. High $p$=7.9×$10^{-14}$, Low *vs*. High = 6.2×$10^{-15}$. (E) Voltage profile in (A) filtered to emphasize theta-frequency oscillations during traversal of a place field. (F) Impact of different levels of AGWN on theta power of the 127 valid place-cell models. Kruskal Wallis test, $p$=6.9×$10^{-6}$, Wilcoxon Signed Rank test, Low *vs*. Medium $p$=1.9×$10^{-4}$, Medium *vs*. High $p$=2.3×$10^{-5}$, Low *vs*. High = 3×$10^{-6}$. (G) Voltage profile in (A) filtered to emphasize subthreshold voltage ramp during traversal of a place field. Notice asymmetry in voltage profile where the temporal duration before the place-field center (dotted line) is larger than that after the place-field center. (H) Impact of different levels of AGWN on voltage ramp amplitude of the 127 valid place-cell models. Kruskal Wallis test, $p$=7.7×$10^{-6}$, Wilcoxon Signed Rank test, Low *vs*. Medium $p$=4×$10^{-3}$, Medium *vs*. High $p$=6.6×$10^{-3}$, Low *vs*. High = 3.2×$10^{-5}$. When present, the red bars represent the respective median values. AGWN $\sigma_{\text{noise}}$ values: *Low*: 5×$10^{-4}$ Hz$^2$, *Medium*: 1×$10^{-3}$ Hz$^2$, *High*: 5×$10^{-3}$ Hz$^2$.

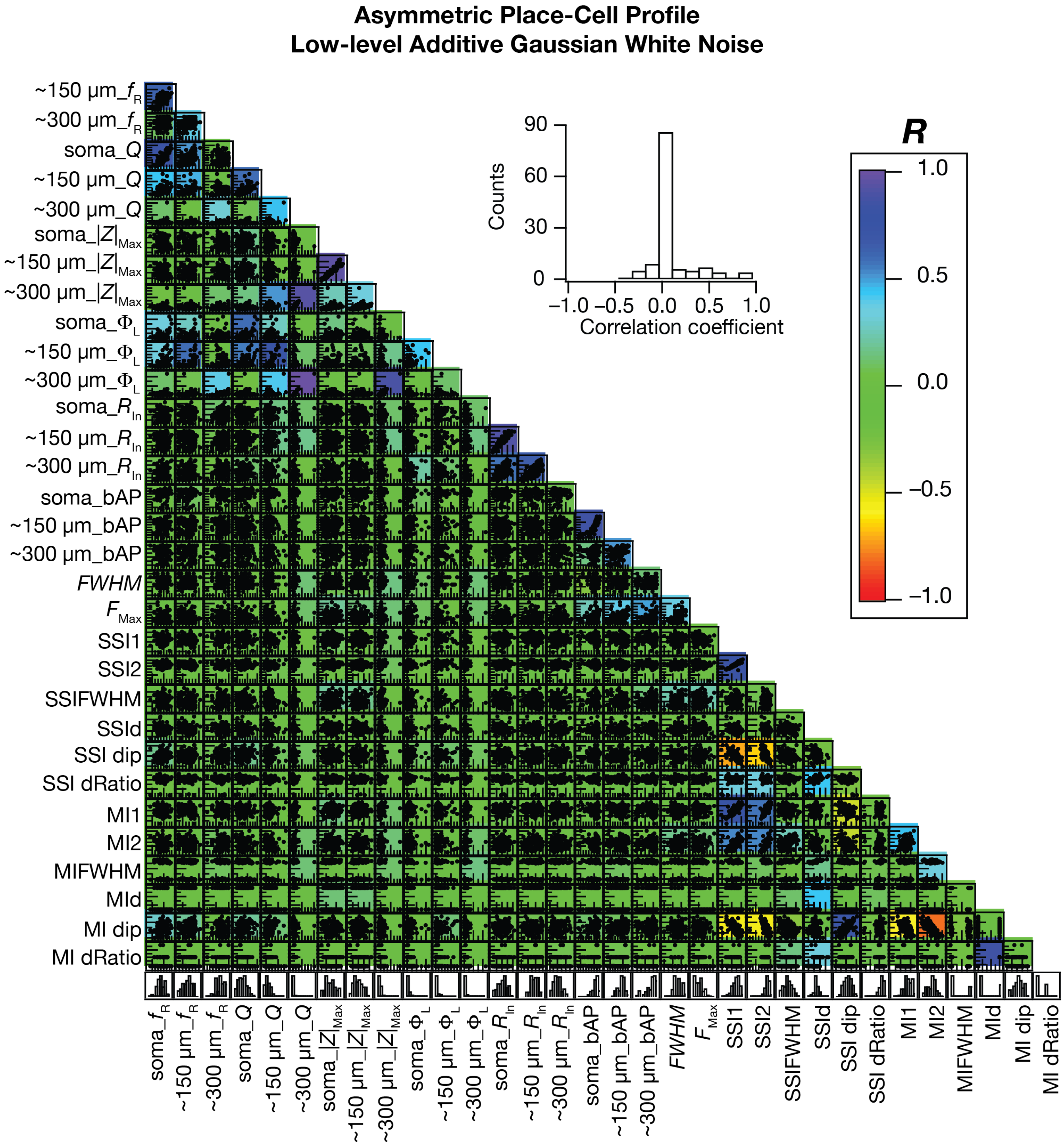

**Supplementary Figure S8. Impact of low-level additive Gaussian white noise on pairwise correlations between intrinsic measurements, place-cell firing properties and information measurements, in models with asymmetric place-field firing.** Pairwise scatter plot matrix of intrinsic physiological measures, place cell firing properties and information measures (defined in Fig. 7) superimposed on the corresponding correlation coefficient matrix for the 127 valid models. Inset shows the histogram of all the correlation coefficient values.

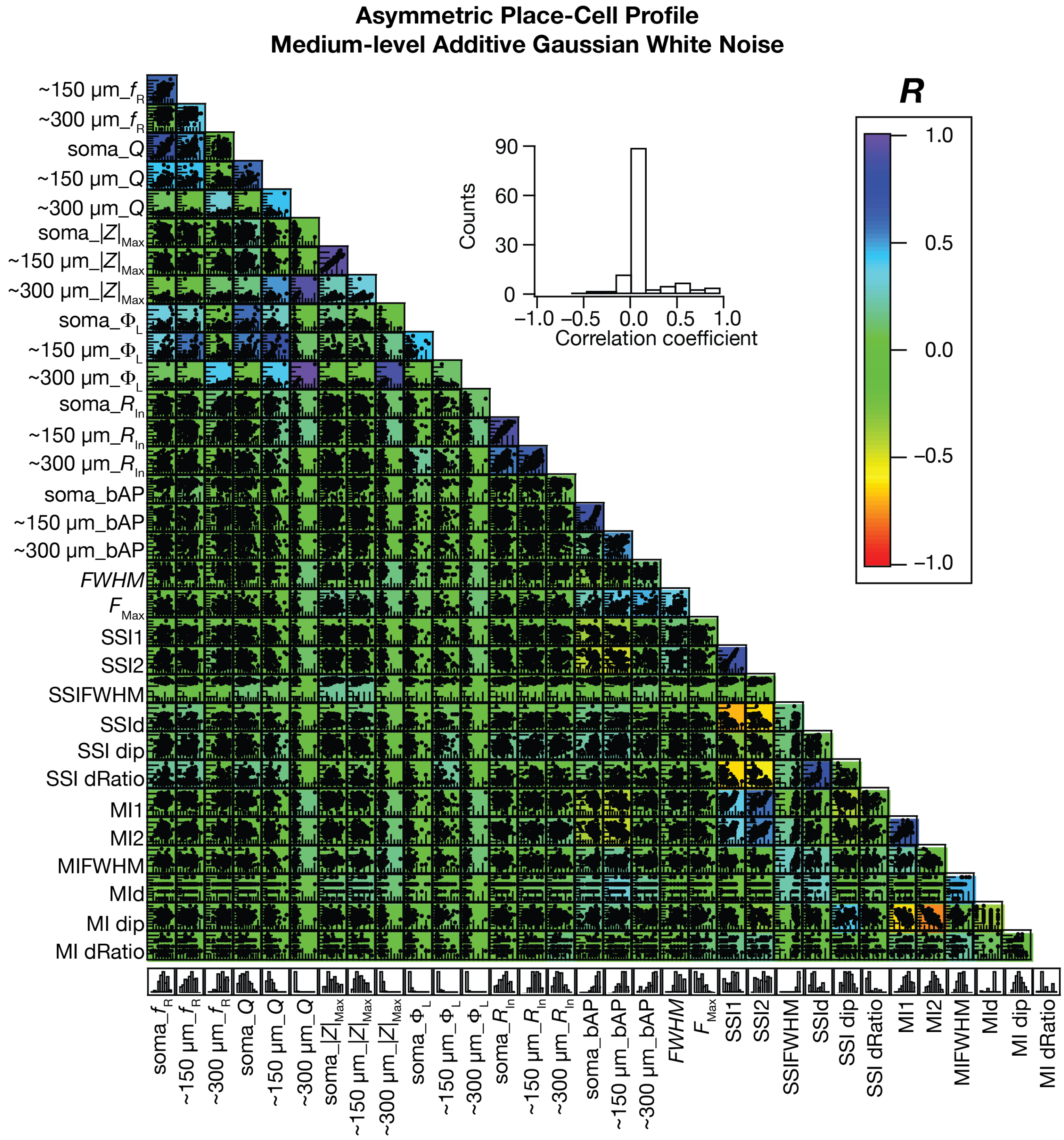

**Supplementary Figure S9. Impact of medium-level additive Gaussian white noise on pairwise correlations between intrinsic measurements, place-cell firing properties and information measurements, in models with asymmetric place-field firing.** Pairwise scatter plot matrix of intrinsic physiological measures, place cell firing properties and information measures (defined in Fig. 7) superimposed on the corresponding correlation coefficient matrix for the 127 valid models. Inset shows the histogram of all the correlation coefficient values.

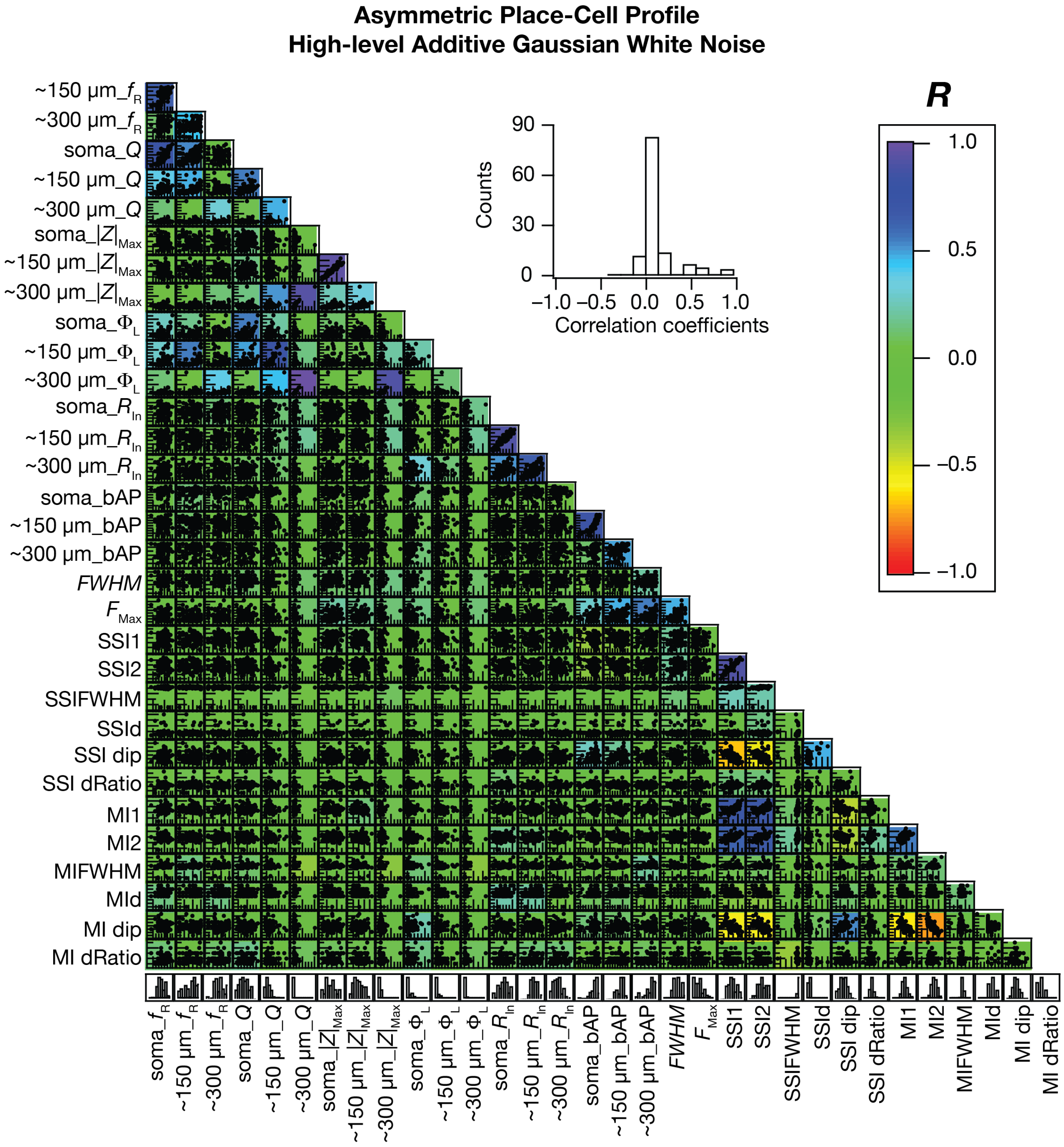

**Supplementary Figure S10. Impact of high-level additive Gaussian white noise on pairwise correlations between intrinsic measurements, place-cell firing properties and information measurements, in models with asymmetric place-field firing.** Pairwise scatter plot matrix of intrinsic physiological measures, place cell firing properties and information measures (defined in Fig. 7) superimposed on the corresponding correlation coefficient matrix for the 127 valid models. Inset shows the histogram of all the correlation coefficient values.

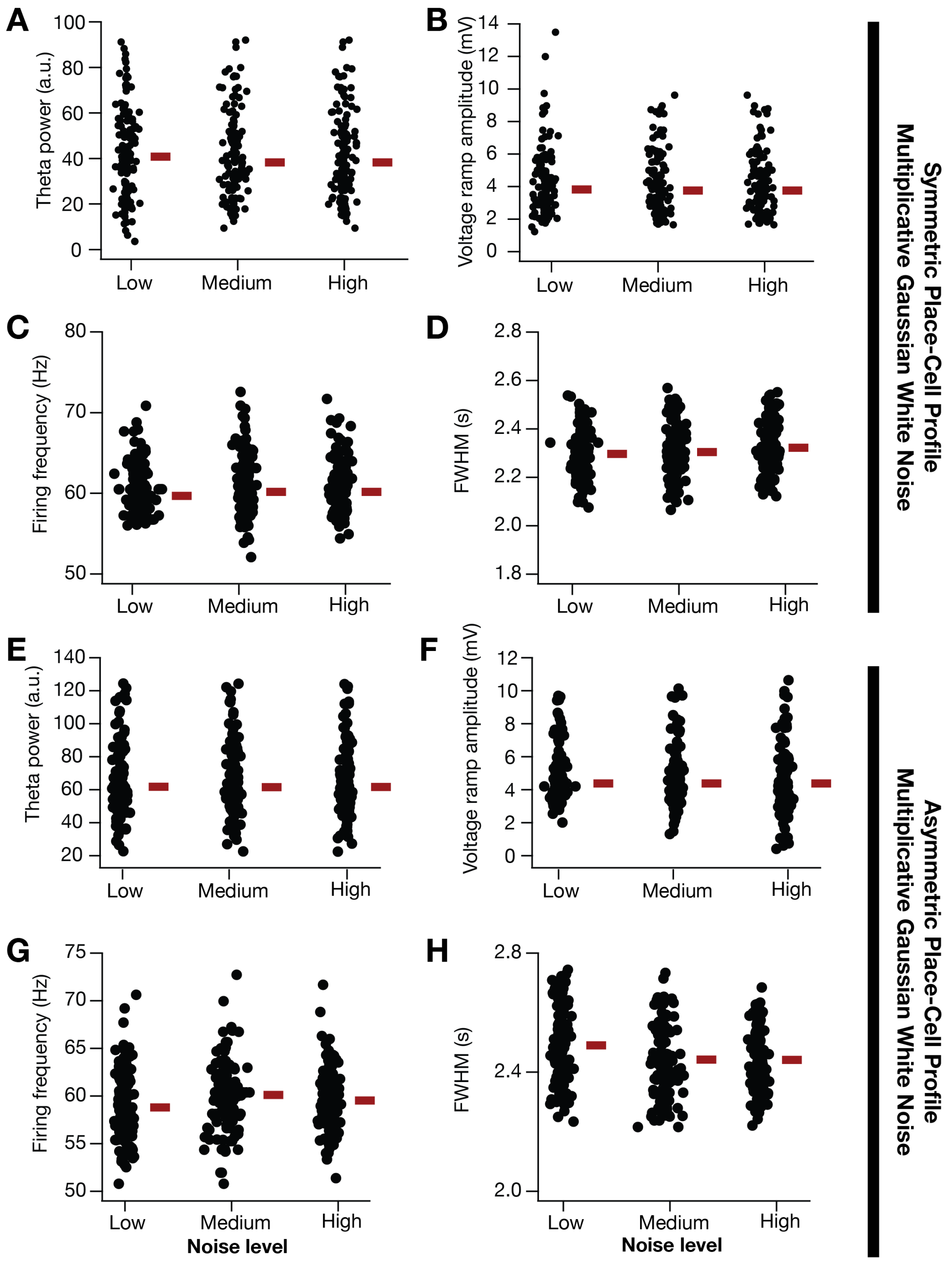

**Supplementary Figure S11. Impact of multiplicative Gaussian white noise (MGWN) on models with symmetric and asymmetric place-field firing profiles.** (A–D) Impact of different levels of MGWN on theta power (A), voltage ramp amplitude (B), peak firing frequency, $F_{max}$ (C) and full-width at half maximum, *FWHM* (D) of the 127 valid models with symmetric place-field firing profiles. The red bars represent the respective median values. (E–H) Same as (A–D), for asymmetric place-field profiles. MGWN variance values: *Low*: 0.01 Hz$^2$, *Medium*: 0.1 Hz$^2$, *High*: 0.5 Hz$^2$.

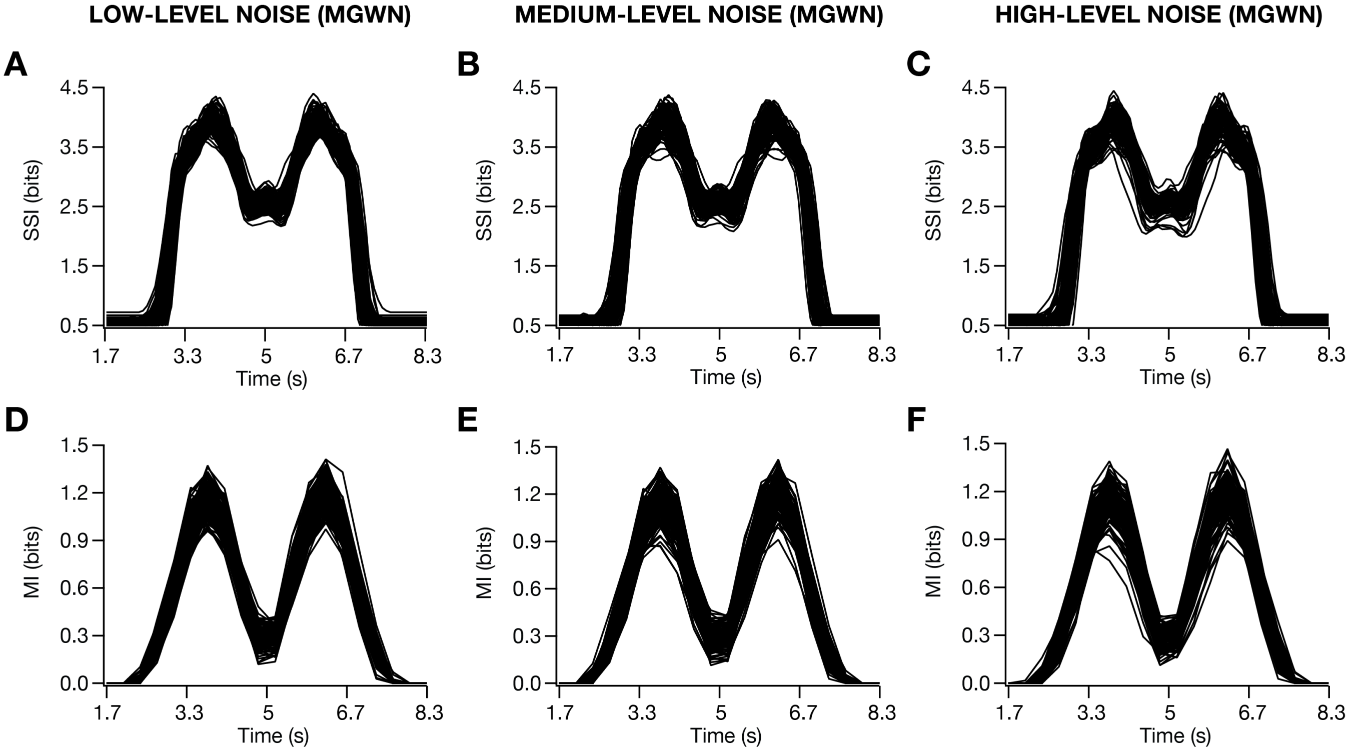

**Supplementary Figure S12. Impact of enhanced trial-to-trial variability, imposed as a multiplicative Gaussian white noise (MGWN), on spatial information transfer in place-cell models with symmetric place-field firing.** Stimulus specific information (SSI) profiles (A–C) and mutual information profiles (D–F) as functions of time, shown for low (plots on the left), medium (plots in the middle), high (plots on the right) levels of MGWN. MGWN variance values: *Low*: 0.01 $Hz^2$, *Medium*: 0.1 $Hz^2$, *High*: 0.5 $Hz^2$.

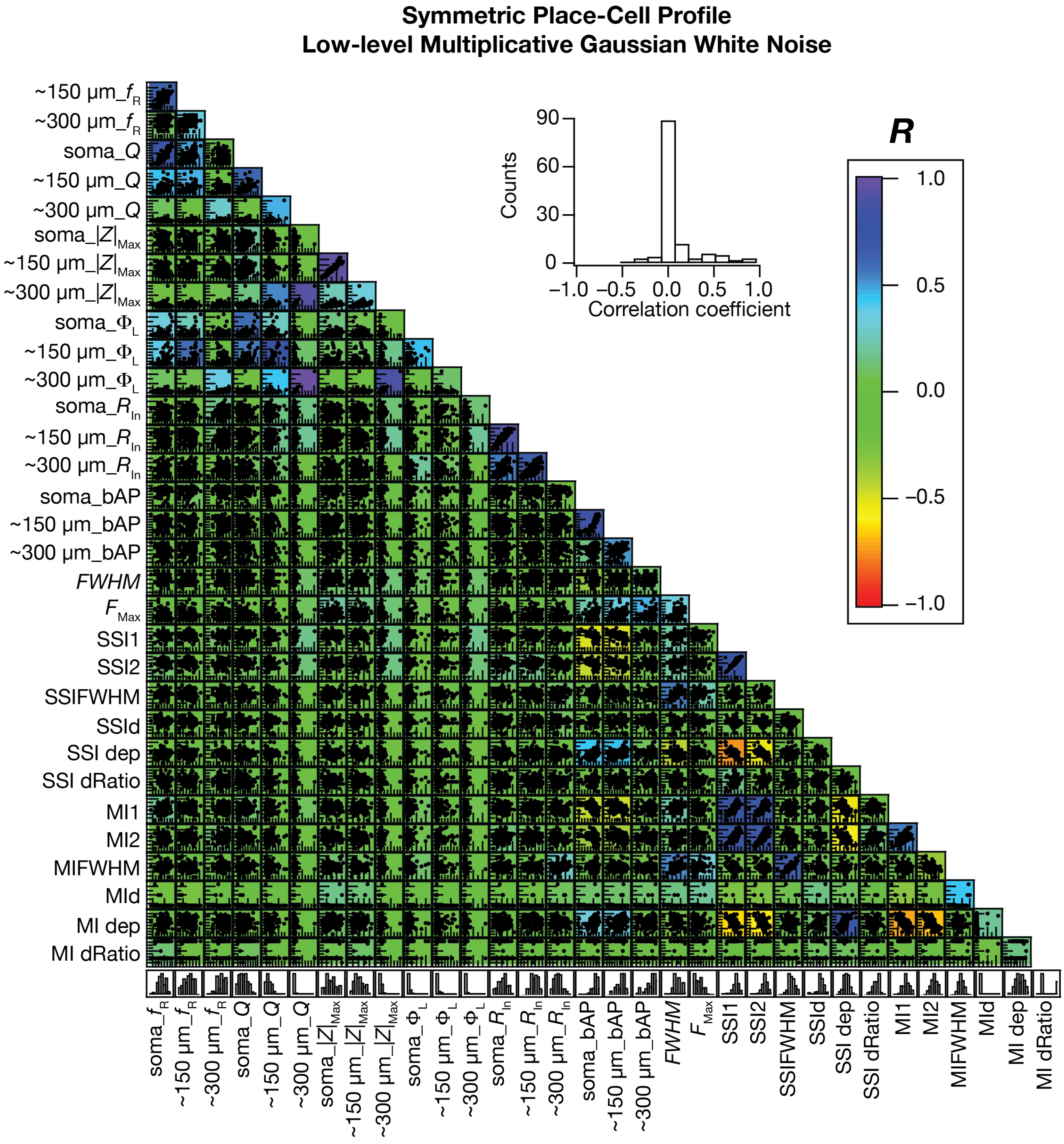

**Supplementary Figure S13. Impact of low-level multiplicative Gaussian white noise on pairwise correlations between intrinsic measurements, place-cell firing properties and information measurements.** Pairwise scatter plot matrix of intrinsic physiological measures, place cell firing properties and information measures (defined in Fig. 7) superimposed on the corresponding correlation coefficient matrix for the 127 valid models. Inset shows the histogram of all the correlation coefficient values.

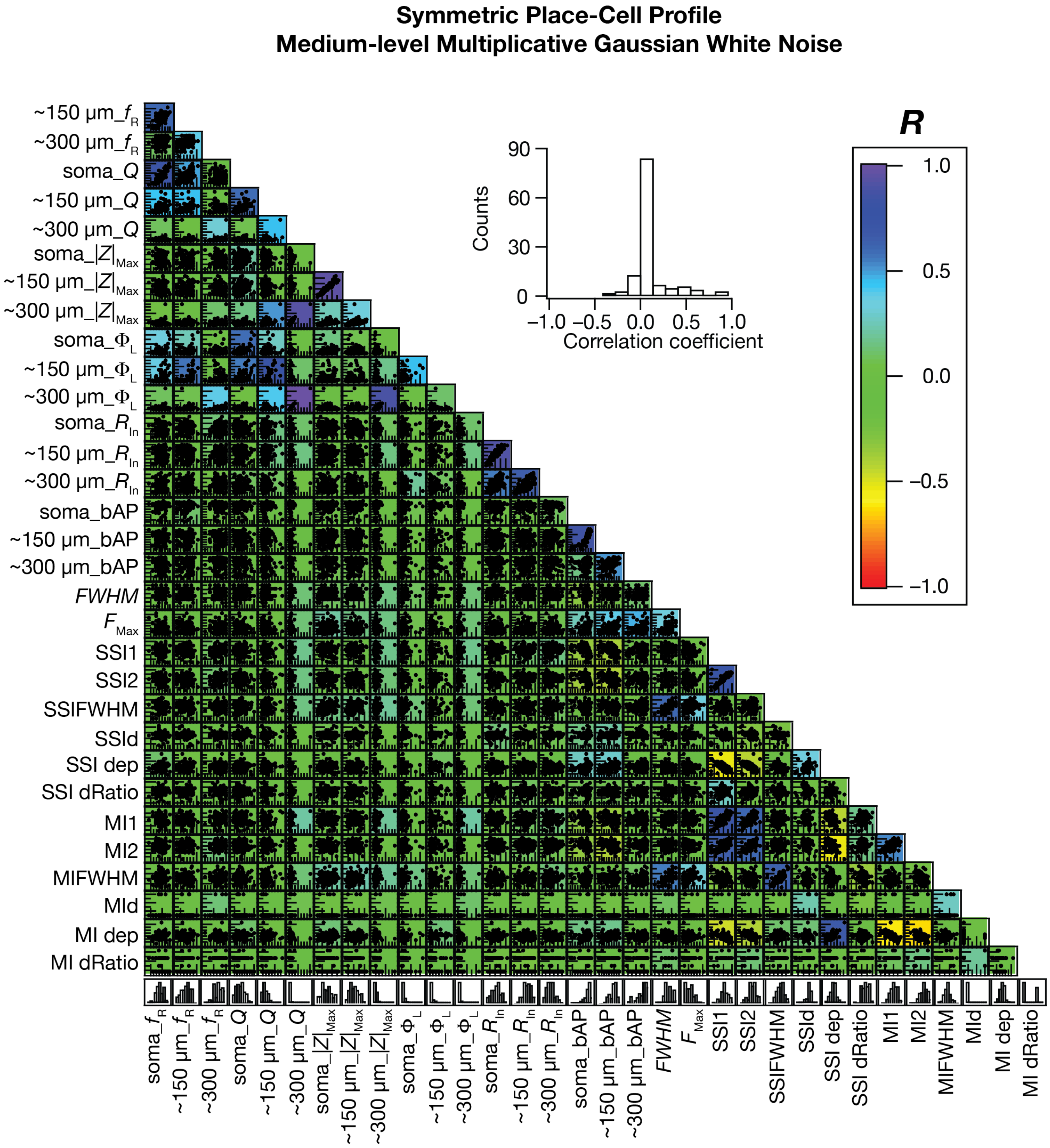

**Supplementary Figure S14. Impact of medium-level multiplicative Gaussian white noise on pairwise correlations between intrinsic measurements, place-cell firing properties and information measurements.** Pairwise scatter plot matrix of intrinsic physiological measures, place cell firing properties and information measures (defined in Fig. 7) superimposed on the corresponding correlation coefficient matrix for the 127 valid models. Inset shows the histogram of all the correlation coefficient values.

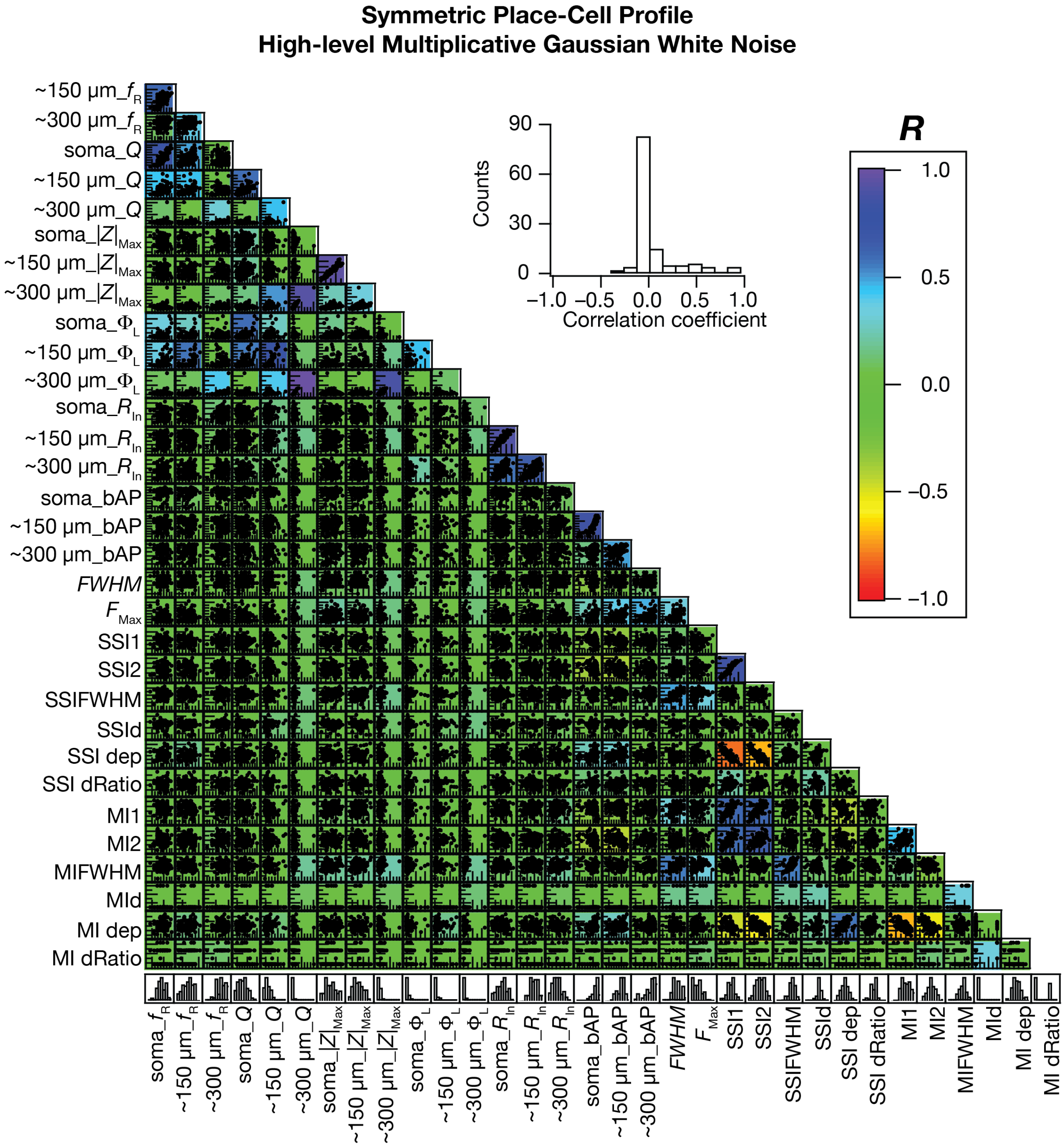

**Supplementary Figure S15. Impact of high-level multiplicative Gaussian white noise on pairwise correlations between intrinsic measurements, place-cell firing properties and information measurements.** Pairwise scatter plot matrix of intrinsic physiological measures, place cell firing properties and information measures (defined in Fig. 7) superimposed on the corresponding correlation coefficient matrix for the 127 valid models. Inset shows the histogram of all the correlation coefficient values.

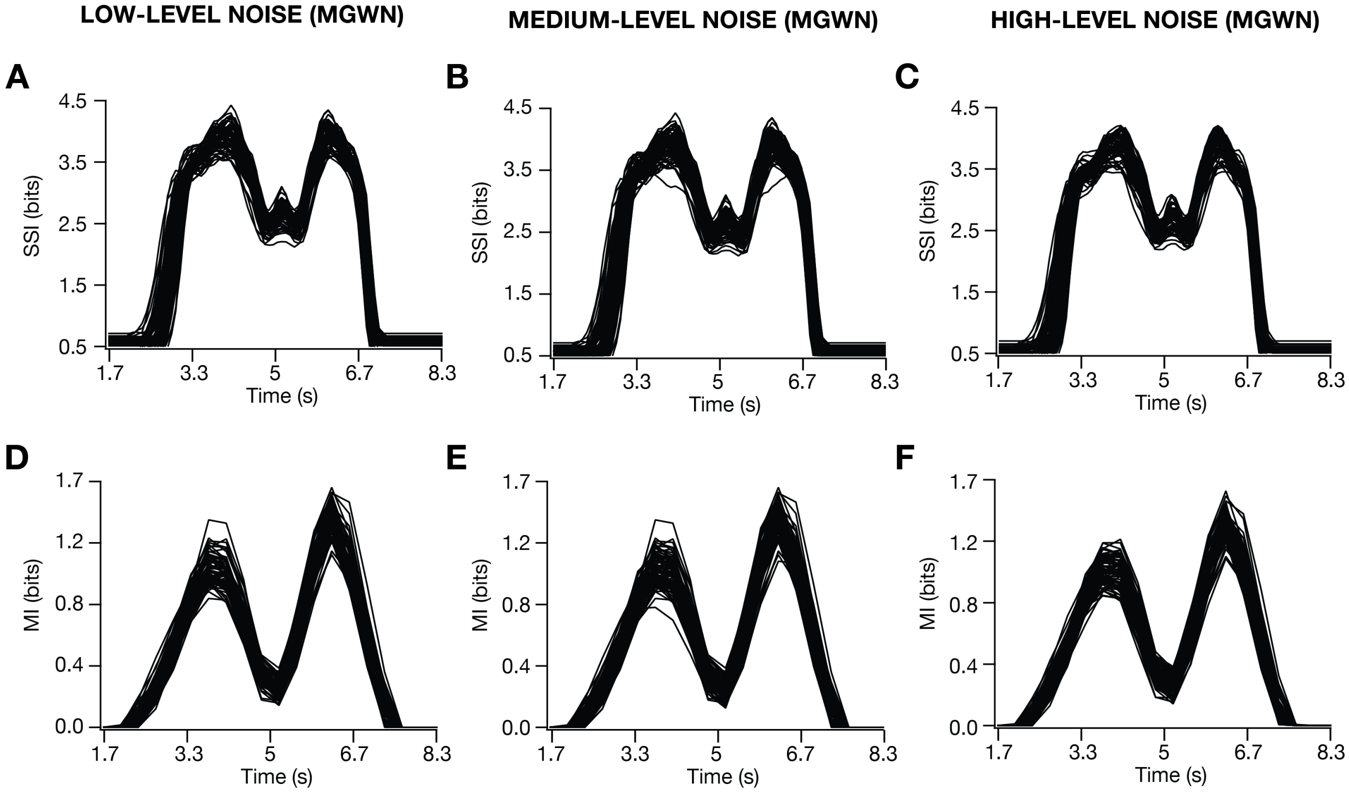

**Supplementary Figure S16. Impact of enhanced trial-to-trial variability, imposed as a multiplicative Gaussian white noise (MGWN), on spatial information transfer in place-cell models with asymmetric place-field firing.** Stimulus specific information (SSI) profiles (A–C) and mutual information profiles (D–F) as functions of time, shown for low (plots on the left), medium (plots in the middle), high (plots on the right) levels of MGWN. MGWN variance values: *Low*: 0.01 $Hz^2$, *Medium*: 0.1 $Hz^2$, *High*: 0.5 $Hz^2$.

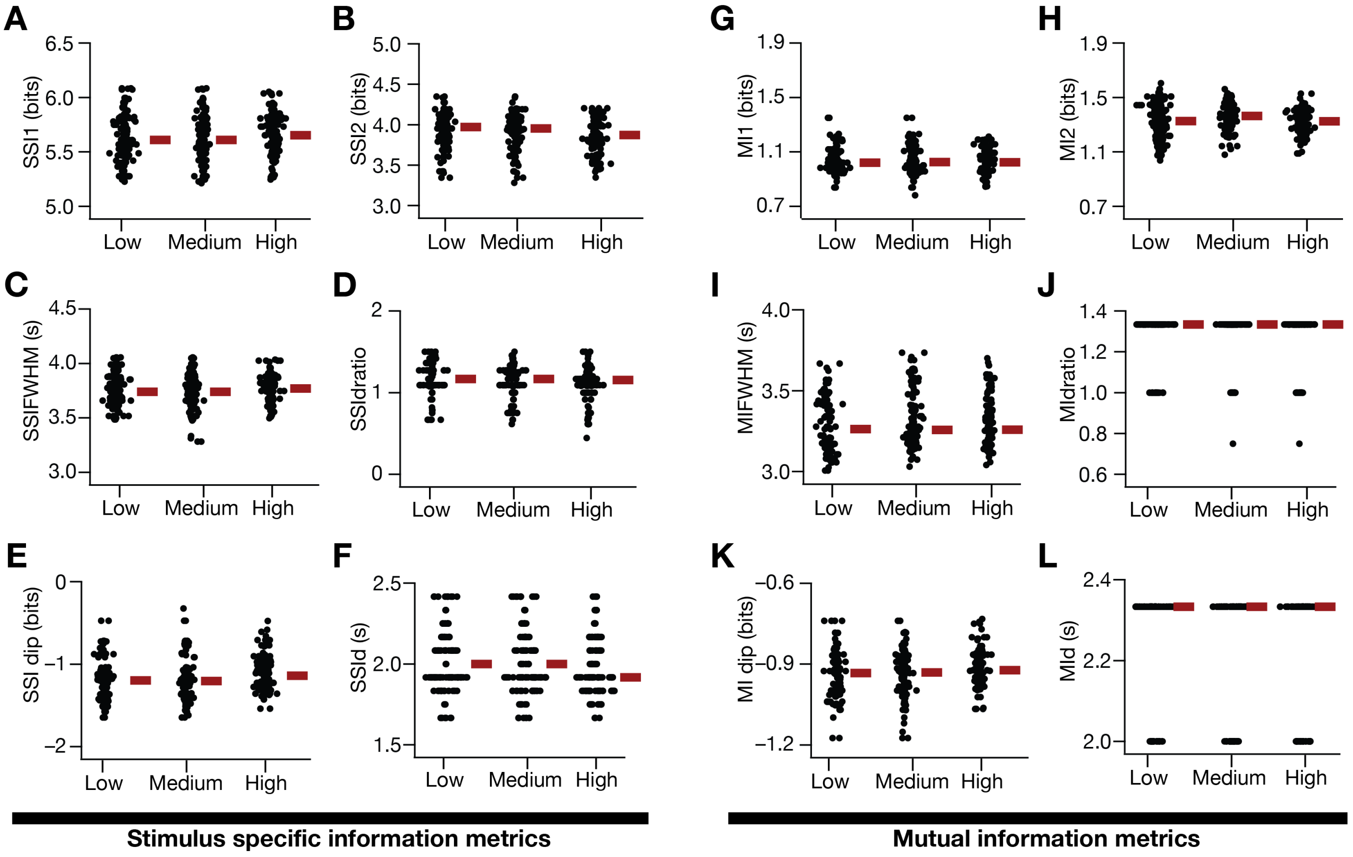

**Supplementary Figure S17. Quantification of spatial information transfer as a function of enhanced trial-to-trial variability, imposed as a multiplicative Gaussian white noise (MGWN) in models with asymmetric place-field firing.** (A–F) SSI metrics for the population of valid models depicting the impact of three levels of noise on the first (B, *SSI1*) and second (C, *SSI2*) peaks of SSI, the full width half maximum of the SSI profile (D, *SSIFWHM*), the ratio of the first peak-to-center distance to the center-to-second peak distance (E, *SSI dRatio*), the difference between the SSI value at the place field center to the peak SSI value (F, *SSI dip*) and the difference between the location of *SSI1* and *SSI2* (G, *SSI d*). (G–L) Same as (A–F) for mutual information profiles of the valid model population. MGWN variance values: *Low*: 0.01 Hz$^2$, *Medium*: 0.1 Hz$^2$, *High*: 0.5 Hz$^2$.

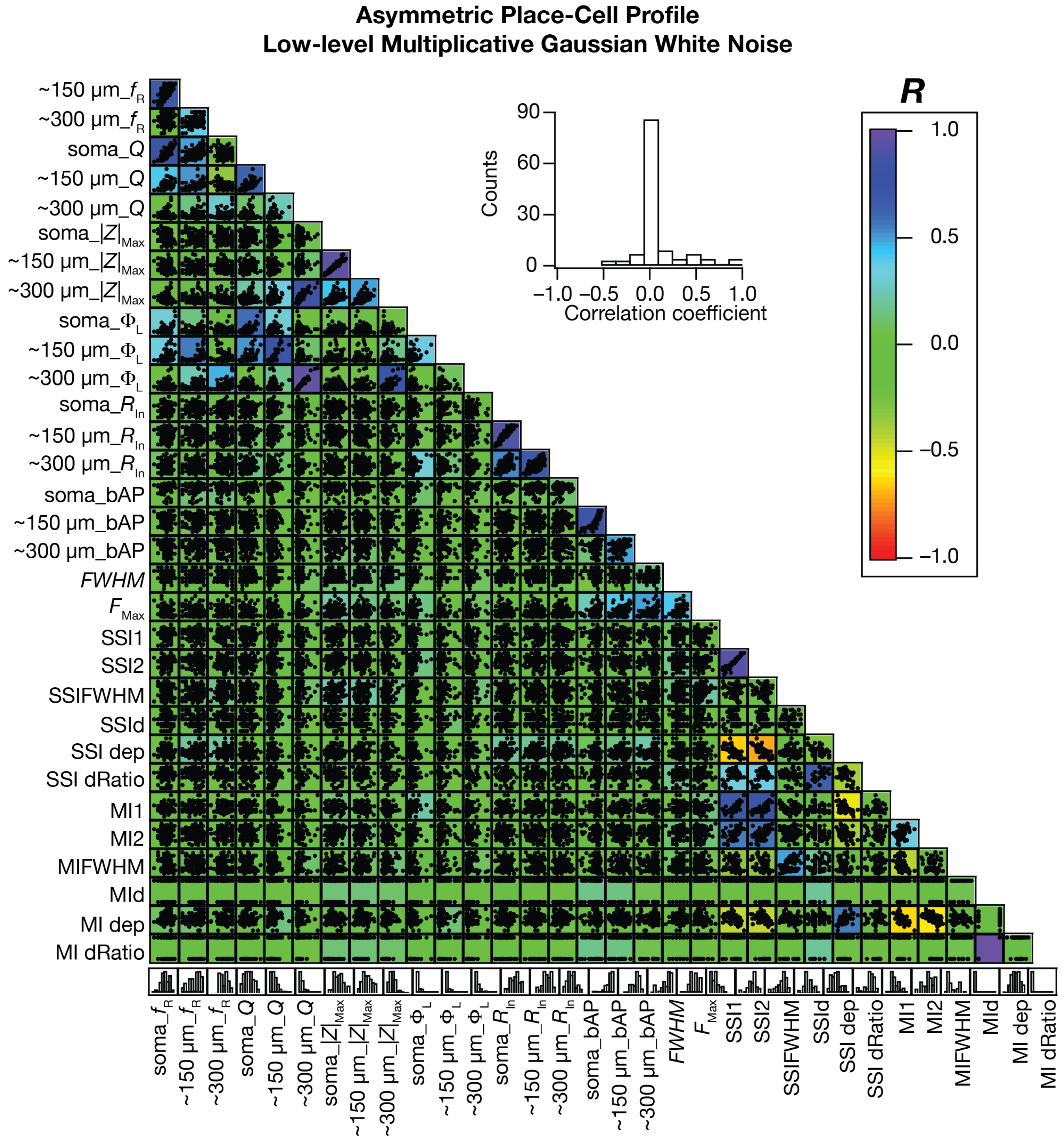

**Supplementary Figure S18. Impact of low-level multiplicative Gaussian white noise on pairwise correlations between intrinsic measurements, place-cell firing properties and information measurements, in models with asymmetric place-field firing.** Pairwise scatter plot matrix of intrinsic physiological measures, place cell firing properties and information measures (defined in Fig. 7) superimposed on the corresponding correlation coefficient matrix for the 127 valid models. Inset shows the histogram of all the correlation coefficient values.

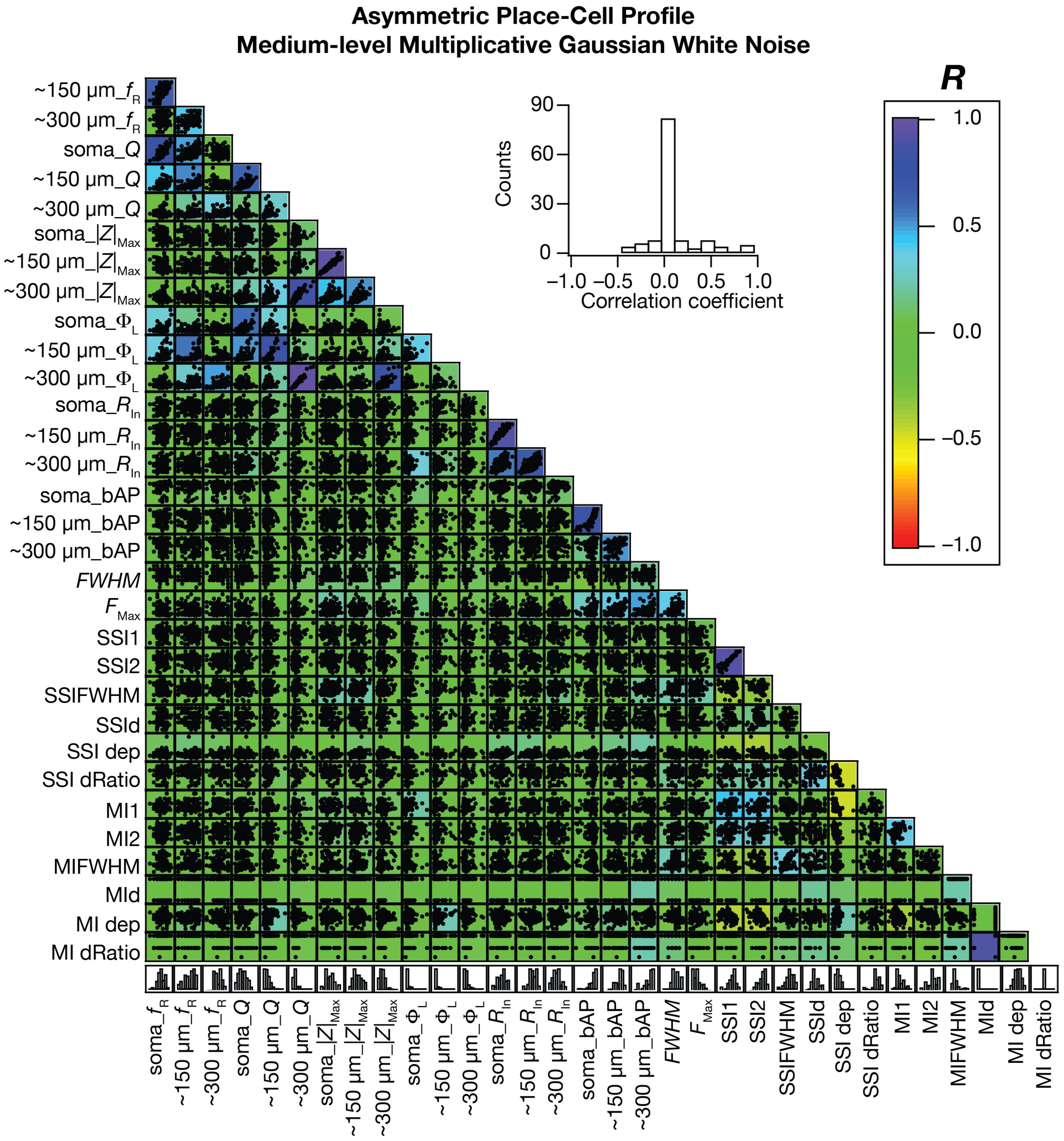

**Supplementary Figure S19. Impact of medium-level multiplicative Gaussian white noise on pairwise correlations between intrinsic measurements, place-cell firing properties and information measurements, in models with asymmetric place-field firing.** Pairwise scatter plot matrix of intrinsic physiological measures, place cell firing properties and information measures (defined in Fig. 7) superimposed on the corresponding correlation coefficient matrix for the 127 valid models. Inset shows the histogram of all the correlation coefficient values.

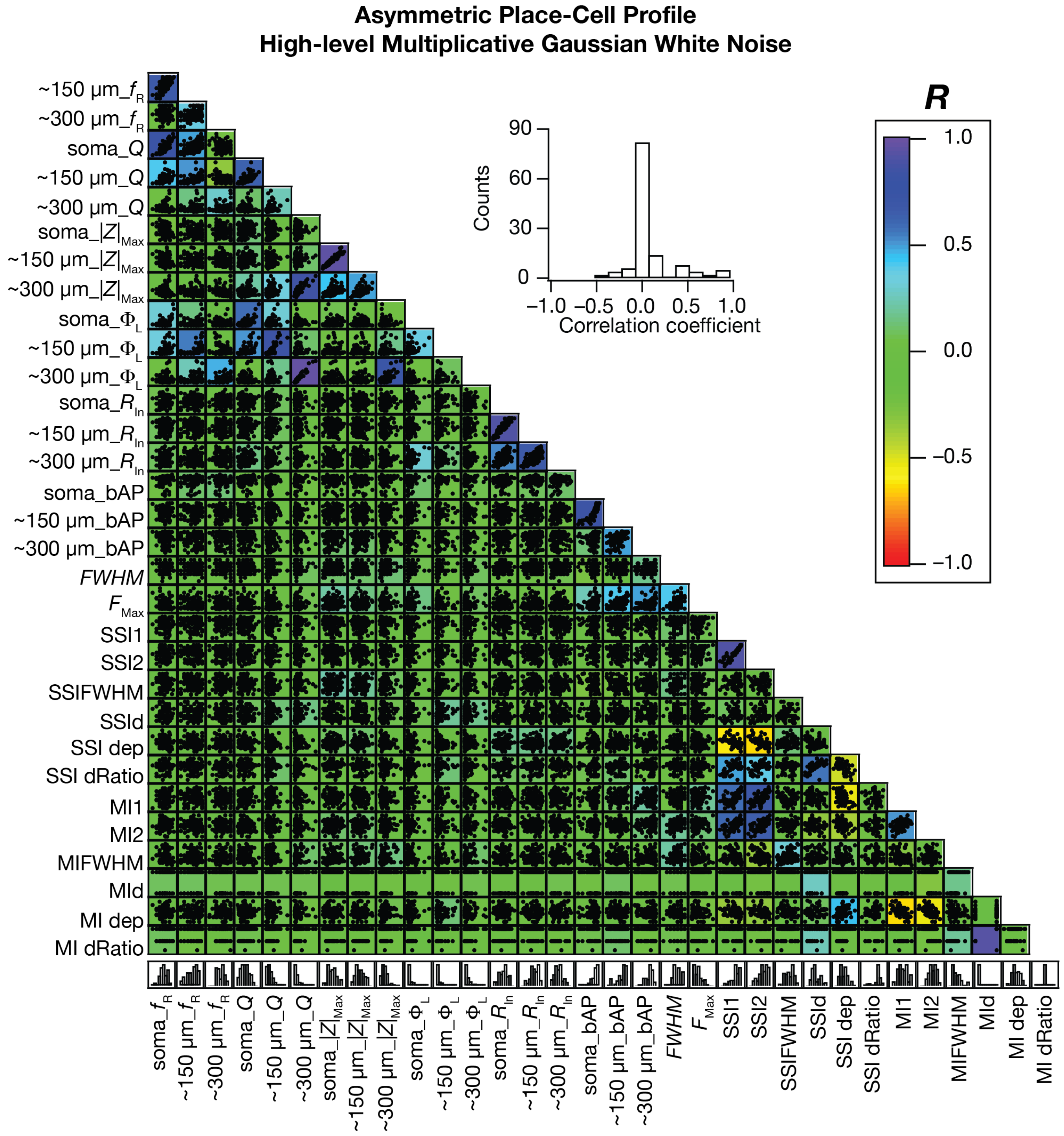

**Supplementary Figure S20. Impact of high-level multiplicative Gaussian white noise on pairwise correlations between intrinsic measurements, place-cell firing properties and information measurements, in models with asymmetric place-field firing.** Pairwise scatter plot matrix of intrinsic physiological measures, place cell firing properties and information measures (defined in Fig. 7) superimposed on the corresponding correlation coefficient matrix for the 127 valid models. Inset shows the histogram of all the correlation coefficient values.

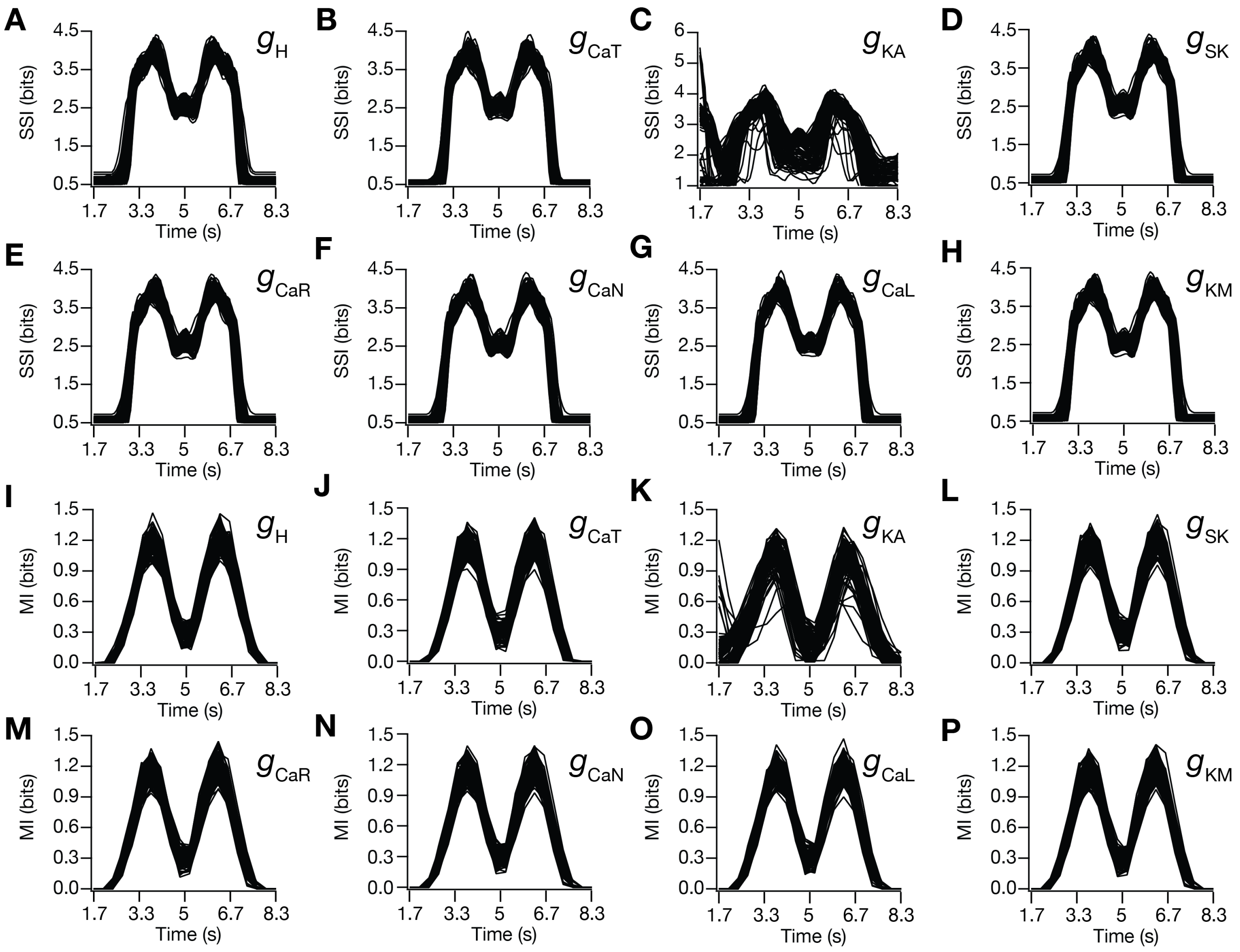

**Supplementary Figure 21. Impact of virtual knockout of ion channels on the information profiles in the model population.** (A–P) Impact of virtually knocking out individual channels on the population of models in terms of SSI profiles (A–H) and MI profiles (I–P). Shown are profiles obtained after setting each specific conductance to zero in each model, with the rest of the model remaining intact. $g_{\mathrm{H}}$: *h* channel; $g_{\mathrm{CaT}}$: *T*-type calcium channel; $g_{\mathrm{KA}}$: *A*-type potassium channel; $g_{\mathrm{SK}}$: small conductance calcium-activated potassium channel; $g_{\mathrm{CaR}}$: *R*-type calcium channel; $g_{\mathrm{CaN}}$: *N*-type calcium channel; $g_{\mathrm{CaL}}$:, *L*-type calcium channel; $g_{\mathrm{KM}}$: *M*-type potassium channel.

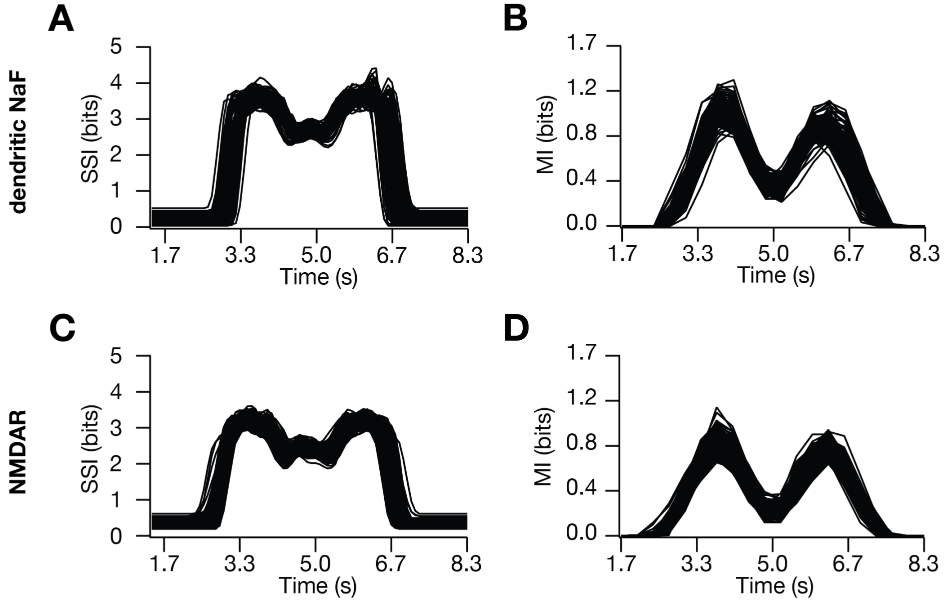

**Supplementary Figure 22. Impact of removal of dendritic sodium channels or NMDA receptors on the information profiles in the model popultation.** (A–B) Impact of virtually knocking out dendritic NaF channels on the population of models in terms of SSI (A) and MI (B) profiles. Shown are profiles obtained after setting apical dendritic fast sodium conductance to zero in each model, with the rest of the model remaining intact. (C–D) Impact of virtually knocking out NMDARs on the population of models in terms of SSI (C) and MI (D) profiles. Shown are profiles obtained after setting the value of NMDAR permeability to zero for all synapses in each model, with the rest of the model remaining intact.

**Supplementary Table S1:** The SSI and MI measurements for the five similar models shown in Fig. S2.

| Information Measures | 1 | 2 | 3 | 4 | 5 |
|---|---|---|---|---|---|
| **SSI1** | 3.80017 | 3.77615 | 3.70755 | 4.08849 | 4.14399 |
| **SSI2** | 3.77416 | 3.77557 | 3.68526 | 4.10592 | 4.06252 |
| **SSIFWHM** | 3.85822 | 3.86936 | 3.48788 | 3.559773 | 3.919847 |
| **SSIdRatio** | 1.15385 | 1 | 0.6875 | 1.15385 | 1.07692 |
| **SSIdip** | -1.21945 | -1.01558 | -1.5041 | -1.37414 | -1.49871 |
| **SSId** | 2.333333 | 2.333333 | 2.25 | 2.333333 | 2.25 |
| **MI1** | 1.13212 | 1.11905 | 1.12318 | 1.26352 | 1.15849 |
| **MI2** | 1.17915 | 1.10346 | 1.04006 | 1.29898 | 1.2747 |
| **MIFWHM** | 3.6514 | 3.70838 | 3.895893 | 3.403 | 3.71434 |
| **MIdRatio** | 1.33333 | 0.75 | 0.75 | 1.33333 | 0.75 |
| **MIdip** | -0.87061 | -0.83072 | -0.96913 | -0.94316 | -0.99808 |
| **Mid** | 2.333333 | 2.333333 | 2.333333 | 2.333333 | 2.333333 |
